# Structural variation and eQTL analysis in two experimental populations of chickens divergently selected for feather pecking behavior

**DOI:** 10.1101/2022.09.16.508210

**Authors:** Clemens Falker-Gieske, Jörn Bennewitz, Jens Tetens

## Abstract

Feather pecking (FP) is a damaging non-aggressive behavior in laying hens with a heritable component. Its occurrence has been linked to the immune system, the circadian clock, and foraging behavior. Furthermore, dysregulation of miRNA biogenesis, disturbance of the gamma-aminobutyric acid (GABAergic) system, as well as neurodevelopmental deficiencies are currently under debate as factors influencing the propensity for FP behavior. Past studies, which focused on the dissection of the genetic factors involved in FP, relied on single nucleotide polymorphisms (SNPs) and short insertions and deletions < 50 bp (InDels). These variant classes only represent a certain fraction of the genetic variation of an organism. Hence, we re-analyzed whole genome sequencing data from two experimental populations, which have been divergently selected for FP behavior for over more than 15 generations, and performed variant calling for structural variants (SVs) as well as tandem repeats (TRs) and jointly analyzed the data with SNPs and InDels. Genotype imputation and subsequent genome-wide association studies in combination with expression quantitative trait loci analysis led to the discovery of multiple variants influencing the GABAergic system. These include a significantly associated TR downstream of the GABA receptor subunit beta-3 (*GABRB3*) gene, two micro RNAs targeting several GABA receptor genes, and dystrophin (*DMD*), a direct regulator of GABA receptor clustering. Furthermore, we found the transcription factor *ETV1* to be associated with differential expression of 23 genes, which points towards a role of *ETV1*, together with *SMAD4* and *KLF14*, in the disturbed neurodevelopment of high feather pecking chickens.

## Introduction

Feather pecking (FP) in chickens is a behavioral disorder, which severely impacts animal welfare and causes significant economic losses. It has been proposed that FP is obsessive compulsive-like behavior. [3]. In the past, the damage has been controlled by beak trimming, which has now been prohibited in many European countries. Numerous studies found an involvement of environmental factors such as light intensity, nutrition, stocking density, and lack of foraging (reviewed by [4] and [5]). Furthermore, evidence has been accumulating that the immune system plays a major role in the development of FP behavior [1, 6–8]. The gut microbiota have also been proposed to be involved in FP behavior but this has been disproven in several studies [9, 10]. Since feather pecking is a complex heritable trait (reviewed by [11]) the dissection of the genetic causes is essential for the development of effective breeding strategies to eradicate the causative alleles. To achieve that goal chickens were divergently selected for FP behavior over more than 15 generations based on their estimated breeding values for the behavior. Breeding of these lines was initiated in Denmark and continued in Hohenheim, Germany. In Hohenheim two populations were established – an F_2_ cross and 12 half-sib families (in the preceding text referred to as F_2_ and HS). A detailed description of the experimental populations and the research conducted with them was reviewed by Bennewitz and Tetens [12]. Based on the results that were acquired by whole-genome and transcriptome sequencing with the Hohenheim selection lines, FP appears to be a disorder of the γ-aminobutyric acid (GABA) system in conjunction with a disturbance of embryonic neurodevelopment by a lack of leukocytes in the developing brain. Several variants in or in close proximity to GABA receptor genes were identified in genome-wide association studies (GWAS) conducted on medium density single nucleotide polymorphism (SNP) array and imputed sequence level genotypes [2, 13, 14]. Furthermore, brain transcriptome analysis of high and low feather peckers (HFP and LFP) before and after light stimulation revealed that HFP respond with very few changes in gene expression in comparison to LFP with numerous GABA receptor genes upregulated in LFP. Only *GABRB2* (Gamma-Aminobutyric Acid Type A Receptor Subunit Beta2) was among differentially expressed genes (DEGs) in HFP but down-instead of upregulated – a pattern that we observed for most DEGs in HFP brains [14]. We attribute this low level of gene expression changes in response to light to a high level of excitation in HFP brains due to the lack of multiple GABA receptors. Since GABA is the major inhibitory neurotransmitter a lack of its receptors in the brain would lead to a high neuronal excitatory state, which explains the observed hyperactivity and the obsessive compulsive-like behavior observed in HFP. Since *Dicer1* was among the downregulated DEGs we assume that miRNA processing is disturbed, as it is also the case in schizophrenia patients [15], which in turn leads to low GABA receptor expression levels. In the general comparison of brain transcriptomes between HFP and LFP hens, we observed an enrichment of immune system-related DEGs [1], which we could further pinpoint in an expression quantitative trait loci (eQTL) analysis to a small deletion 652 bp downstream of the *KLF14* gene. In total, the differential expression of 40 genes between HFP and LFP chickens was significantly associated with this *KLF14* variant, a majority of which are involved in leukocyte biology [8]. It has been shown in mice that CD4 T cells are essential for healthy development from the fetal to the adult brain. A defect in CD4 T cell maturation affected on synapse development and lead to behavioral abnormalities [16]. The evidence suggests that this mechanism is responsible for disturbed embryonic brain development in HFP chickens, which contributes to feather pecking behavior.

One commonality of all the studies that we conducted on the genetics involved in FP is the overlap in associated genes with human psychiatric disorders, most prevalent schizophrenia. Structural genomic variants (SVs) play a notable role in human psychiatric disorders [17], which is also the case for tandem repeats (TRs) [18]. Commonly investigated classes of SVs include insertions, deletions, duplications, inversions, and translocations, which arise from various combinations of DNA gains, losses, or rearrangements [19]. Here we present the first in-depth study on the potential role of SVs and TRs in FP, which led to a deeper understanding of the mechanisms responsible for this behavioral disorder.

## Material and methods

### Animals and husbandry

All experimental procedures [1], rearing and husbandry conditions [20] as well as phenotyping [2] were described in previous studies. Briefly, the phenotypic data were generated by direct observations made by seven independent investigators. The phenotypes are expressed as the number of FP bouts actively delivered during a standardized time span. As these count data heavily distributed from normality, Box-Cox-transformation was applied [2, 13].

### Structural variants and short tandem repeats discovery

Illumina whole genome sequencing data were mapped to chicken genome version GRCg6a (GCF_000002315.5 RefSeq assembly) and used to call SNPs and short (< 50bp) insertions and deletions (InDels) in our previous study [2]. SVs were called as described by Blaj *et al*. [21] with slightly different settings: A high confidence SV call set was produced from the output of three variant callers: smove v0.2.6 (Brent, P. (2018). Smoove. https://brentp.github.io/post/smoove/), DELLY v0.7.7 [22], and manta v1.6.0 [23]. SURVIROR v1.0.7 [24] was used to combine the output of the three variant callers with the following settings: maximum distance between breakpoints of 1000 bp, minimum number of supporting callers: 2, SV type and strands were taken into account, and the minimum SV size was set to 30 bp. Variants with a call-rate < 0.8 and variants with QUAL < 1000 were removed. Tandem repeats were called with GangSTR [25]. A library of known TRs as input for GangSTR was acquired from the UCSC data repository (https://hgdownload.soe.ucsc.edu/goldenPath/galGal6/bigZips/). TRs were filtered to retain genotypes with a minimum sequence depth (DP) of 10, a quality score (Q) higher than 0.8, and a call-rate of 0.8.

### Haplotype construction and imputation

To impute medium density chip genotypes to whole genome level SNPs and InDels we employed the same strategy as we previously described [2] with the deviation that all imputation steps were performed with Beagle 5.2 [26] and the setting ne=1000. SVs and TRs were merged with SNPs and InDels from our previous study, which were acquired with the genome analysis toolkit (GATK) [27]. This merged call set was phased with Beagle 5.2 for the estimation of haplotypes and used as a reference panel to impute medium density chip genotypes to SVs and TRs with the same strategy as for SNPs and InDels. To remove SNPs and InDels from the imputed SVs/TRs GATK v4.1.8.1 SelectVariants was utilized.

### Detection of quantitative trait loci

Prior to GWAS multiallelic variants from the SVs/TR dataset were converted to biallelic variants with the norm function from bcftools (v. 1.14) [28]. All GWAS were conducted with gcta v. 1.92.3 beta3 [29] applying a mixed linear model association analysis with a leaving-one-chromosome-out approach. The phenotype used for GWAS, “*Feather Pecks Delivered Box-Cox Transformed*” (FPD_BC), has been described by Iffland et al. [13]. Phenotypes for expression GWAS (eGWAS) were normalized gene expression data from our previous study [8]. Information on the hatch was used as a covariate in all GWAS and eGWAS. Genomic relationship matrices were created from the target 60k SNP Chip genotypes. Meta-analyses of GWAS results were performed with METAL [30] as previously described [2]. The proportion of variance in phenotype explained by a given SNP (PVE) was calculated according to Shim *et al*. [31].

### Association weight matrix construction

The AWM was created as described in our previous study [8] by deploying the strategy for AWM construction by Reverter and Fortes [32] followed by the detection of significant gene-gene interactions with their PCIT algorithm [33]. Input variants were chosen as follows: *P* value < 1 × 10^−4^ for the main phenotype (FPD_BC) or *P* value < 1 × 10^−4^ in least ten of the eGWAS. That way, variants affecting the main phenotype and gene expression were, were considered in the analysis. This led to the selection of 57 input variants, 0.16% of all detected SVs and TRs for the HS population. The R script by Reverter and Fortes was modified by setting the *P* value threshold for primary and secondary SNP selection to 1 × 10^−4^. The gene-gene interaction map was constructed with Cytoscape and gene class information was acquired with PANTHER [34] and UniProt [35].

### Transcription factor enrichment

To detect significant binding site enrichment for the transcription factor *ETV1* the CiiiDER software (build May 15^th^, 2020) [36] was employed with frequency matrix MA0761.2 (https://jaspar.genereg.net/matrix/MA0761.2/). Additional members of the ETS (E twenty-six) family of transcription factors used in the analysis were *ETV2* (MA0762.1), *ETV3* (MA0763.1), *ETV4* (MA0764.2), *ETV5* (MA0765.1), *ETV6* (MA0645.1), and *ETV7* (MA1708.1). Genes that were associated with the *ETV1* variant (*P* value > 1 × 10^−4^; Supplementary Information S1) were used as input and DEGs with a Log_2_ fold change < 0.2 were used as background genes. The analysis was performed as in our previous study [8].

## Results

### Genome-wide association studies with different classes of genetic variants

In total 63,824 TRs and 11,098 SVs were discovered in the joint variant calling of whole genome sequenced HS and F_0_ chickens. The SVs contained 6,014 deletions, 2,741 inversions, 1,334 duplications, and 995 translocations. The variant calling of SNPs and InDels was reported in our previous study and yielded 12,864,421 SNPs and 2,142,539 InDels [2]. To grasp the whole depth of the datasets at hand we repeated the imputation of SNPs and InDels with the most recent Beagle version (v. 5.2) [26] and set the effective population size to 1000, which is known to improve the imputation accuracy in small populations [37]. GWAS results for the trait FPD_BC were analyzed for both experimental designs, the F_2_ cross and the HS population, separately and after combining the results in a metaanalysis. Manhattan plots from GWAS with imputed SNPs/InDels and imputed SVs/TRs are shown in Figure 1a. Common peaks for both variants classes on GGA1 and GGA2 were observable in the F_2_ cross, as well as on GGA1 and GGA3 of the HS population. QTL are summarized in Table 1 and lead SVs and TRs with their predicted effects are listed in Table 2.

**Figure 1:**
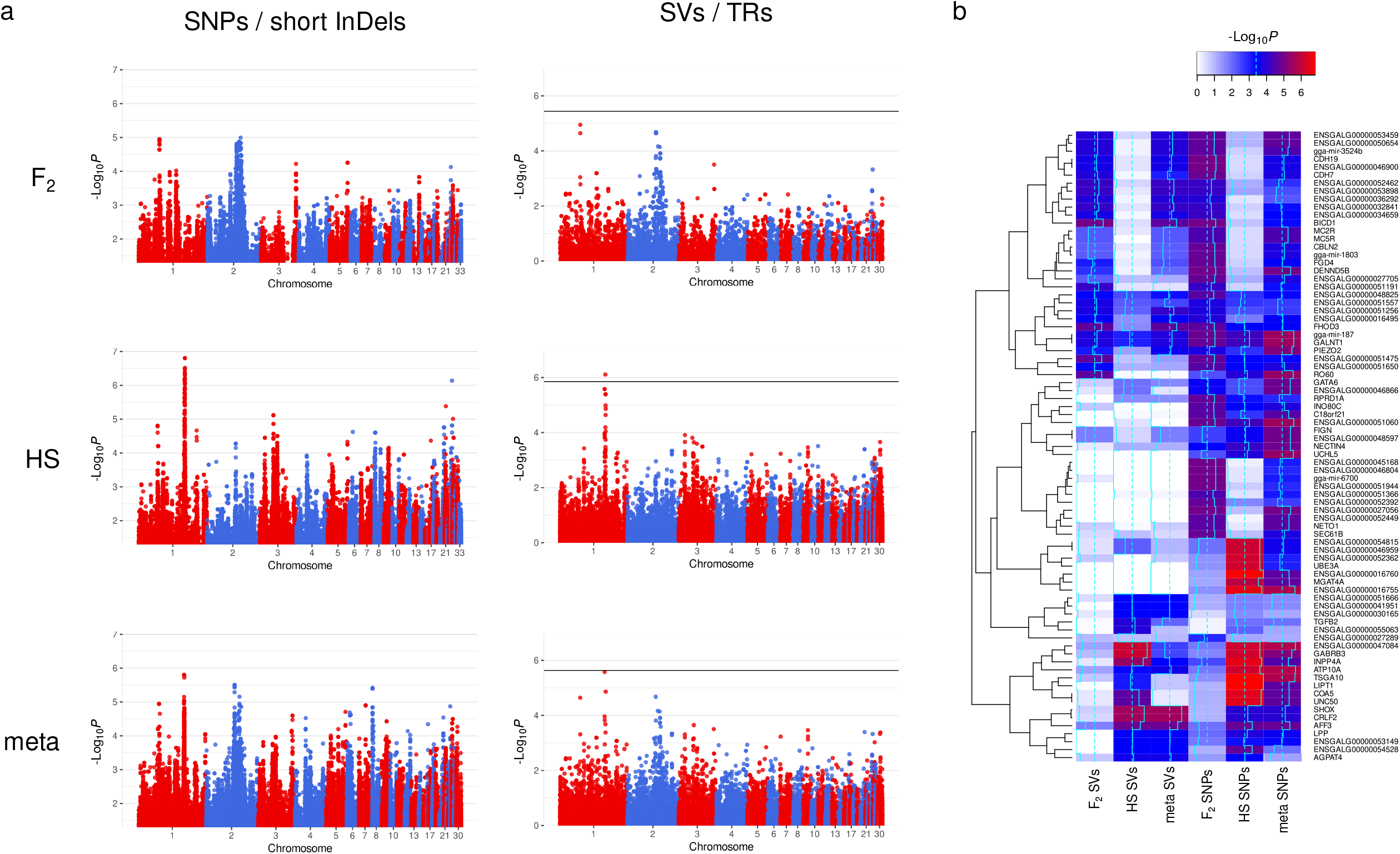
(a) Genome-wide association studies (GWAS) performed with single nucleotide polymorphisms (SNPs) in combination with short (< 50bp) insertions and deletions (InDels) and structural variants (SVs) in combination with tandem repeats (TRs). All variant classes were investigated in the F_2_ cross and half-sib (HS) families of chickens divergently selected for feather pecking and subsequently combined in a meta-analysis. (b) Heatmap of the 20 highest associated genes from each GWAS. The –Log_10_*P* values of the highest associated variants for each gene were used.

**Table 1:** Positional ranges in base pairs of QTL that were discovered in both experimental populations (F_2_ cross and half-sib (HS) families) for both groups of variant classes: Single nucleotide polymorphisms (SNPs) and short insertions/deletions (InDels) as well as structural variants (SVs) and tandem repeats (TRs).

| F<sub>2</sub> SNPs and short InDels | F<sub>2</sub> SVs and TRs | HS SNPs and short InDels | HS SVs and TRs |
| --- | --- | --- | --- |
| GGA1:59,032,772-59,548,358 | GGA1:59,138,261-59,216,858 | GGA1:132,578,705-133,803,185 | GGA1:130,621,356-134,077,752 |
| GGA2:81,060,432-97,072,903 | GGA2:83,493,152-94,489,210 | GGA3:18,434,015-54,601,143 | GGA3:19,120,538-70,385,553 |

**Table 2:**
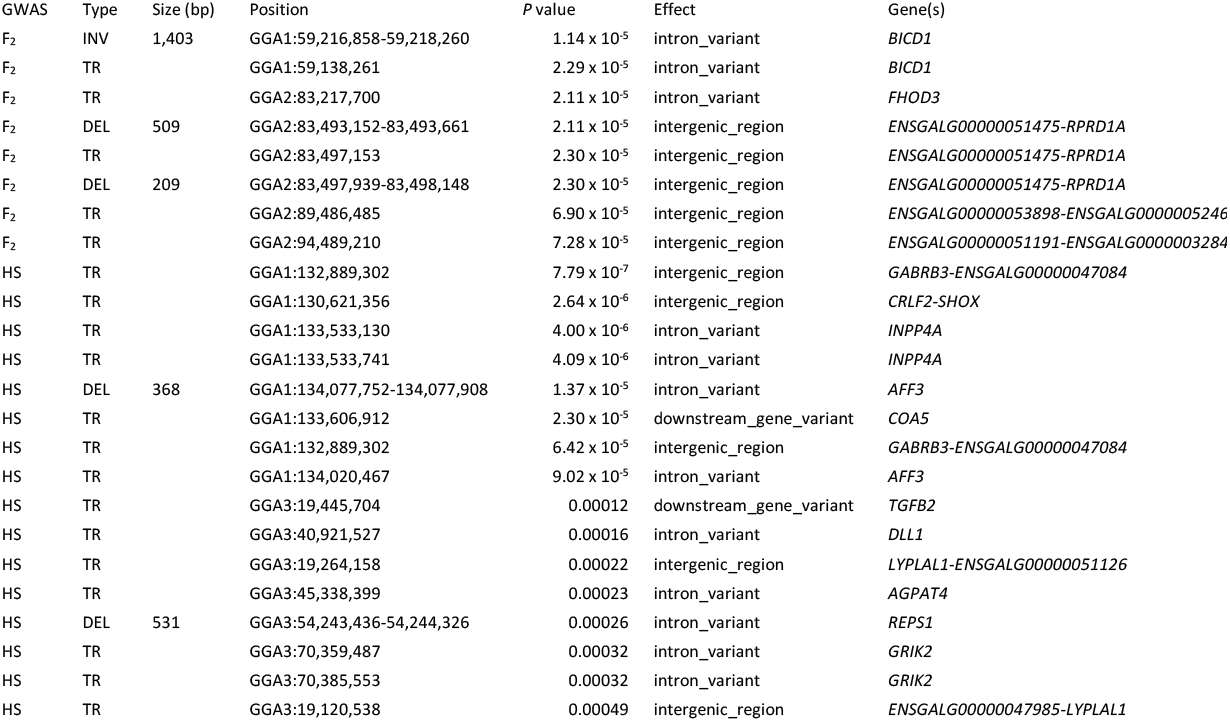
Structural variants (inversions (INV) and deletions (DEL)) and tandem repeats (TRs) that showed the highest association in genome wide association studies for feather pecking behavior in an F_2_ cross and half-sib (HS) families. Variant effects and closest genes were predicted with SnpEff.

| GWAS | Type | Size (bp) | Position | P value | Effect | Gene(s) |
| --- | --- | --- | --- | --- | --- | --- |
| F<sub>2</sub> | INV | 1,403 | GGA1:59,216,858-59,218,260 | 1.14 x 10<sup>-5</sup> | intron_variant | *BICD1* |
| F<sub>2</sub> | TR |  | GGA1:59,138,261 | 2.29 x 10<sup>-5</sup> | intron_variant | *BICD1* |
| F<sub>2</sub> | TR |  | GGA2:83,217,700 | 2.11 x 10<sup>-5</sup> | intron_variant | *FHOD3* |
| F<sub>2</sub> | DEL | 509 | GGA2:83,493,152-83,493,661 | 2.11 x 10<sup>-5</sup> | intergenic_region | *ENSGALG000000051475-RPRD1A* |
| F<sub>2</sub> | TR |  | GGA2:83,497,153 | 2.30 x 10<sup>-5</sup> | intergenic_region | *ENSGALG000000051475-RPRD1A* |
| F<sub>2</sub> | DEL | 209 | GGA2:83,497,939-83,498,148 | 2.30 x 10<sup>-5</sup> | intergenic_region | *ENSGALG000000051475-RPRD1A* |
| F<sub>2</sub> | TR |  | GGA2:89,486,485 | 6.90 x 10<sup>-5</sup> | intergenic_region | *ENSGALG000000053898-ENSGALG00000005246* |
| F<sub>2</sub> | TR |  | GGA2:94,489,210 | 7.28 x 10<sup>-5</sup> | intergenic_region | *ENSGALG000000051191-ENSGALG00000003284* |
| HS | TR |  | GGA1:132,889,302 | 7.79 x 10<sup>-7</sup> | intergenic_region | *GABRB3-ENSGALG000000047084* |
| HS | TR |  | GGA1:130,621,356 | 2.64 x 10<sup>-6</sup> | intergenic_region | *CRLF2-SHOX* |
| HS | TR |  | GGA1:133,533,130 | 4.00 x 10<sup>-6</sup> | intron_variant | *INPP4A* |
| HS | TR |  | GGA1:133,533,741 | 4.09 x 10<sup>-6</sup> | intron_variant | *INPP4A* |
| HS | DEL | 368 | GGA1:134,077,752-134,077,908 | 1.37 x 10<sup>-5</sup> | intron_variant | *AFF3* |
| HS | TR |  | GGA1:133,606,912 | 2.30 x 10<sup>-5</sup> | downstream_gene_variant | *COA5* |
| HS | TR |  | GGA1:132,889,302 | 6.42 x 10<sup>-5</sup> | intergenic_region | *GABRB3-ENSGALG000000047084* |
| HS | TR |  | GGA1:134,020,467 | 9.02 x 10<sup>-5</sup> | intron_variant | *AFF3* |
| HS | TR |  | GGA3:19,445,704 | 0.00012 | downstream_gene_variant | *TGFB2* |
| HS | TR |  | GGA3:40,921,527 | 0.00016 | intron_variant | *DLL1* |
| HS | TR |  | GGA3:19,264,158 | 0.00022 | intergenic_region | *LYPLAL1-ENSGALG000000051126* |
| HS | TR |  | GGA3:45,338,399 | 0.00023 | intron_variant | *AGPAT4* |
| HS | DEL | 531 | GGA3:54,243,436-54,244,326 | 0.00026 | intron_variant | *REPS1* |
| HS | TR |  | GGA3:70,359,487 | 0.00032 | intron_variant | *GRIK2* |
| HS | TR |  | GGA3:70,385,553 | 0.00032 | intron_variant | *GRIK2* |
| HS | TR |  | GGA3:19,120,538 | 0.00049 | intergenic_region | *ENSGALG000000047985-LYPLAL1* |

The only variant that reached genome-wide significance after Bonferroni correction (-Log_10_*P* > 1.41×10^−6^) was a TR in the HS population 126,821 bp downstream of *GABRB3*. The proportion of variance in the phenotype FPD_BC, which is explained by this variant is 0.047. Given heritability estimates of around 0.15 [38–40] the amount of phenotypic variance explained by the lead variant is considerable. To visualize overlaps between the six different sets of results we created a heat map, which shows the –Log_10_*P* values of the top 20 associated genes from the different GWAS (Figure 1b). Numerous genes showed high association signals in both variant classes in the same experimental population. For instance, *ENSGALG00000047084, GABRB3, INPP4A, ATP10A, TSGA10, LIPT1, COA5, UNC50, SHOX, CRLS2*, and *AFF3* showed high association in the HS population for both variant classes. The same holds true for the F_2_ cross (upper block of 20 genes in Figure 1b), although associations were not as strong as in the HS population. Of considerable interest are those genes, which show association in all GWAS: *ENSGALG00000048825, ENSGALG00000051557, ENSGALG00000051256, ENSGALG00000016495, FHOD3, gga-mir-187, GALNT1*, and *PIEZO2*. Predicted targets of the micro RNA (miRNA) *gga-mir-187* are *GABRA1, GABRB2, GABRB3, GABRG1*, and *GABRG2. GABRB2* is also a predicted target of *gga-mir-3524b* (www.targetscan.org accessed January 27, 2022).

## Expression quantitative trait loci

To clarify whether the SVs and TRs that we detected influence the expression of transcripts that we identified in a previous study to be differentially expressed between LFP and HFP [1], we performed an eQTL analysis. We employed the same strategy as in our past study [8] and performed eGWAS for 86 genes from 167 chickens (84 HFP and 83 LFP) from the HS population (Manhattan plots are summarized in Supplementary Information S2). A total of 35,571 SVs and TRs were screened for association with gene expression and we detected 909 genome-wide significant associated signals. SVs and TRs for which we detected genome-wide association with at least 10 DEGs are shown in Table 3. To identify significant gene-gene interactions from those 86 eGWAS we constructed an association weight matrix [32] followed by the detection of significant correlations with the PCIT algorithm [33]. Central to the gene-gene interaction map is the transcription factor *ETV1*. We selected DEGs with a *P* value < 1×10^−4^ for association with the variant, a tandem repeat with the sequence CCCGGCCCG 70 bp upstream of *ETV1* (GGA2: 27337541). With the 23 selected genes (Supplementary Information S1) we performed a transcription factor binding site enrichment analysis with Ciiider and found a significant enrichment (*P* value = 0.013, Log_2_-enrichment = 0.494) of *ETV1* binding sites (Figure 3a) in proximity to those genes (Figure 3b, Supplementary Information S3). To demonstrate specificity we included all available PWMs for members of the ETS family of transcription factors: *ETV2* – *ETV6*. The only other member of the TF family for which transcription factor binding site enrichment was detected was *ETV4*, but the *P* value was not significant. The complete results of the analysis are summarized in Supplementary Information S4.

**Table 3.** Large deletions (DEL) and tandem repeats (TR) that showed genome-wide significant association in at least 10 expression genome-wide association studies for genes that are differentially expressed in brains of high and low feather pecking chickens.

| Type | Size<br>(bp) | Position | Effect | Gene(s) | No. of associations |
| --- | --- | --- | --- | --- | --- |
| TR |  | GGA25:1,465,942 | intergenic_region | *ENSGALG00000055092-IQGAP3* | 53 |
| TR |  | GGA25:3,317,155 | intron_variant | *S100A16* | 47 |
| TR |  | GGA17:6,359,797 | intergenic_region | *ENSGALG00000035908-ENSGALG00000047338* | 39 |
| DEL | 294 | GGA8:3,889,333-3,889,627 | intron_variant | *RO60* | 30 |
| TR |  | GGA2:56,803,419 | intron_variant | *ATP9B* | 22 |
| TR |  | GGA2:56,944,739 | intergenic_region | *SALL3-ENSGALG00000054175* | 22 |
| TR |  | GGA17:5,355,006 | upstream_gene_variant | *SLC27A4* | 15 |
| TR |  | GGA27:3,942,783 | intron_variant | *GPATCH8* | 11 |
| TR |  | GGA27:3,976,916 | frameshift_variant | *GPATCH8* | 11 |
| TR |  | GGA27:3,977,142 | intron_variant | *GRN* | 11 |
| TR |  | GGA27:3,988,533 | intergenic_region | *TMEM98-ARHGAP27* | 11 |
| TR |  | GGA27:4,009,079 | intron_variant | *PLEKHM1* | 11 |
| TR |  | GGA27:4,022,832 | intron_variant | *ENSGALG00000047878* | 11 |

## Discussion

Numerous studies have been conducted to unravel the genetics behind FP behavior in chickens, as reviewed in [12, 41], but none of these focused on the analysis of structural genetic variation. We combined data of two well-described experimental crosses, an F_2_ design and a HS population, and performed multiple GWAS and an eQTL analysis on different classes of genetic variants. The lead variant from these analyses is a TR 126 kb downstream of the *GABRB3* gene, which explains a considerable amount of the observed phenotypic variance. Furthermore, miRNA *gga-mir-187* is strongly associated in all conducted GWAS (Figure 1b), and among predicted targets of this regulatory RNA are several GABA receptor genes. Based on whole-brain transcriptome analyses of the light response of HFP we previously postulated that downregulation or missing upregulation of GABA receptor expression is caused by miRNA dysregulation due to low levels of *Dicer1* expression in HFP brains [14]. The findings presented here provide further evidence for a disturbance in miRNA regulation and involvement of the GABAergic system in FP behavior. Since these chickens have been selected for feather pecking behavior based on their estimated breeding values for multiple generations, it is possible that mutations leading to low expression of several GABA receptors have been accumulating. Previous studies with the same experimental population already pointed into that direction [2, 13]. But these are not only limited to GABA receptor genes, miRNAs or genes encoding miRNA processing proteins. By conducting an eQTL analysis with SVs and TRs we were able to confirm an involvement of *DMD* (dystrophin) in the regulation of genes that are differentially expressed between HFP and LFP. We already discovered *DMD* in an eQTL analysis with SNPs and InDels [8]. Since dystrophin is a direct regulator of GABA receptor clustering [42], this adds another layer to the GABA receptor disturbance in HFP brains. Several other genes that appear in the gene-gene interaction map based on significant AWM correlations (Figure 2) have been connected to the GABAergic system. These include *COQ4* [43], *ETV1* [44], *NEURL1* [45], *SLC25A26*, and *SLC27A4* [46]. In this regard, *ETV1* is of considerable interest since it is the only transcription factor that we identified in our eQTL analysis with SVs and TRs. According to the Human Protein Atlas *ETV1* expression is specific to salivary glands and the brain, specifically the cerebellum and thalamus (https://www.proteinatlas.org/ENSG00000006468-ETV1; accessed August 2022) *ETV1* is associated with the differential expression of 23 genes between HFP and LFP and *ETV1* binding sites are enriched in proximity to these genes (Figure 3). Among these putative ETV1 targets are *CSF2RB*, which has been associated with major depression and schizophrenia [47], and *MIS18BP1*, a gene implicated in autism spectrum [48]. Furthermore, *ETV1* antibody coimmunoprecipitated the GABAA receptor α6 (GABAARα6) promoter region in mice [44] and is a regulator of gene expression in CD4 and CD8 T cells [49]. Hence, *ETV1* may not only impact GABA receptor expression but might also participate in neurodevelopment. As demonstrated by Pasciuto et *al*. a lack of CD4 T cells in the brain of mice leads to excess immature neuronal synapses and behavioral abnormalities. Among the top DEGs identified by single-cell RNAseq were the transcription factors *Klf2* and *Klf4* [16]. We previously postulated that FP behavior is the result of disturbances in embryonic neurodevelopment and identified *KLF14* as a major regulator in this regard [8]. In that study, we also identified a Smad4 domain-containing transcription factor, namely *ENSGALG00000042129* (*CHTOP*), but we were not able to detect enrichment of binding sites for that transcription factor in proximity to its associated DEGs. However, in the light of the fact that ETV1 forms functional complexes with Smad4 [50] we propose a model in which the transcription factors *ETV1, CHTOP*, and *KLF14* have an additive effect on the density of T cells in the developing brain of HFP. This might lead to structural abnormalities in the brains of HFP and may explain, at least in part, their abnormal behavior. Multiple studies point towards a central role of Krüppel-Like factors in neurodevelopment and behavior [51–55]. However, conclusive results on the function of Krüpple-like factors in the neurodevelopment in chickens have not been reported yet and research is hampered by erroneous annotation of KLF orthologues [56].

**Figure 2:**
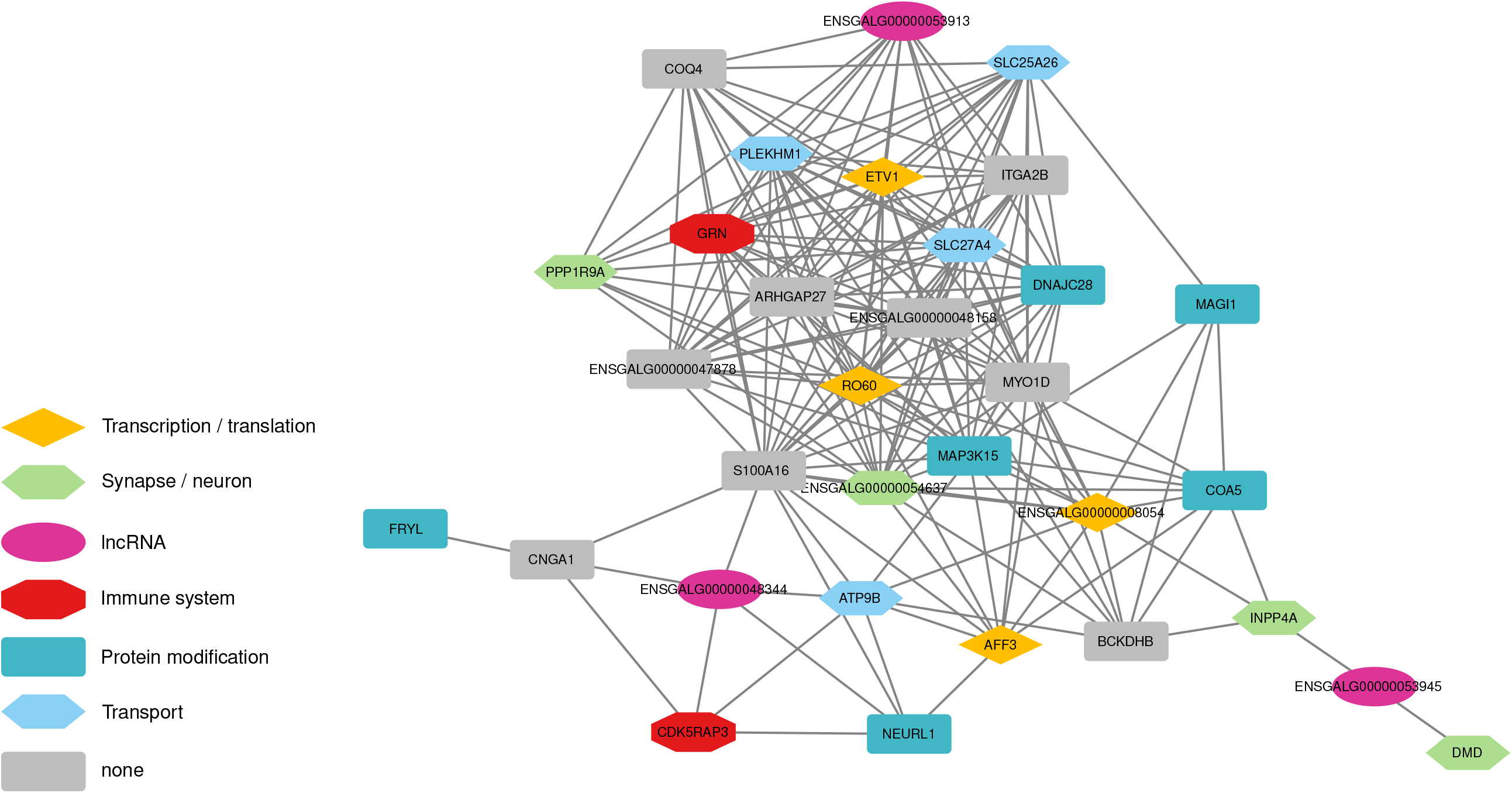
Gene-gene interaction map of significant correlations that were identified from an association weight matrix on expression genome-wide association studies with the PCIT algorithm.

**Figure 3:**
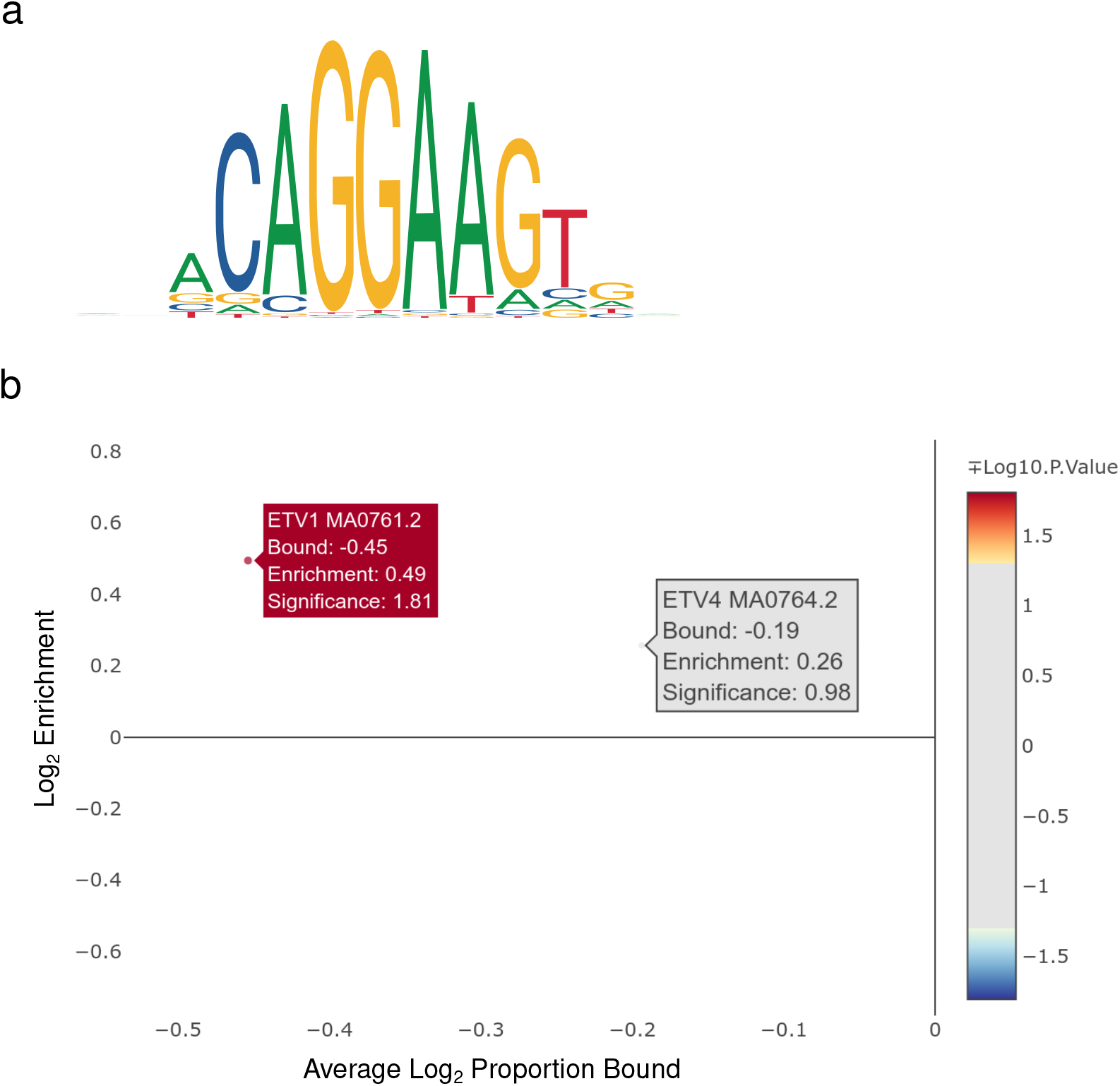
(a) Graphical representation of the binding motif frequency matrix of the *ETV1* transcription factor. (b) Result of transcription factor binding site enrichment analysis for *ETV1* with genes from expression genome-wide associations studies with *P* values > 1 × 10^−4^.

Apart from *DMD* we also discovered the genes *PPP1R9A, INPP4A*, and *COA5* in both eQTL analyses. One of which was performed with SVs and TRs (Figure 2) and one with SNPs and InDels [8]. In schizophrenia and bipolar disorder, the prefrontal cortical expression of *PPP1R9A* was altered [57] and its gene product Neurabin regulates anxiety-like behavior in adult mice [58]. *INPP4A* has been implicated in multiple neurological conditions, namely schizophrenia, autism, epilepsy, and intellectual disability [59–61]. An additional link to schizophrenia is a frameshift variant in the *GPATCH8* gene in exon 9 of transcript GPATCH8-201 (Table 3), an ortholog of *ZNF804A*, which has been shown to impact various mental illnesses via pre-mRNA processing [62]. We discovered multiple links of FP to human psychiatric disorders in our previous studies [1, 2, 14]. The fact that methylation of *KLF14* correlates with psychosis severity in schizophrenia patients [63] strengthens our argument for the applicability of HFP chickens in the research of the basic mechanisms involved in human psychiatric disorders. Furthermore, by conducting multiple GWAS on two experimental populations of FP chickens with different variant classes with whole genome marker density we identified numerous genes that have previously been linked to psychiatric disorders or other neurological conditions or phenotypes (Table 4). These genes coherently fit the mechanistic hypotheses raised so far and at the same time underpin the complex nature of this trait. Among these are anxiety, depression, intellectual disability, autism, compulsive behavior, drug addiction, bipolar disorder, and schizophrenia. This raises the question, which common mechanisms these conditions share, which is a current matter of debate [64]. Blokhin *et al*. proposed that future treatment of human psychiatric disorders should be tailored under the use of a genome-wide “targetome”. The strong genetic burden of HFP chickens with mutations affecting neuropsychiatric genes makes this line of chickens a valid model system to study the effects of tailored drug treatment strategies.

**Table 4:**
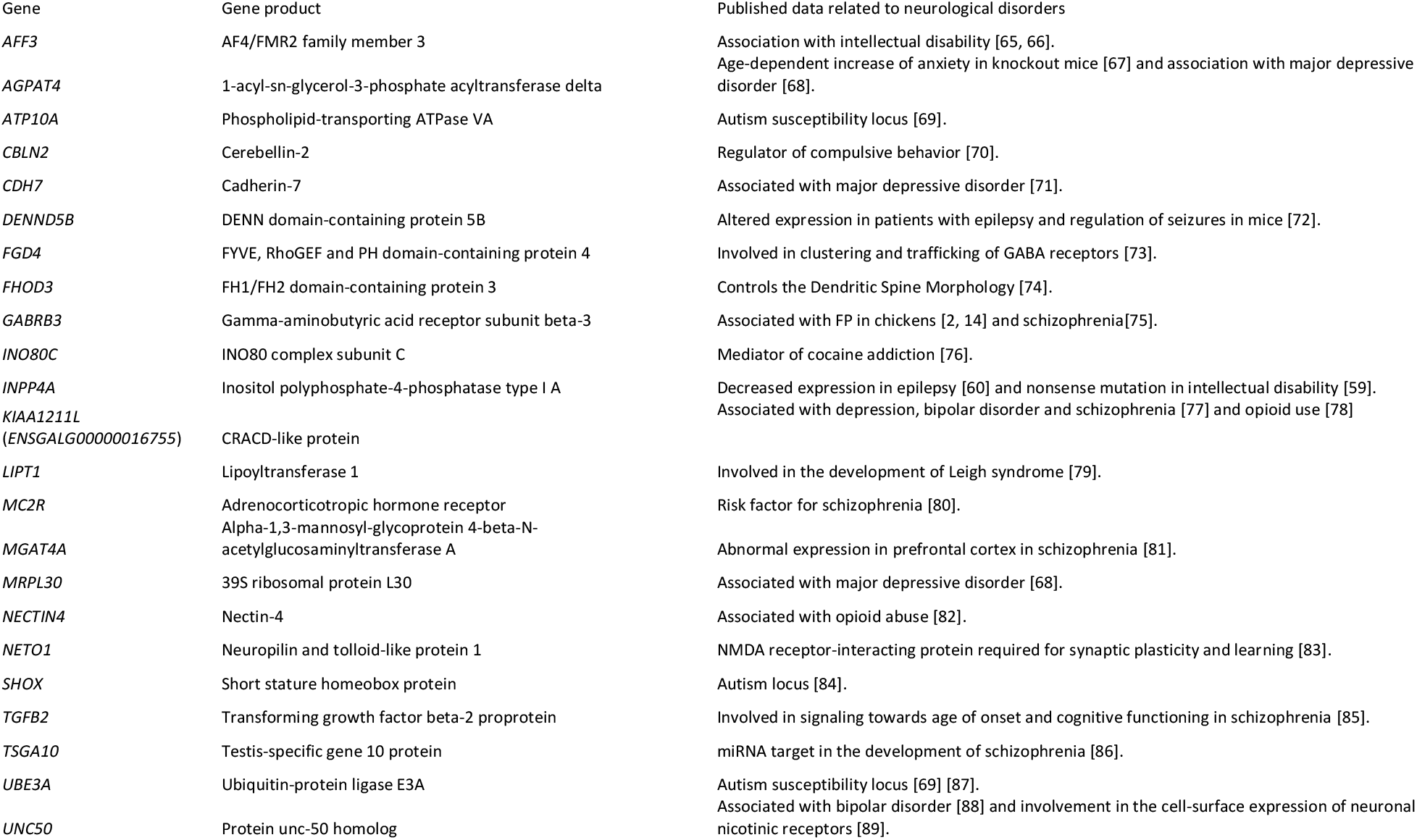
Summary of published data related to neurological disorders on the 20 highest associated genes from each genome wide association study on feather pecking behavior in laying hens.

## Supporting information

Differentially expressed genes between high and low feather pecking chickens, which were associated with a tandem repeat 70 bp downstream of the ETV1

Manhattan plots of expression genome wide associations studies conducted on genes, which were differentially expressed between high and low feather pe

Graphical representation of ETV1 binding sites in proximity to associated differentially expressed genes

Results of transcription factor binding site enrichment analysis with Ciiider for transcription factors of the ETS (E twenty-six) family to differenti

## Statements and Declarations

### Availability of data and materials

All methods applied here have been outlined in previous studies [1, 2]. The raw RNA sequencing data has been deposited at the NCBI Sequence Read Archive (BioProject ID PRJNA656654) and the raw whole genome sequencing data as well (BioProject ID PRJNA664592).

### Competing interests

The authors declare to have no competing interests of any kind.

### Ethics approval

The research protocol was approved by the German Ethical Commission of Animal Welfare of the Provincial Government of Baden-Wuerttemberg, Germany (code: HOH 35/15 PG, date of approval: April 25, 2017).

### Consent to participate

Not applicable

### Consent for publication

Not applicable

### Funding

The study was funded by the German Research Foundation (DFG) under file numbers TE622/4-2 and BE3703/8-2. The funders had no role in study design, data collection and analysis, and interpretation of data and in writing the manuscript. Publication fee was covered by the Open Access Publication Funds of the Göttingen University

### Contributions

Conceptualization: Clemens Falker-Gieske, Jens Tetens; Methodology: Clemens Falker-Gieske, Jens Tetens; Formal analysis and investigation: Clemens Falker-Gieske; Writing - original draft preparation: Clemens Falker-Gieske; Writing - review and editing: Jens Tetens, Jörn Bennewitz; Funding acquisition: Jens Tetens, Jörn Bennewitz; Resources: Jens Tetens, Jörn Bennewitz; Supervision: Jens Tetens.

## Acknowledgements

We acknowledge support by the Open Access Publication Funds of the Göttingen University.

## Supplementary Information

Supplementary Information S1: Differentially expressed genes between high and low feather pecking chickens, which were associated with a tandem repeat 70 bp downstream of the *ETV1* transcription factor.

Supplementary Information S2: Manhattan plots of expression genome wide associations studies conducted on genes, which were differentially expressed between high and low feather pecking chickens.

Supplementary Information S3: Graphical representation of *ETV1* binding sites in proximity to associated differentially expressed genes.

Supplementary Information S4: Results of transcription factor binding site enrichment analysis with Ciiider for transcription factors of the ETS (E twenty-six) family to differentially expressed genes that were associated with a tandem repeat downstream of *ETV1*.

## Notes

### Competing Interest Statement

The authors have declared no competing interest.

