## Supplementary material for "Structural variation and eQTL analysis in two experimental populations of chickens divergently selected for feather pecking behavior": Manhattan plots of expression genome wide associations studies conducted on genes, which were differentially expressed between high and low feather pe

Supplementary Information S2: Manhattan plots of expression genome wide associations studies conducted on genes, which were differentially expressed between high and low feather pecking chickens.

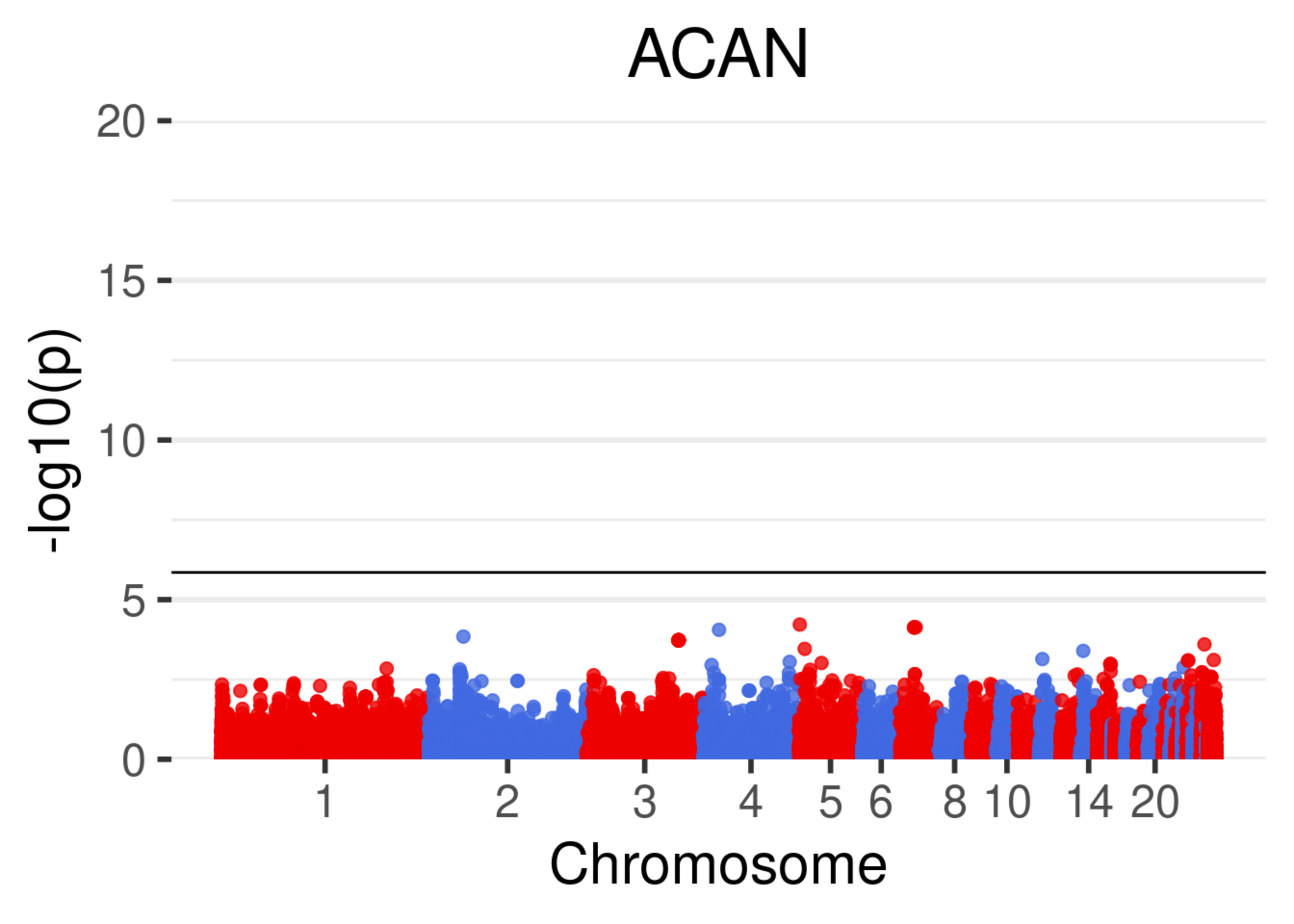
ACAN
20
15
10
5
0
-log10(p)
1
2
3
4
5
6
8
10
14
20
Chromosome

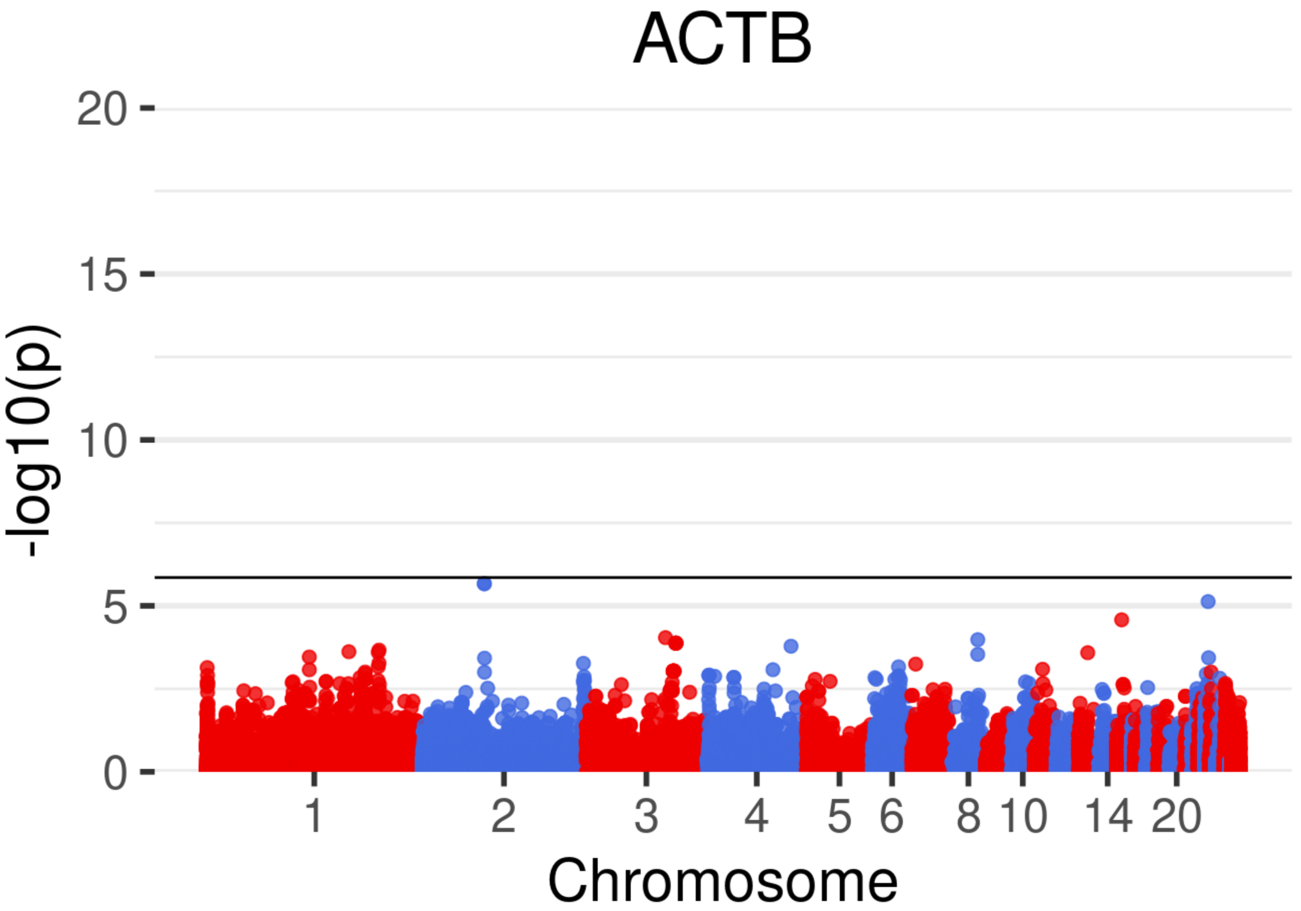

ACTB
20
15
10
5
0
-log10(p)
1
2
3
4
5
6
8
10
14
20
Chromosome

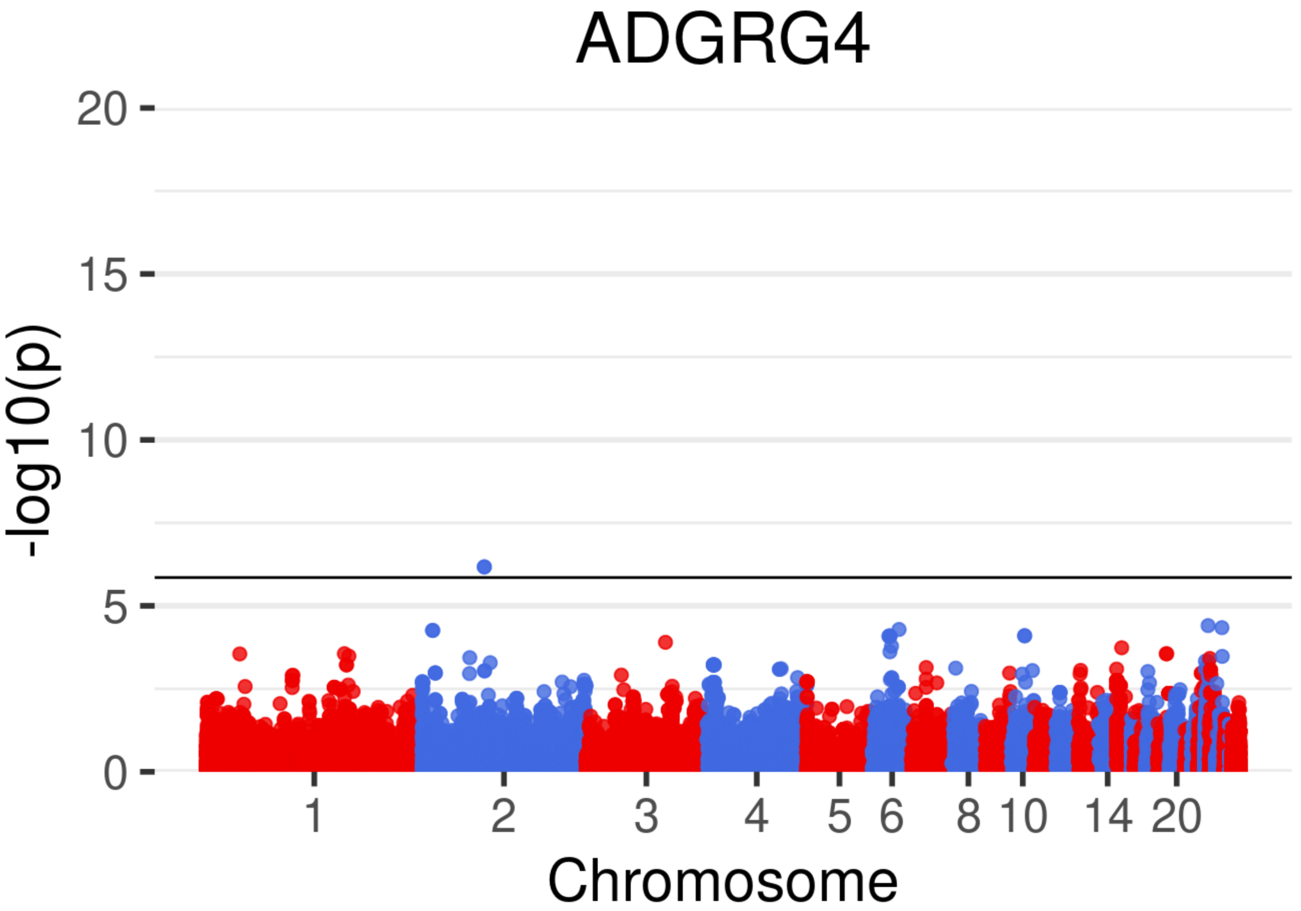
ADGRG4
20
15
10
5
0
-log10(p)
1
2
3
4
5
6
8
10
14
20
Chromosome

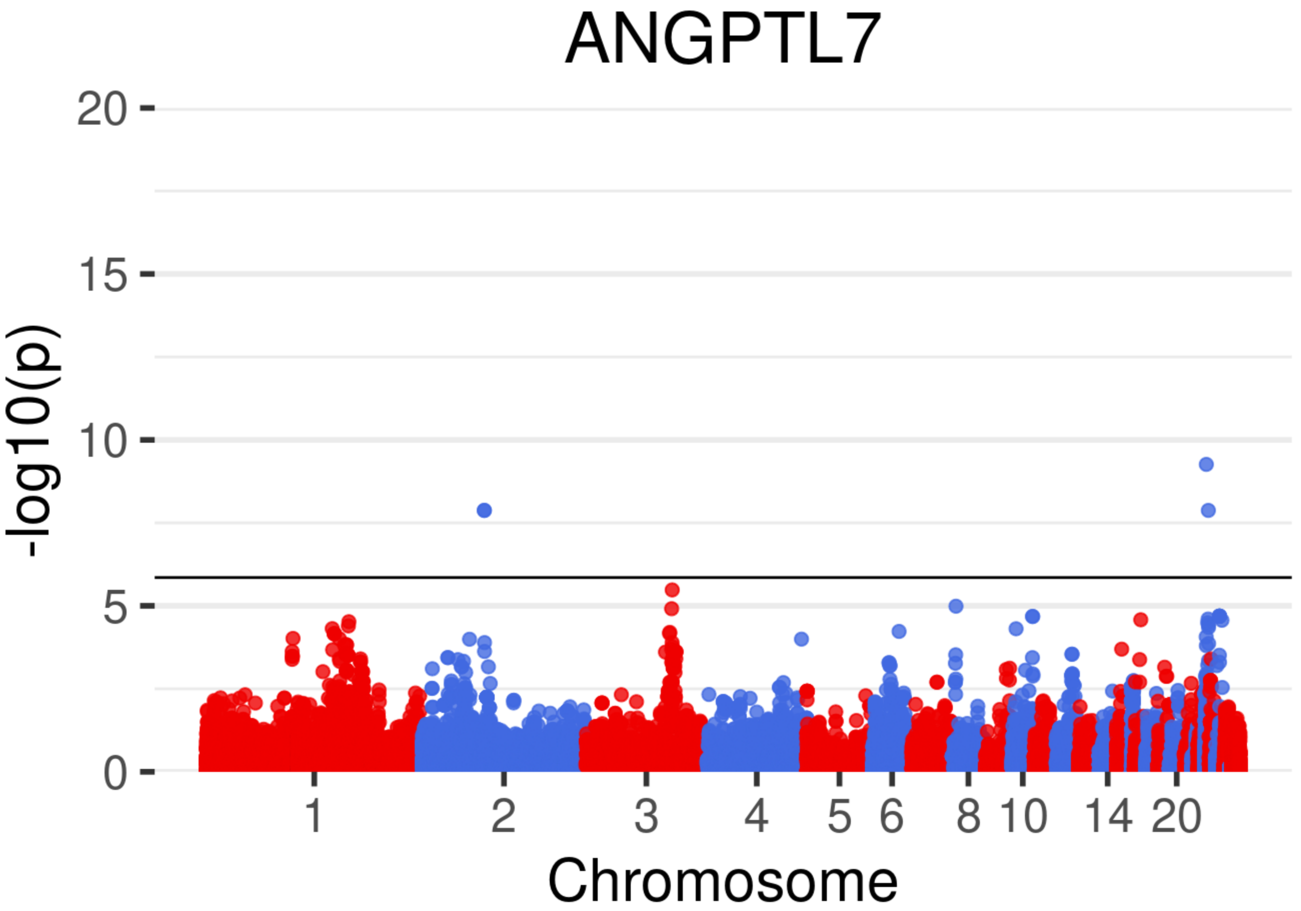

ANGPTL7
20
15
10
5
0
-log10(p)
1
2
3
4
5
6
8
10
14
20
Chromosome

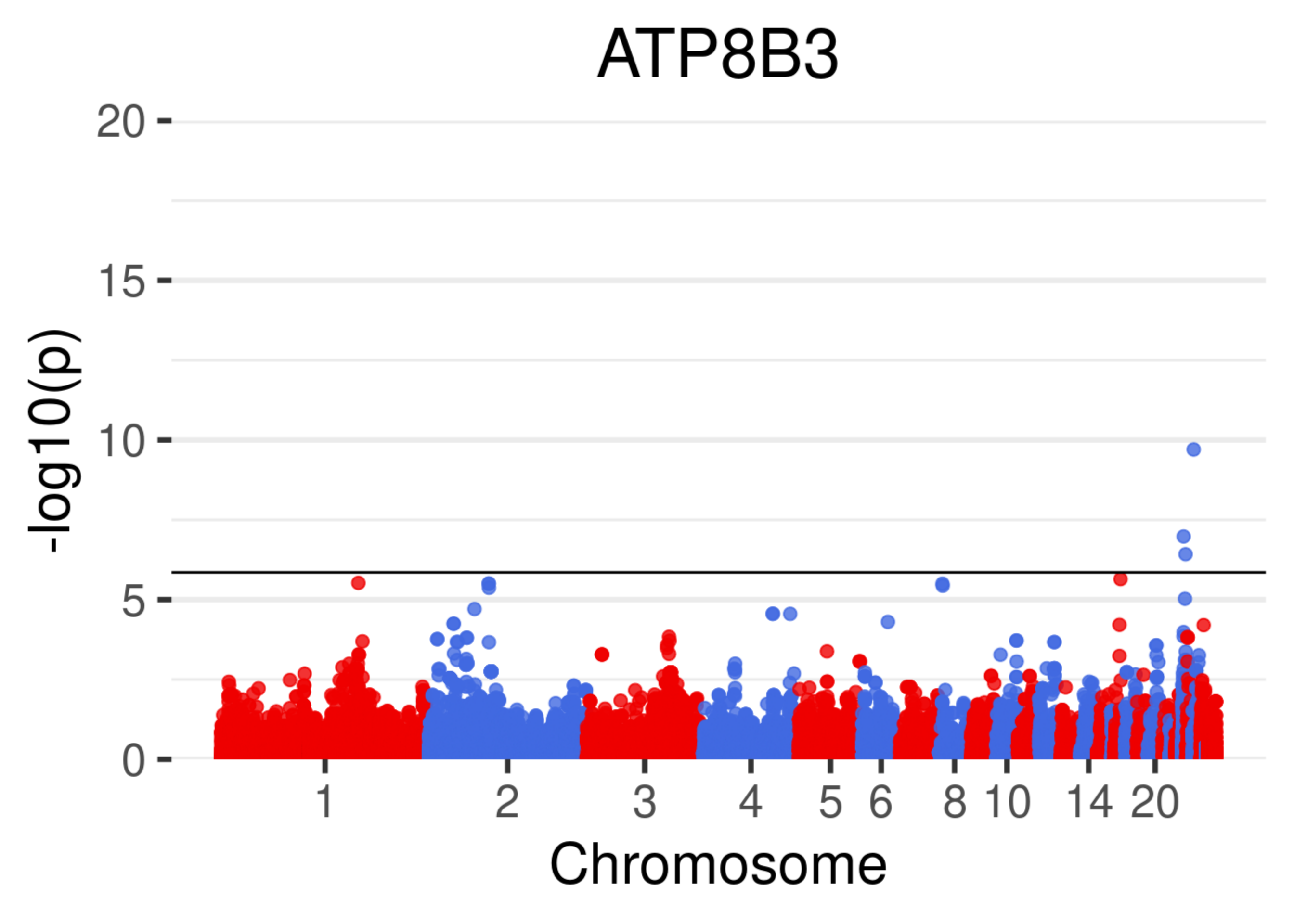
ATP8B3
-log10(p)
20
15
10
5
0
1
2
3
4
5
6
8
10
14
20
Chromosome

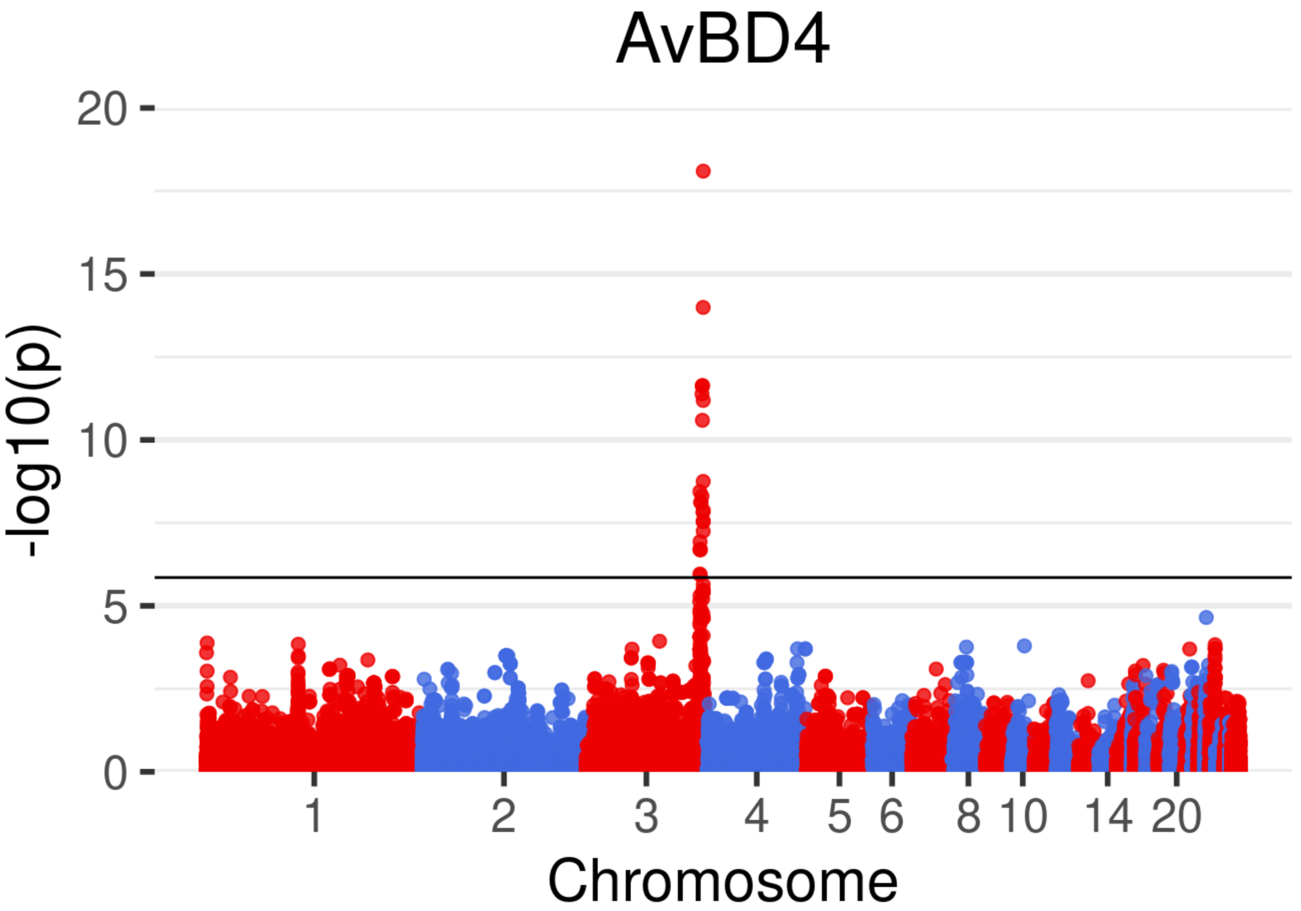
AvBD4
20
15
10
5
0
-log10(p)
1
2
3
4
5
6
8
10
14
20
Chromosome

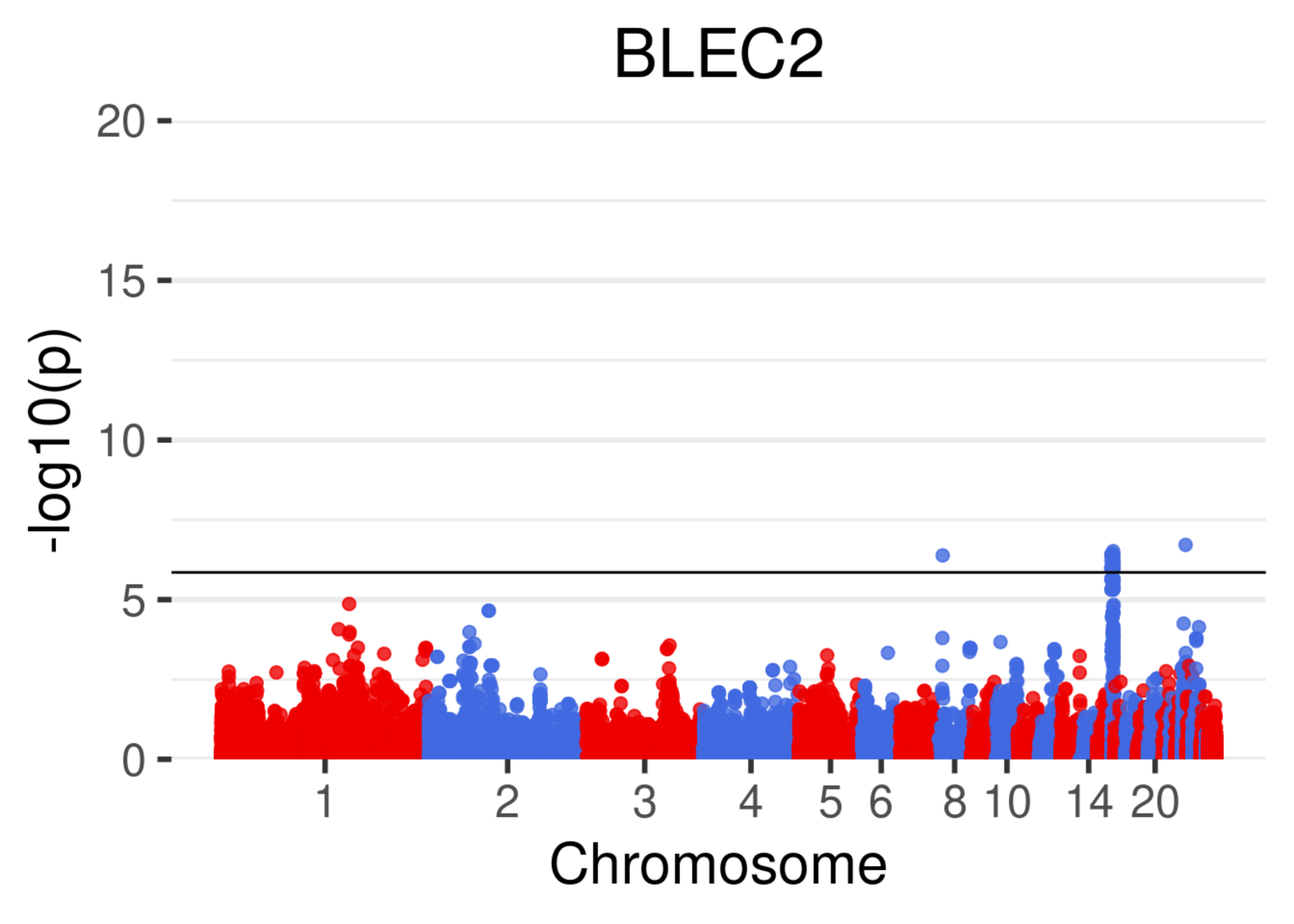
BLEC2
-log10(p)
20
15
10
5
0
1
2
3
4
5
6
8
10
14
20
Chromosome

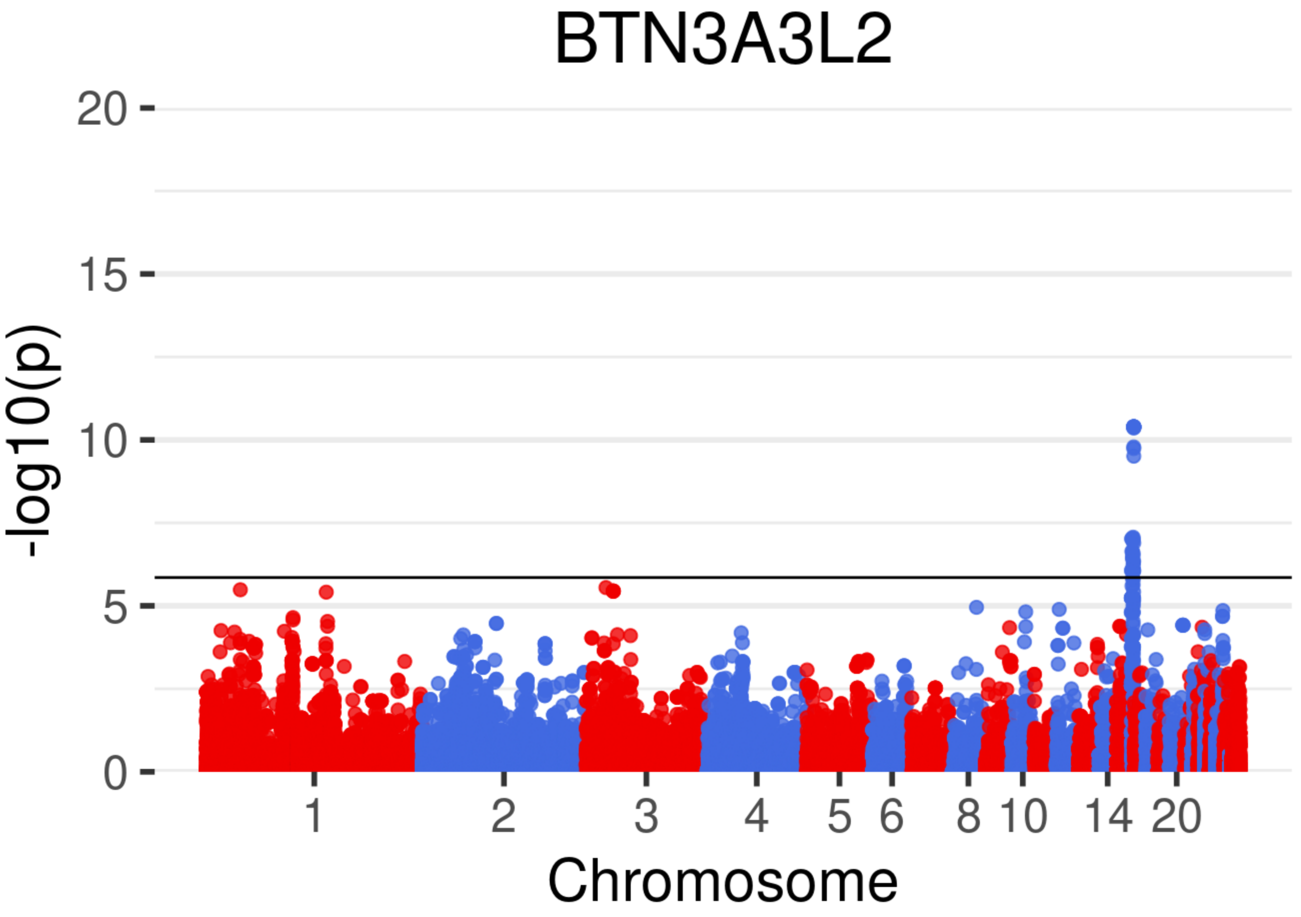
BTN3A3L2
20
15
10
5
0
-log10(p)
1
2
3
4
5
6
8
10
14
20
Chromosome

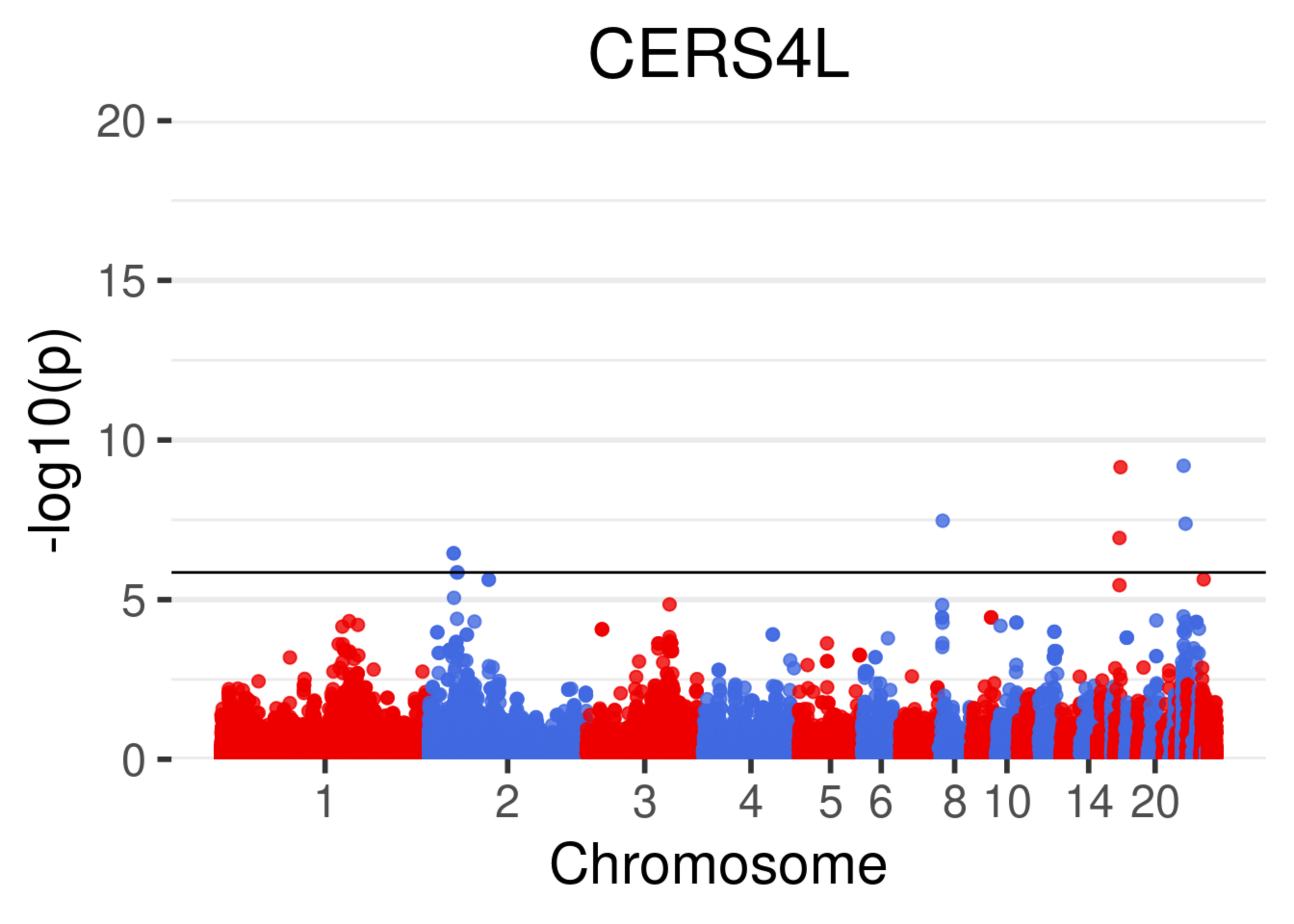
CERS4L
20
15
10
5
0
-log10(p)
1
2
3
4
5
6
8
10
14
20
Chromosome

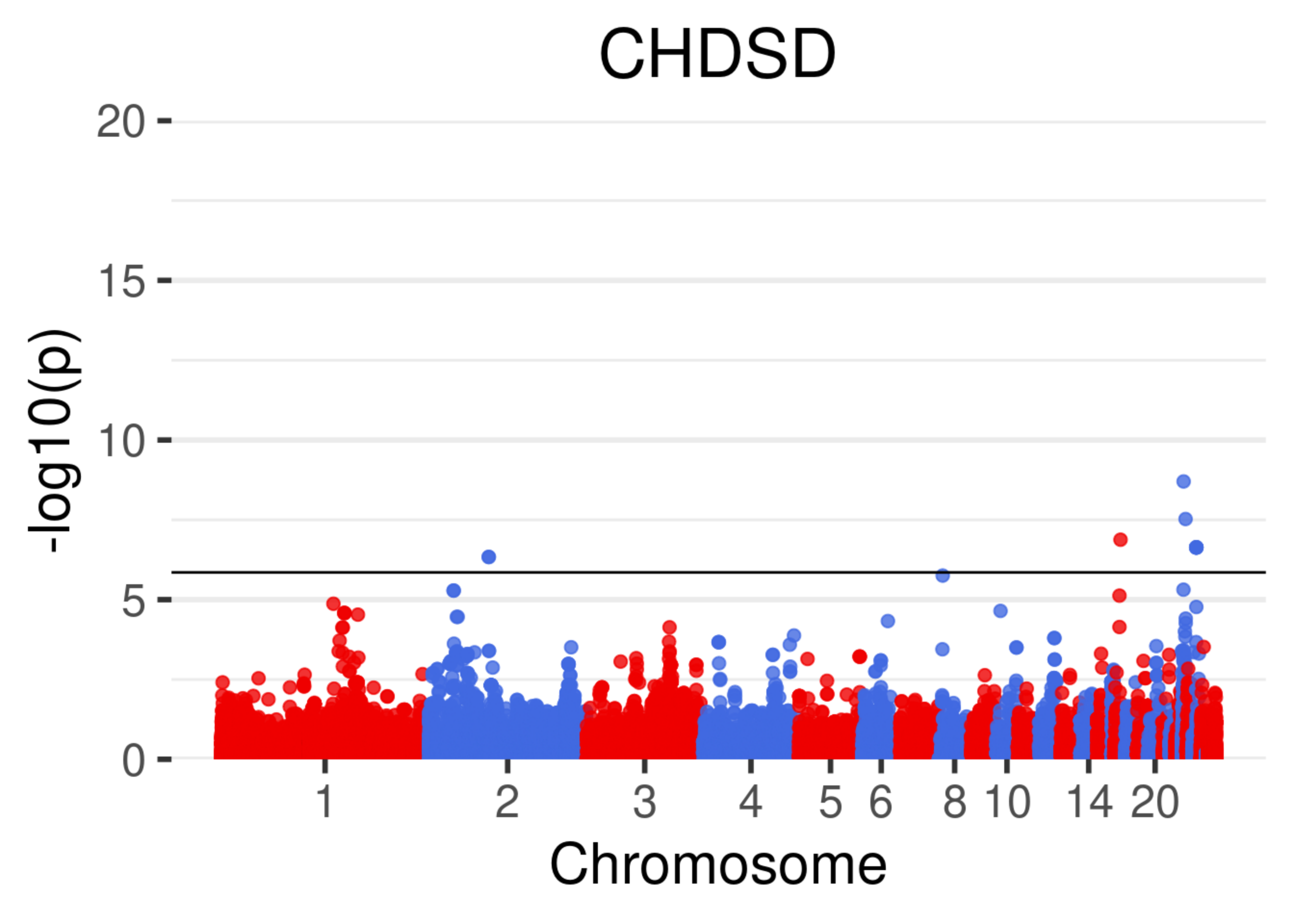
CHDSD
20
15
10
5
0
-log10(p)
1
2
3
4
5
6
8
10
14
20
Chromosome

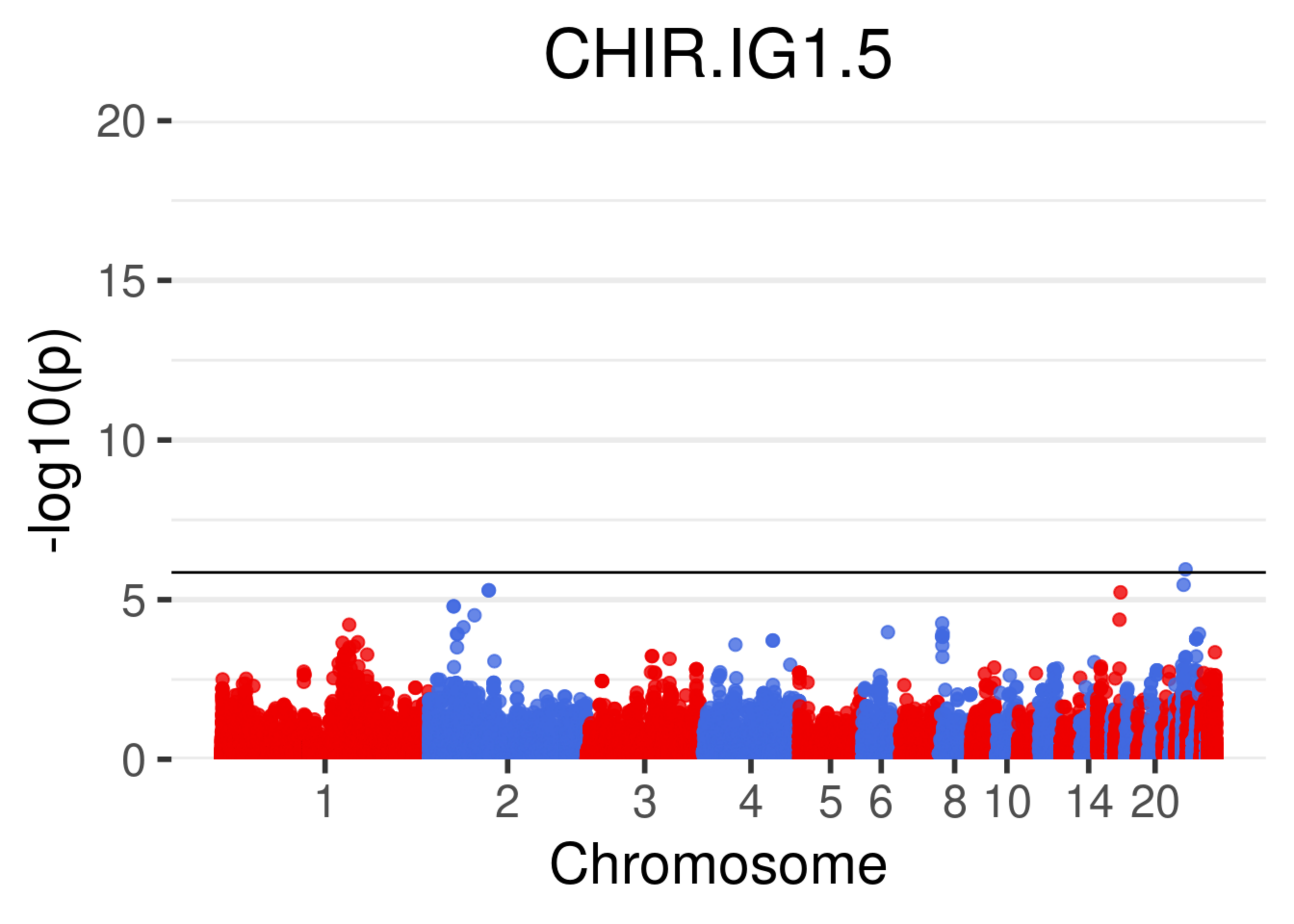
CHIR.IG1.5
20
15
10
5
0
-log10(p)
1
2
3
4
5
6
8
10
14
20
Chromosome

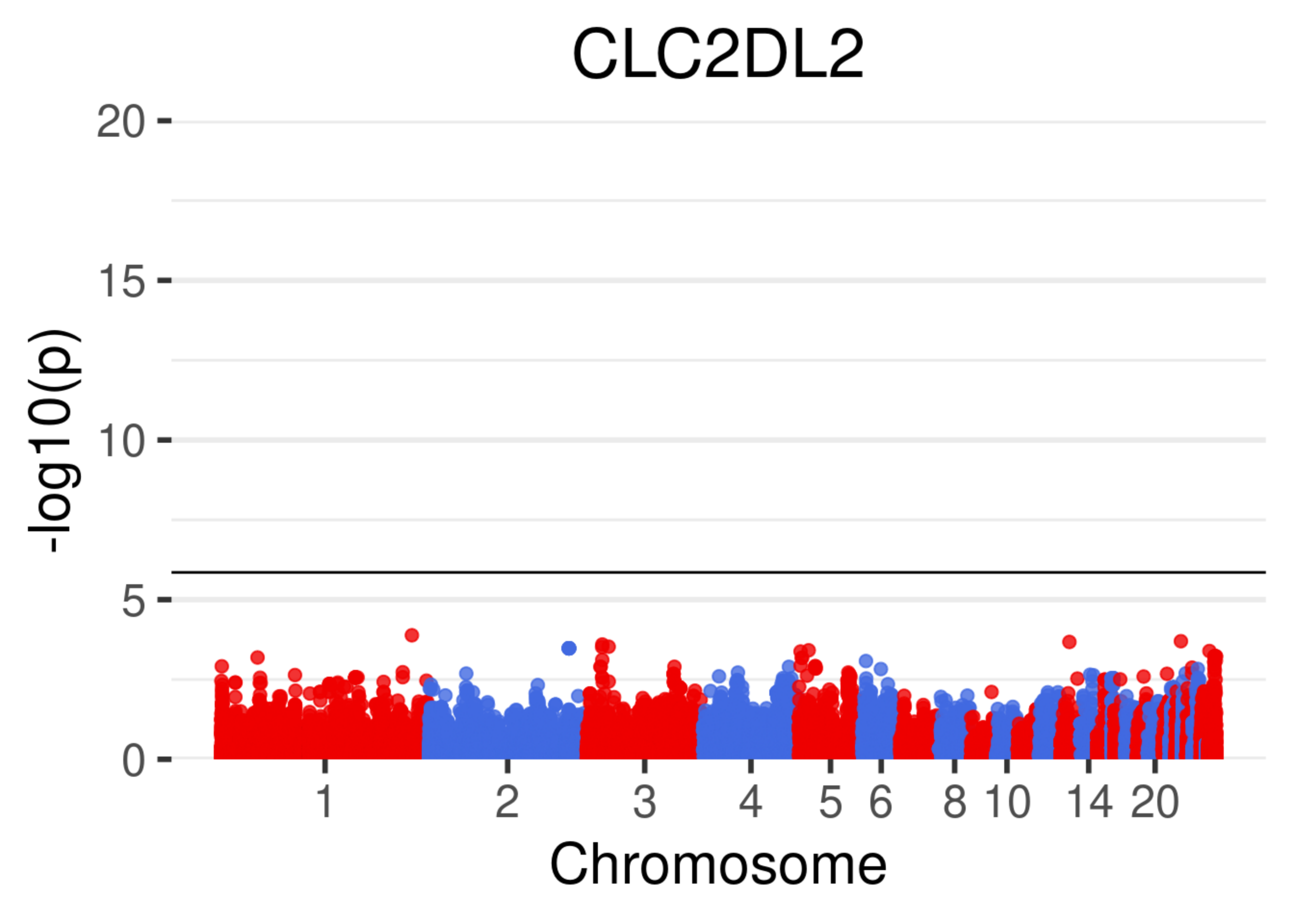
CLC2DL2
-log10(p)
20
15
10
5
0
1
2
3
4
5
6
8
10
14
20
Chromosome

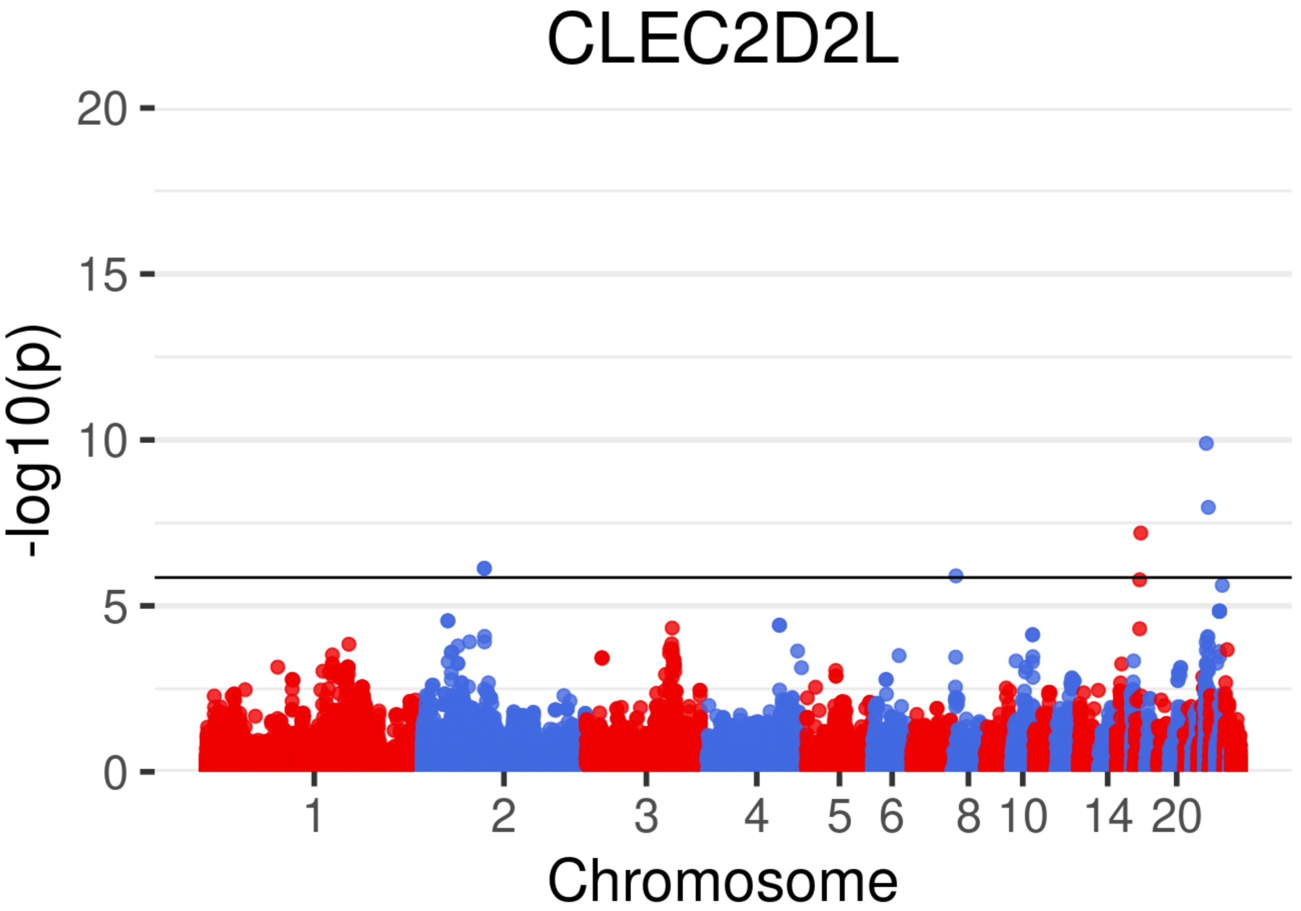

CLEC2D2L
-log10(p)
20
15
10
5
0
1
2
3
4
5
6
8
10
14
20
Chromosome

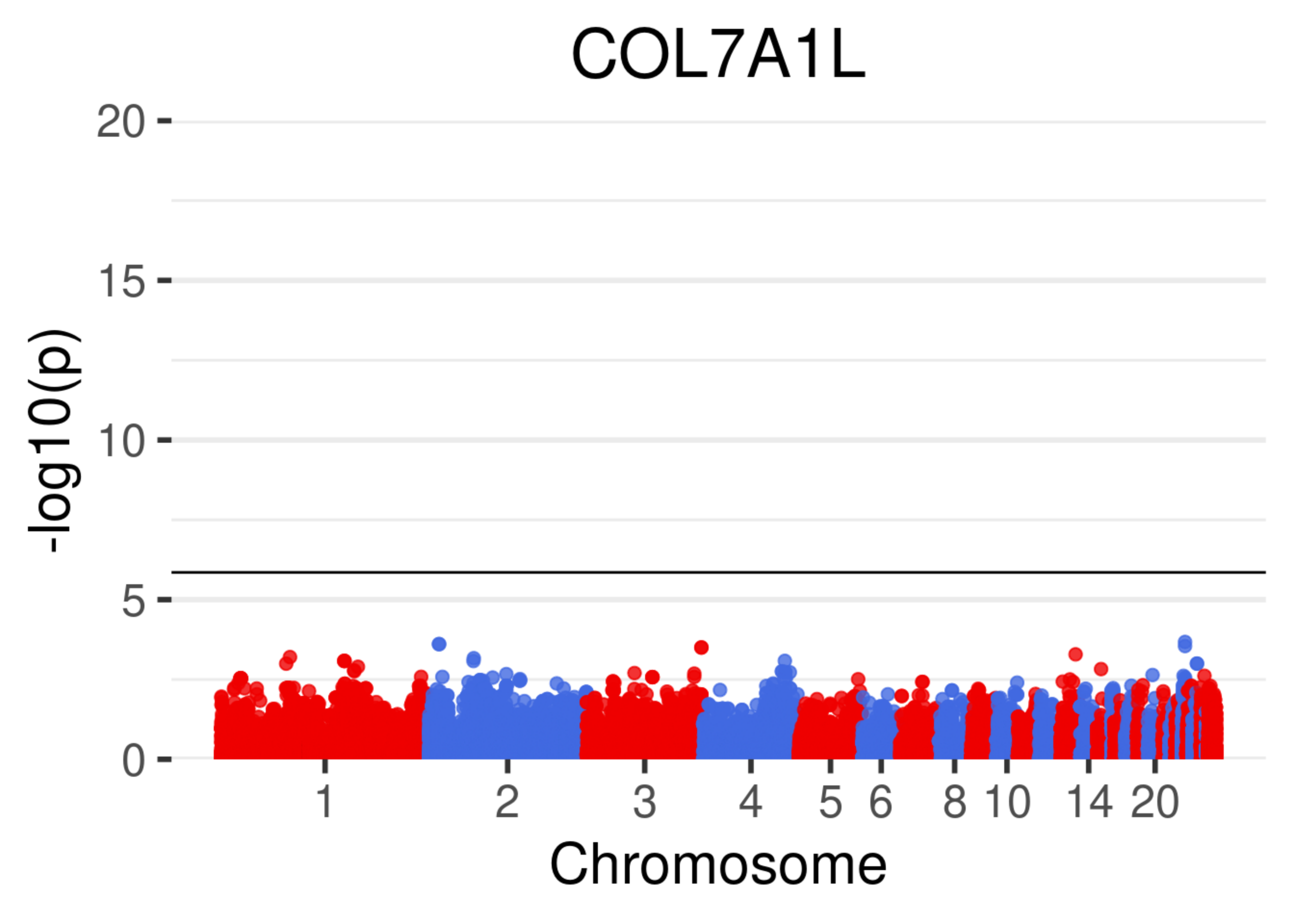
COL7A1L
20
15
10
5
0
-log10(p)
1
2
3
4
5
6
8
10
14
20
Chromosome

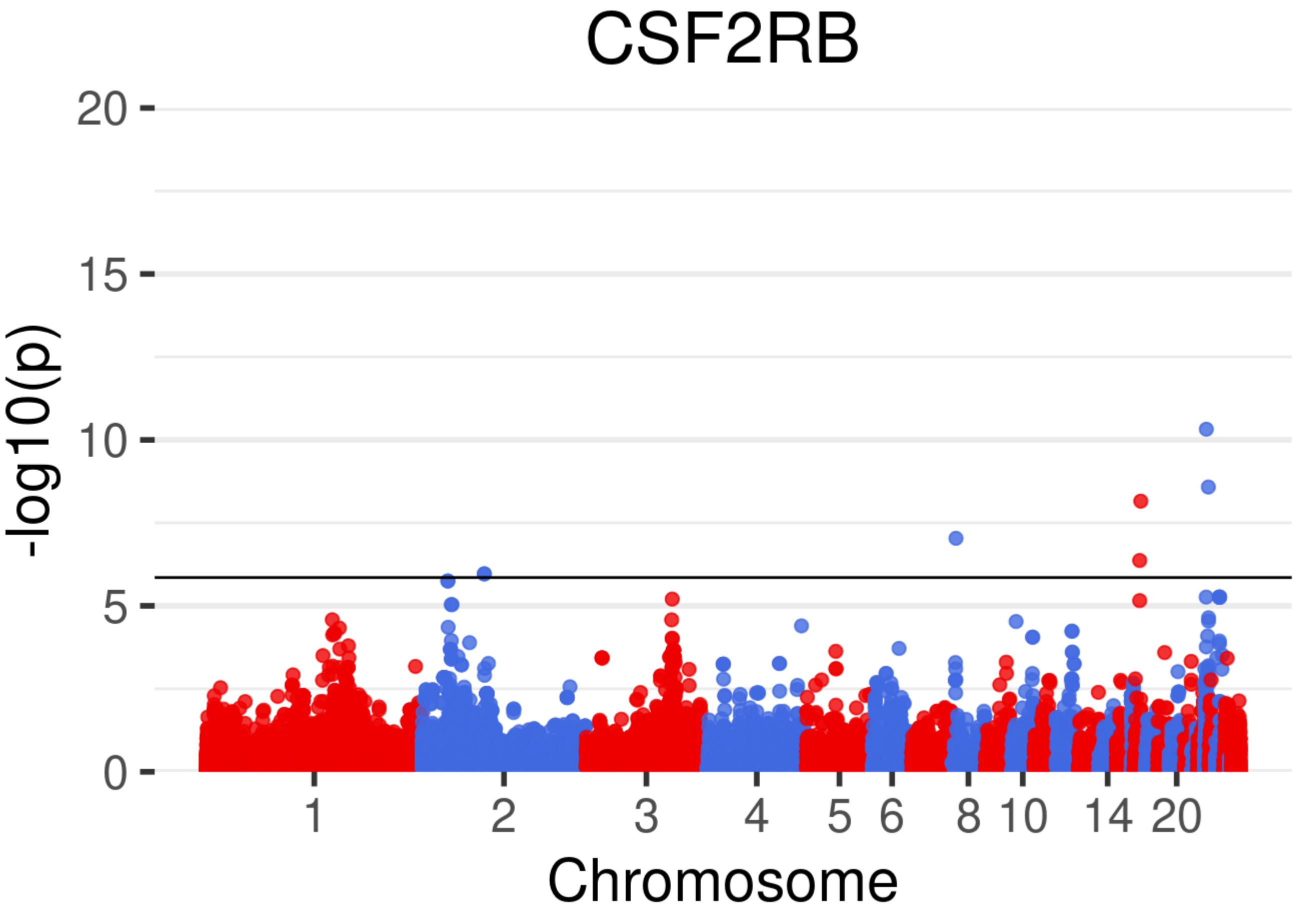
CSF2RB
-log10(p)
20
15
10
5
0
1
2
3
4
5
6
8
10
14
20
Chromosome

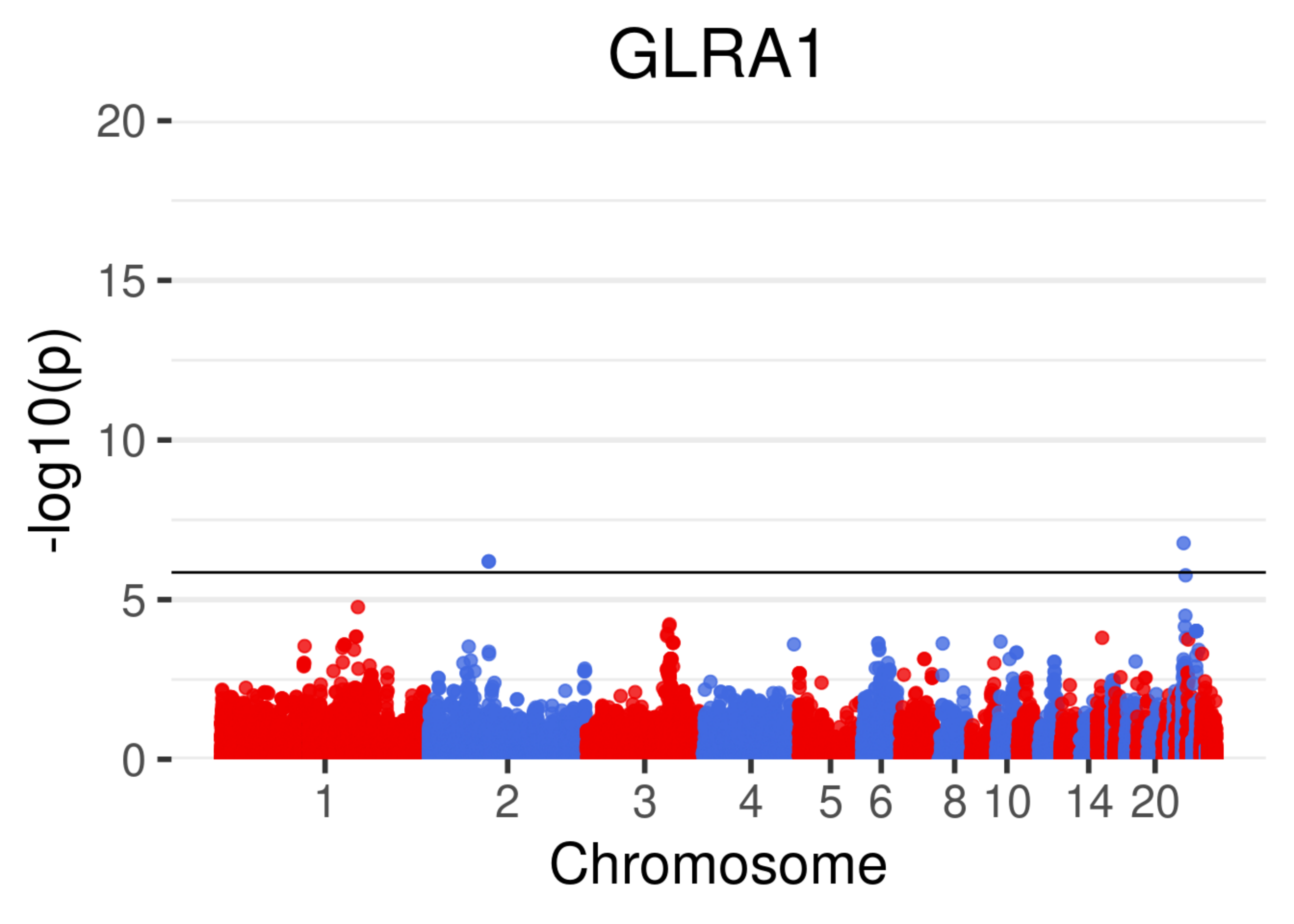
GLRA1
20
15
10
5
0
-log10(p)
1
2
3
4
5
6
8
10
14
20
Chromosome

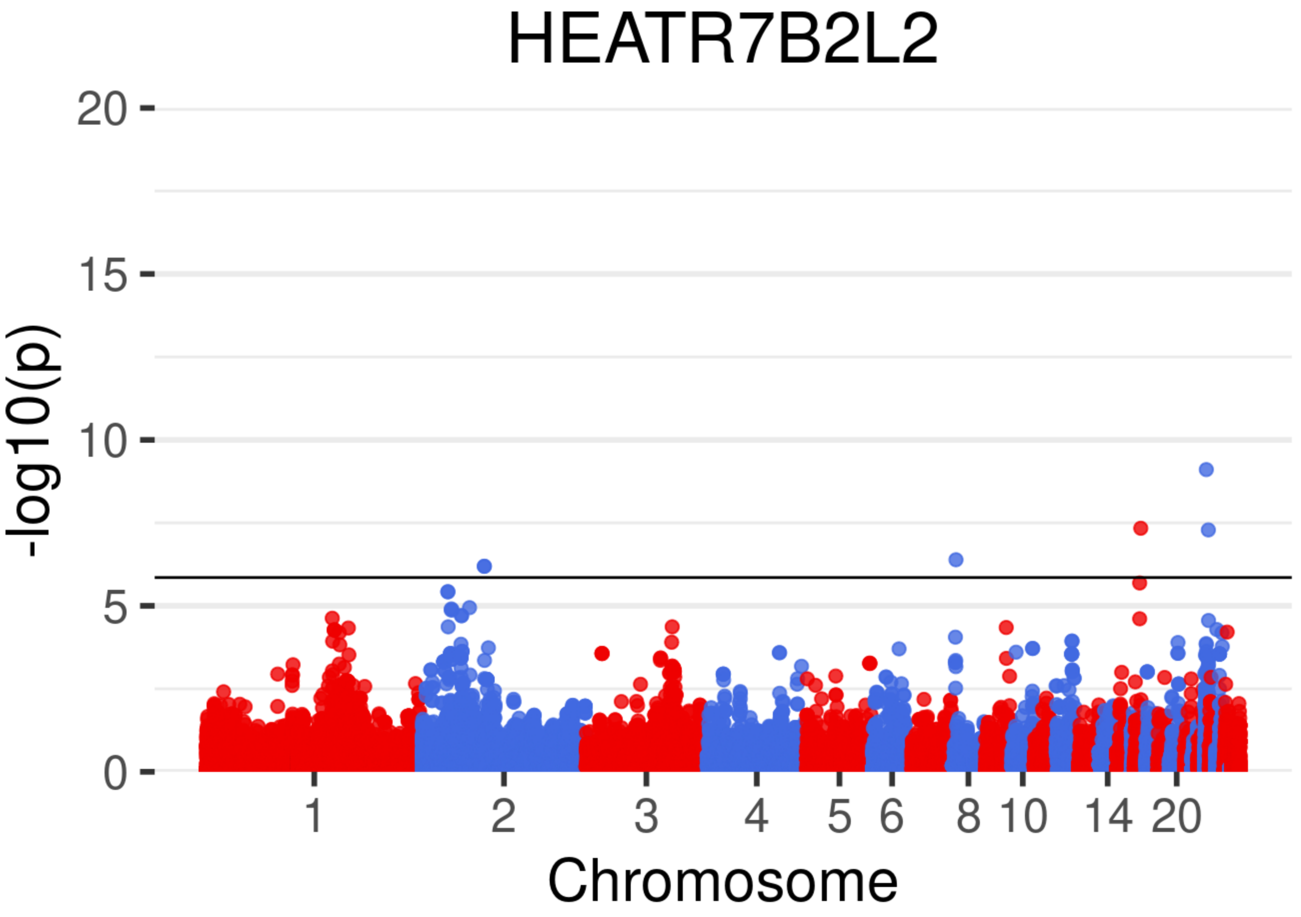

HEATR7B2L2
20
15
10
5
0
-log10(p)
1
2
3
4
5
6
8
10
14
20
Chromosome

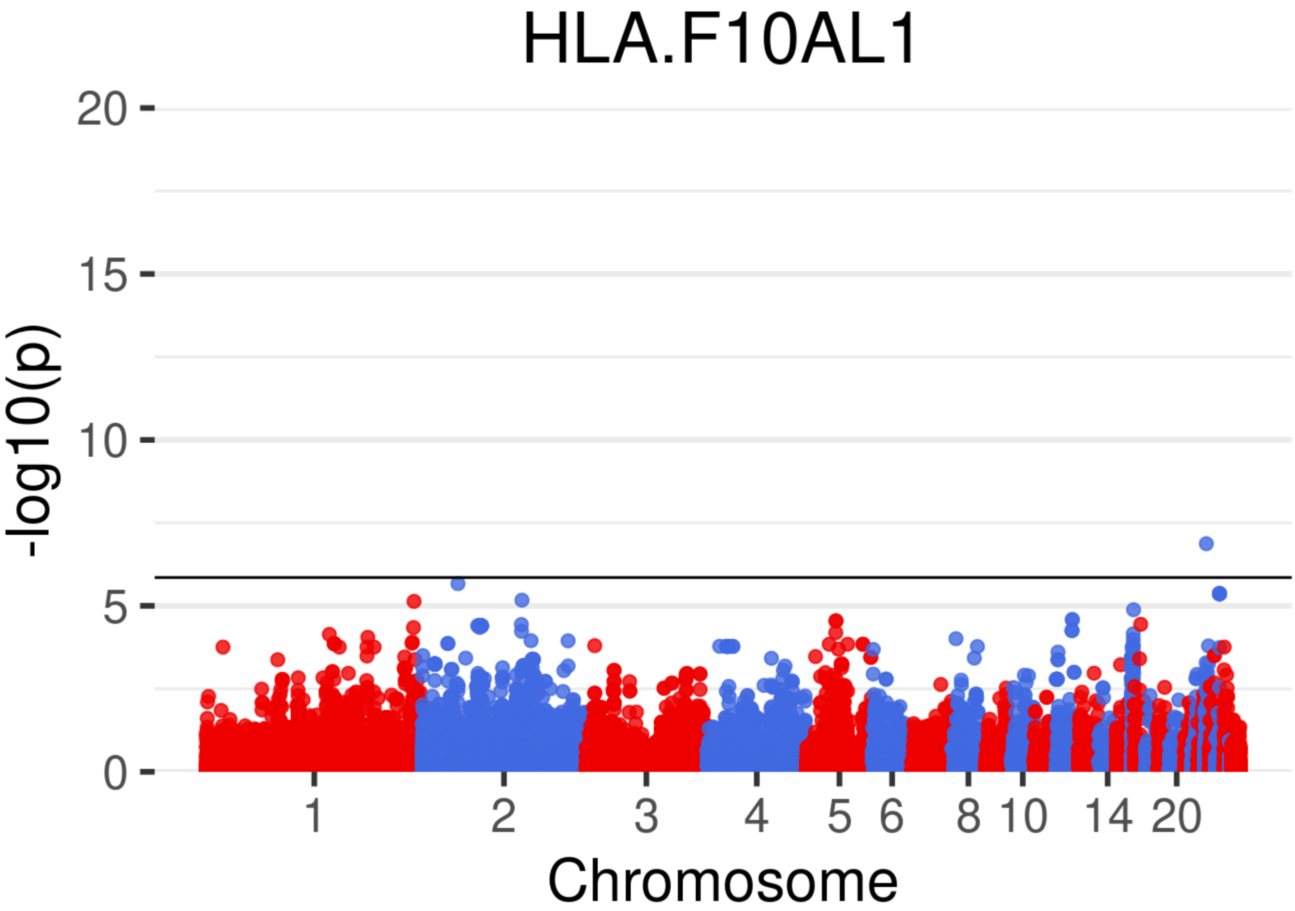
HLA.F10AL1
20
15
10
5
0
-log10(p)
1
2
3
4
5
6
8
10
14
20
Chromosome

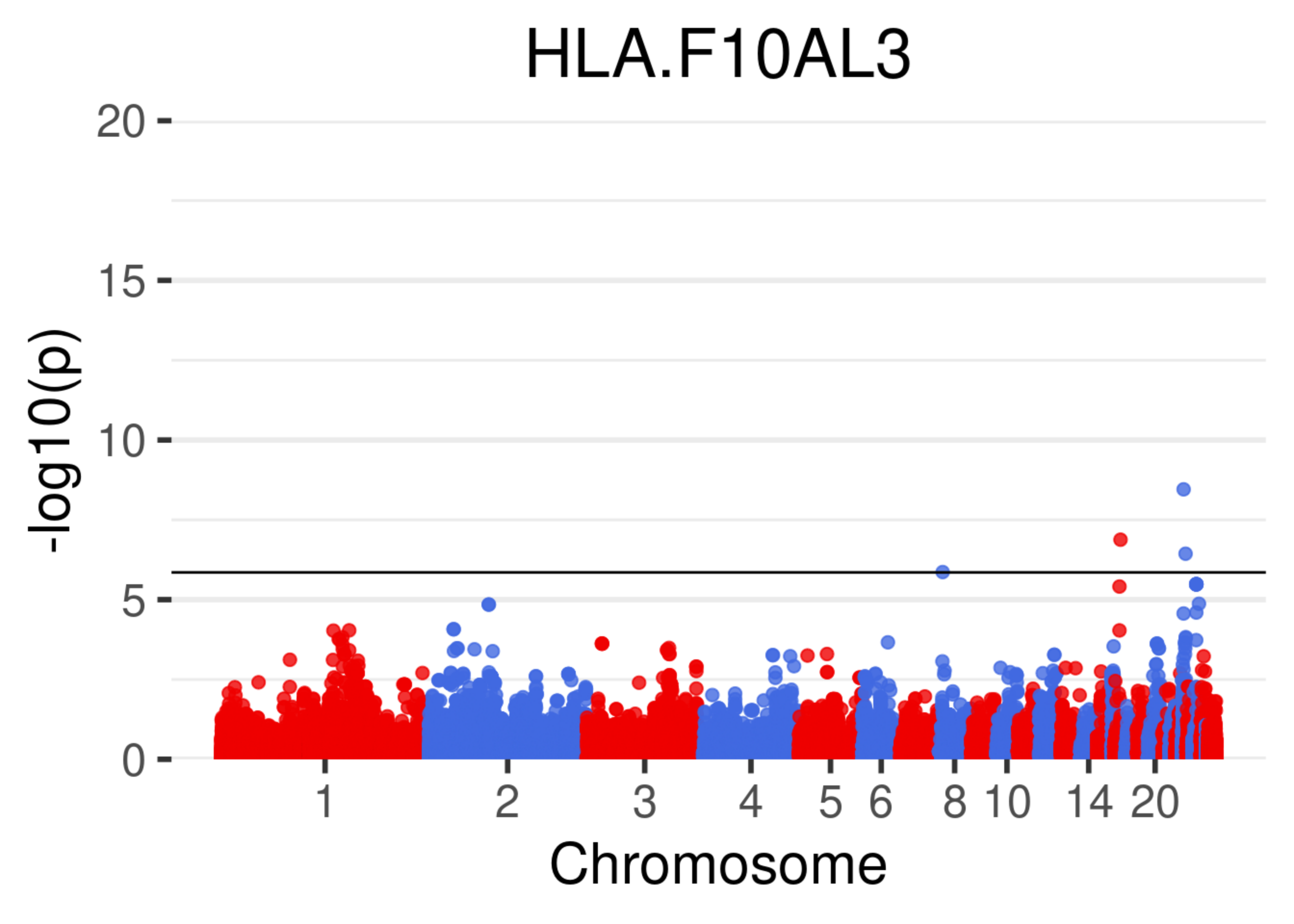

HLA.F10AL3
-log10(p)
0
5
10
15
20
1
2
3
4
5
6
8
10
14
20
Chromosome

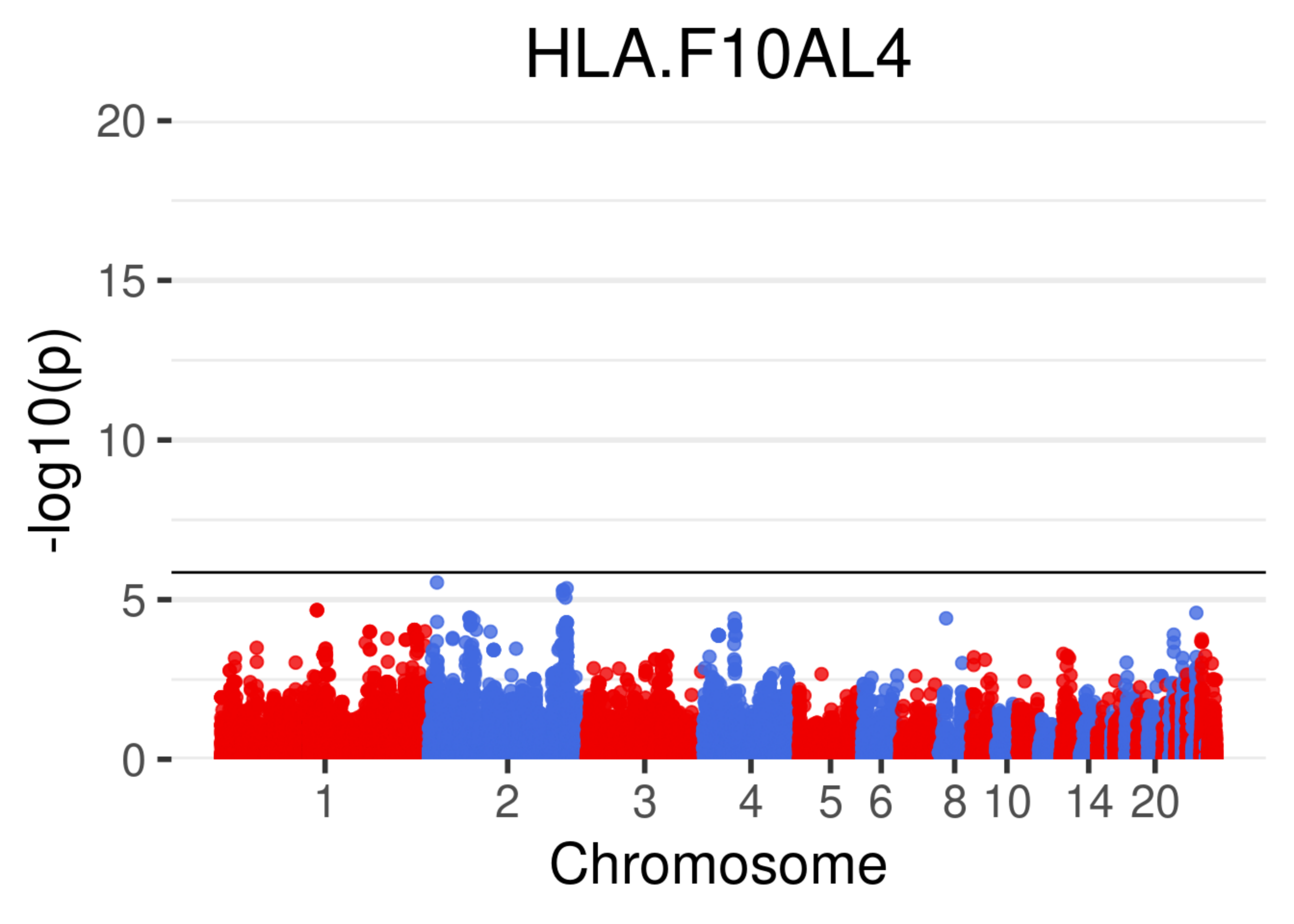

HLA.F10AL4
-log10(p)
20
15
10
5
0
1
2
3
4
5
6
8
10
14
20
Chromosome

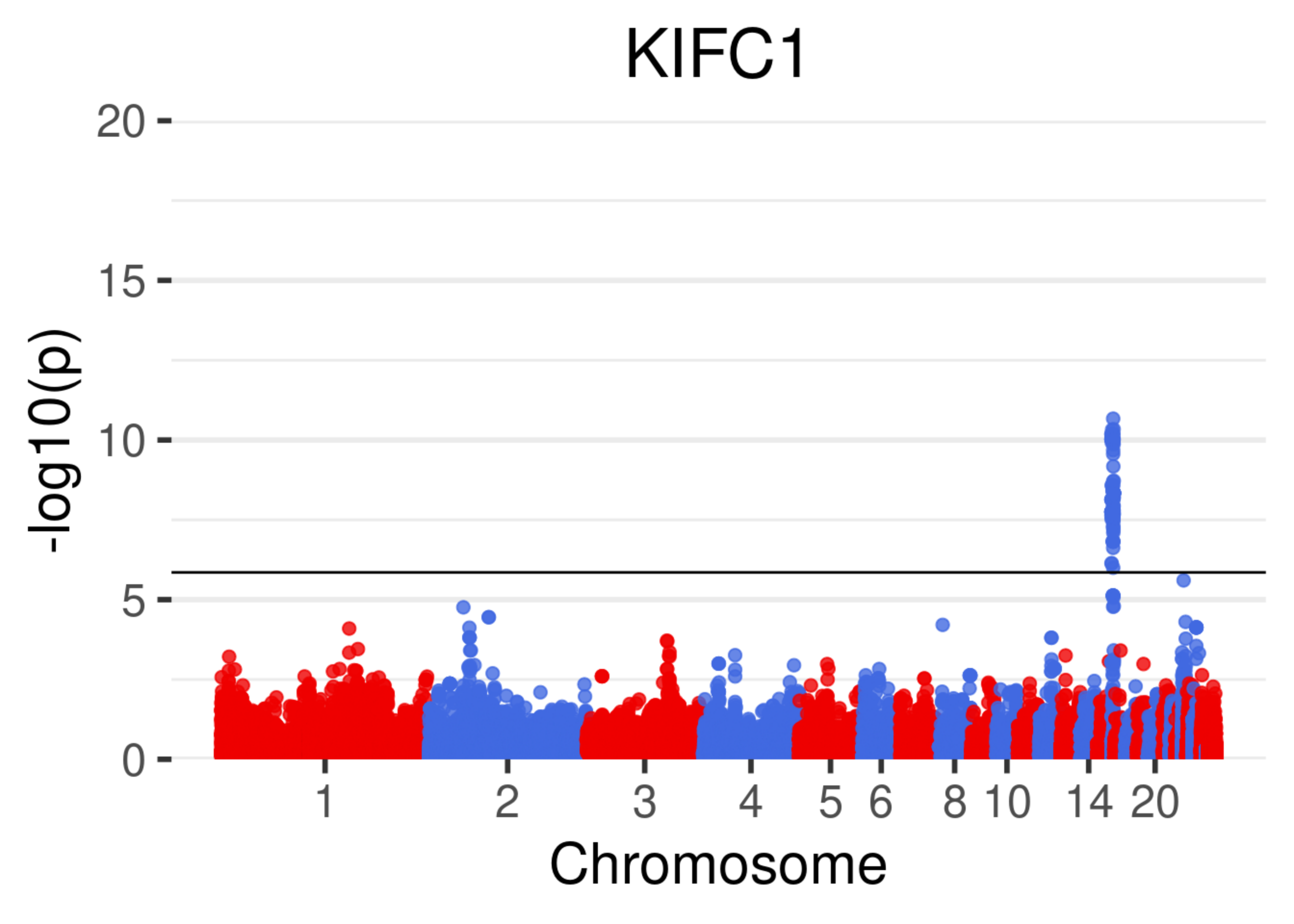

KIFC1
20
15
10
5
0
-log10(p)
1
2
3
4
5
6
8
10
14
20
Chromosome

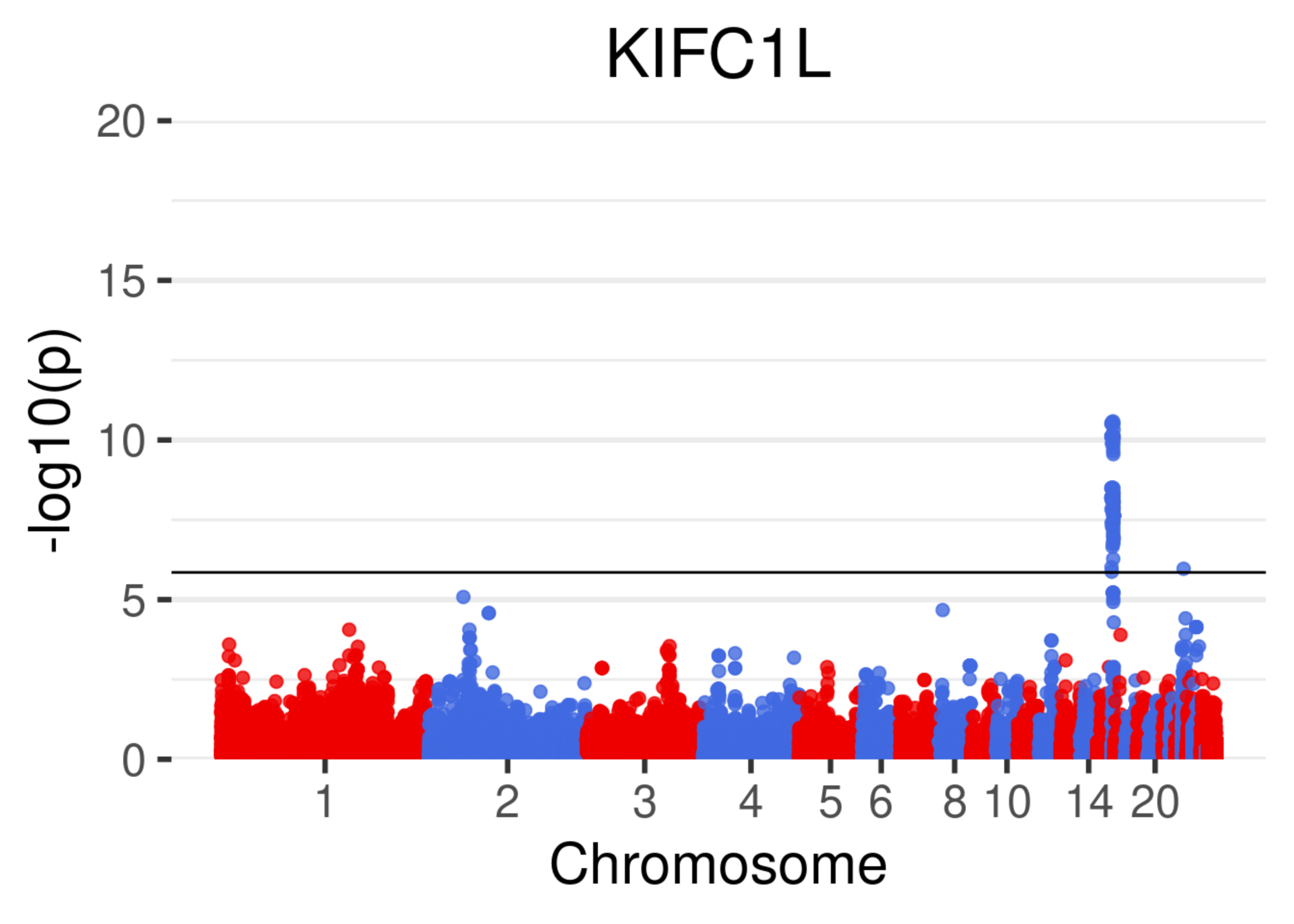
KIFC1L
-log10(p)
20
15
10
5
0
1
2
3
4
5
6
8
10
14
20
Chromosome

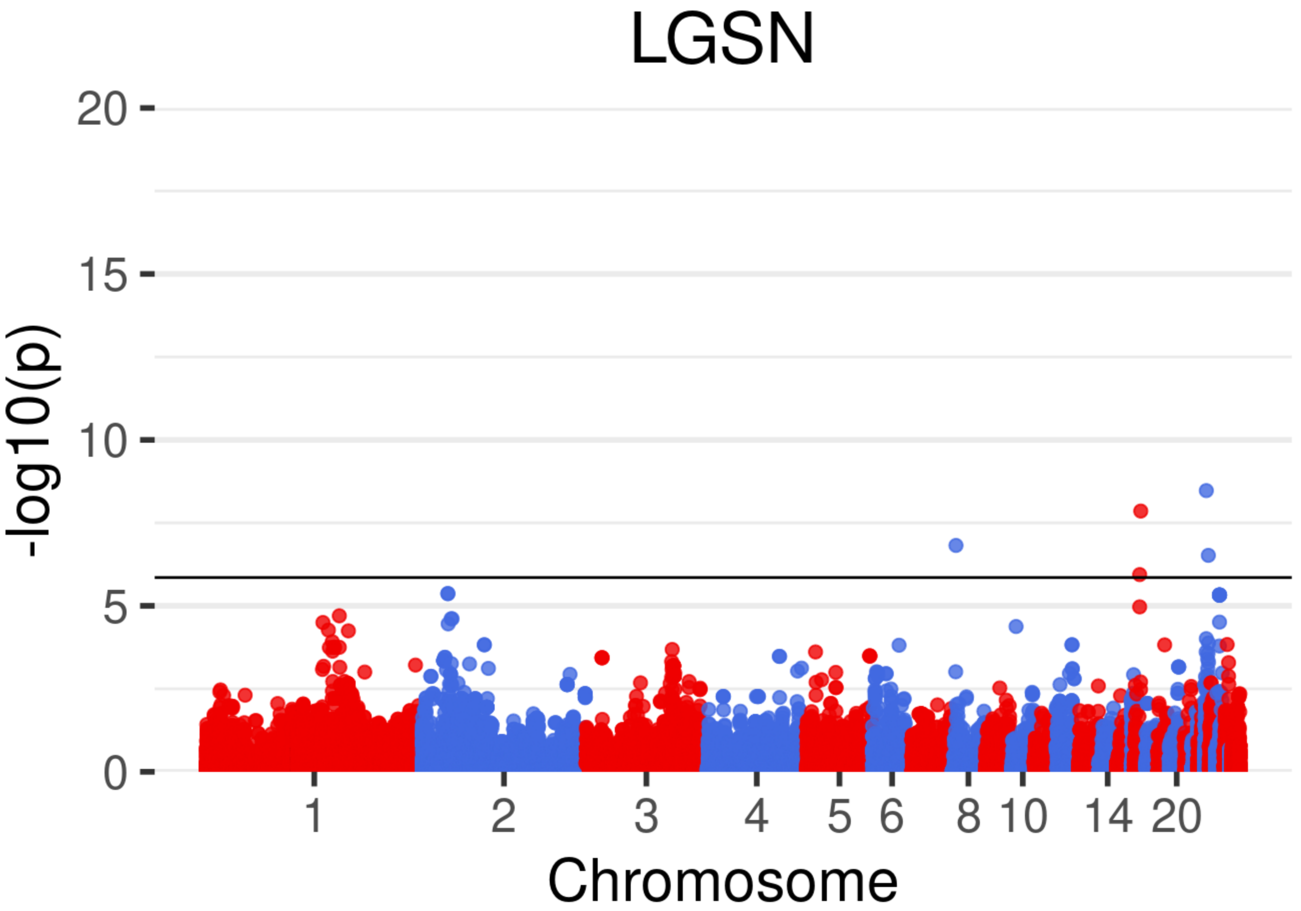
LGSN
20
15
10
5
0
-log10(p)
1
2
3
4
5
6
8
10
14
20
Chromosome

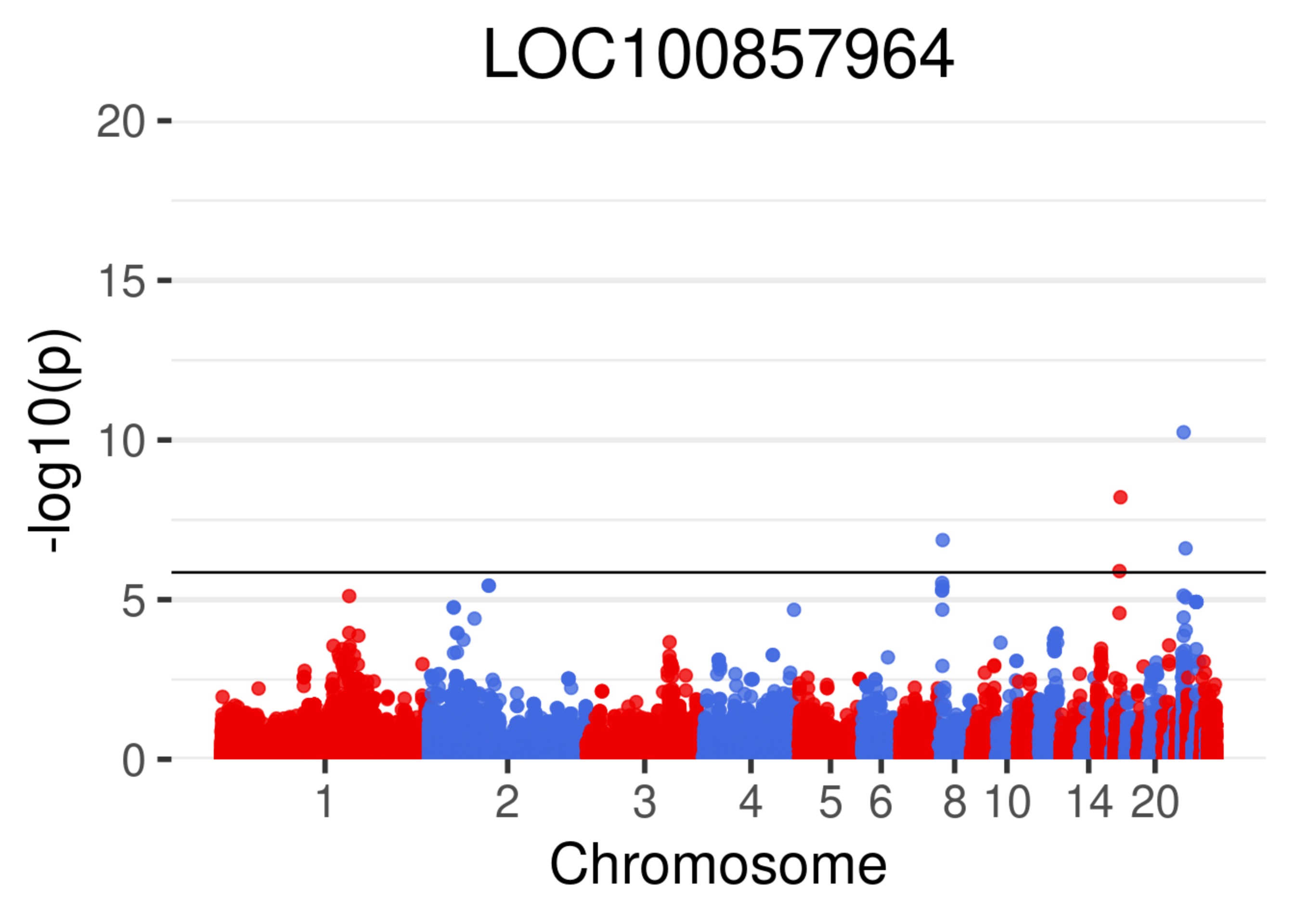
LOC100857964
-log10(p)
20
15
10
5
0
1
2
3
4
5
6
8
10
14
20
Chromosome

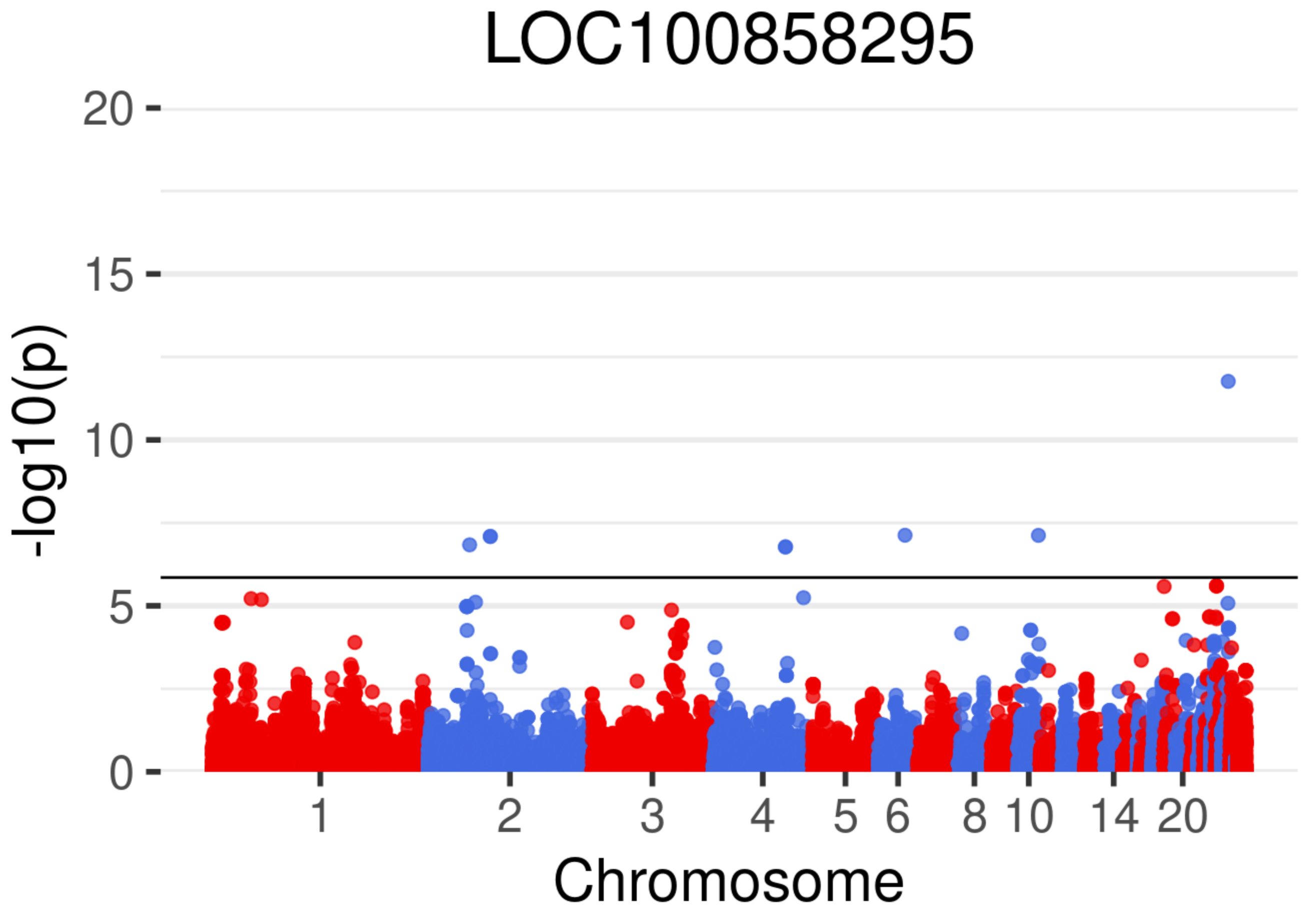

LOC100858295
-log10(p)
0
5
10
15
20
1
2
3
4
5
6
8
10
14
20
Chromosome

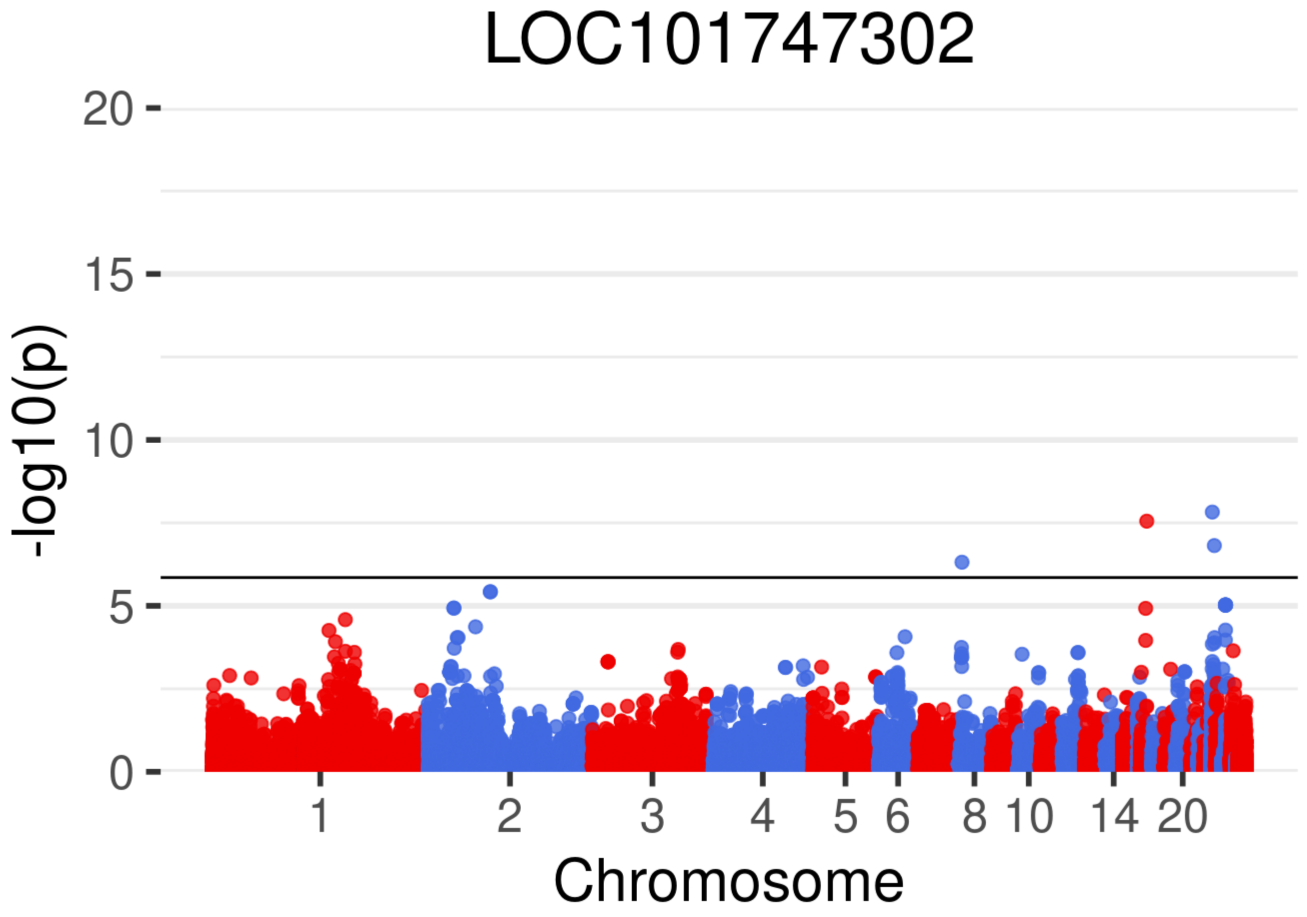
LOC101747302
-log10(p)
20
15
10
5
0
1
2
3
4
5
6
8
10
14
20
Chromosome

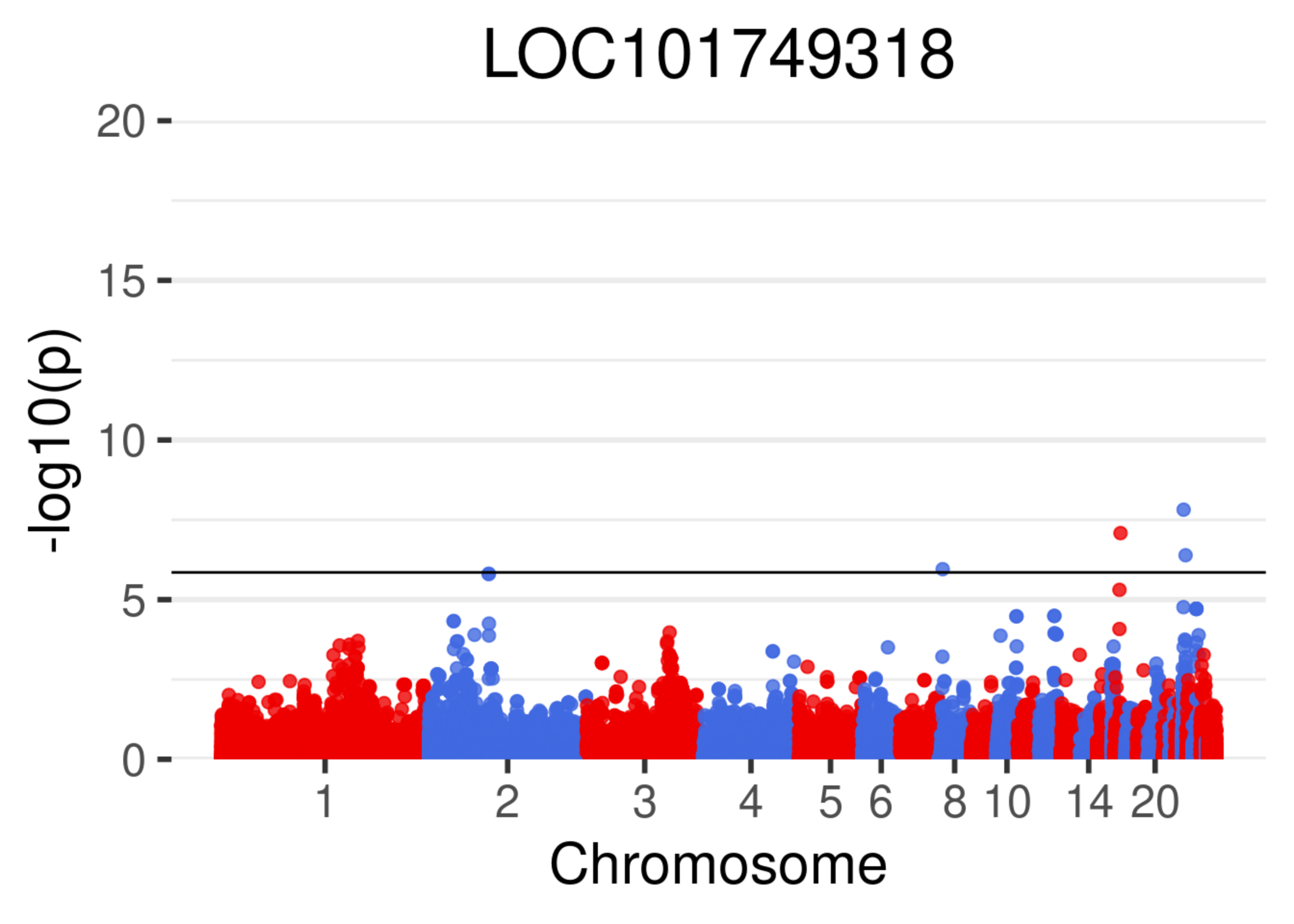
LOC101749318
-log10(p)
20
15
10
5
0
1
2
3
4
5
6
8
10
14
20
Chromosome

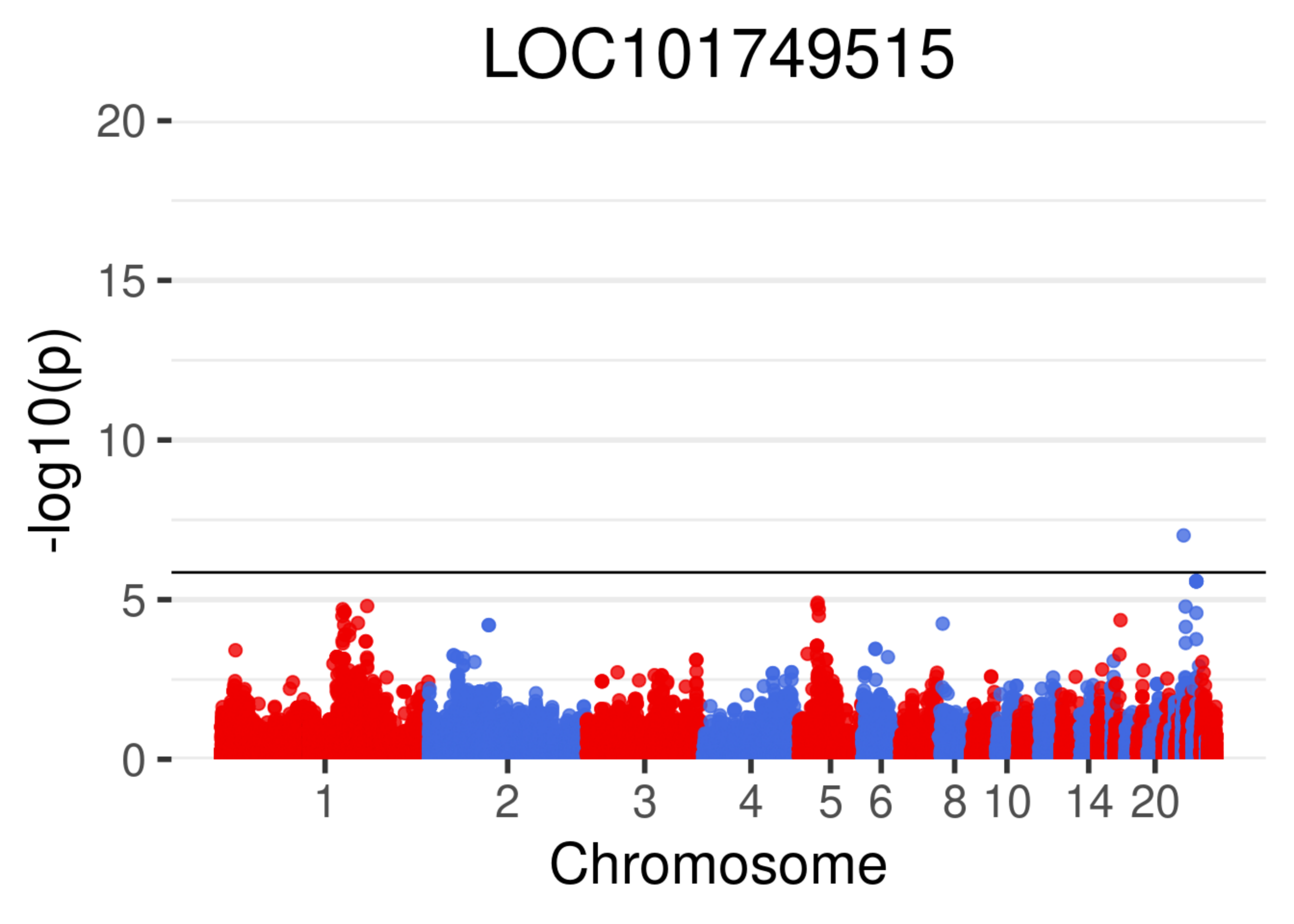
LOC101749515
20
15
10
5
0
-log10(p)
1
2
3
4
5
6
8
10
14
20
Chromosome

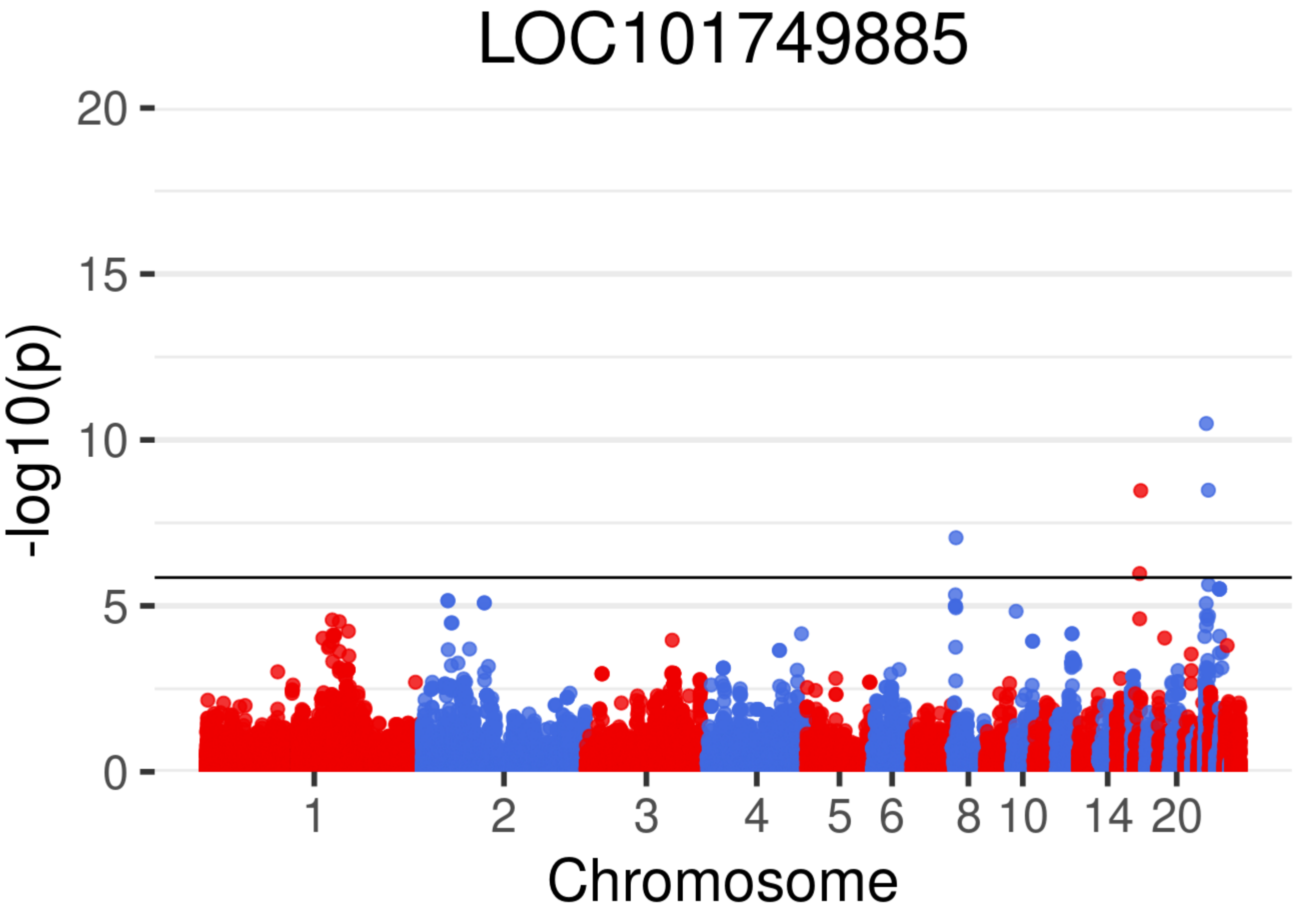
LOC101749885
-log10(p)
20
15
10
5
0
1
2
3
4
5
6
8
10
14
20
Chromosome

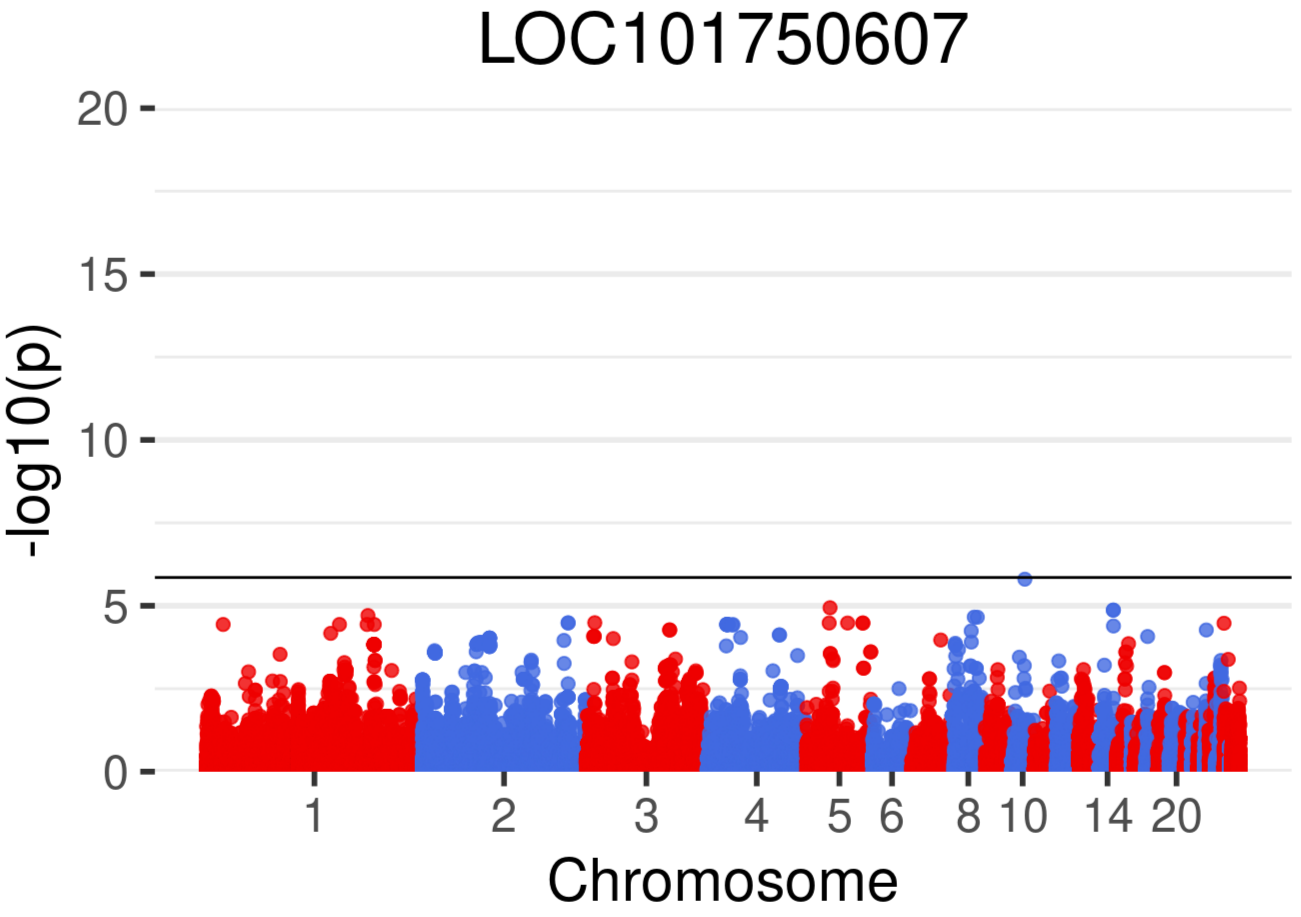

LOC101750607
-log10(p)
20
15
10
5
0
1
2
3
4
5
6
8
10
14
20
Chromosome

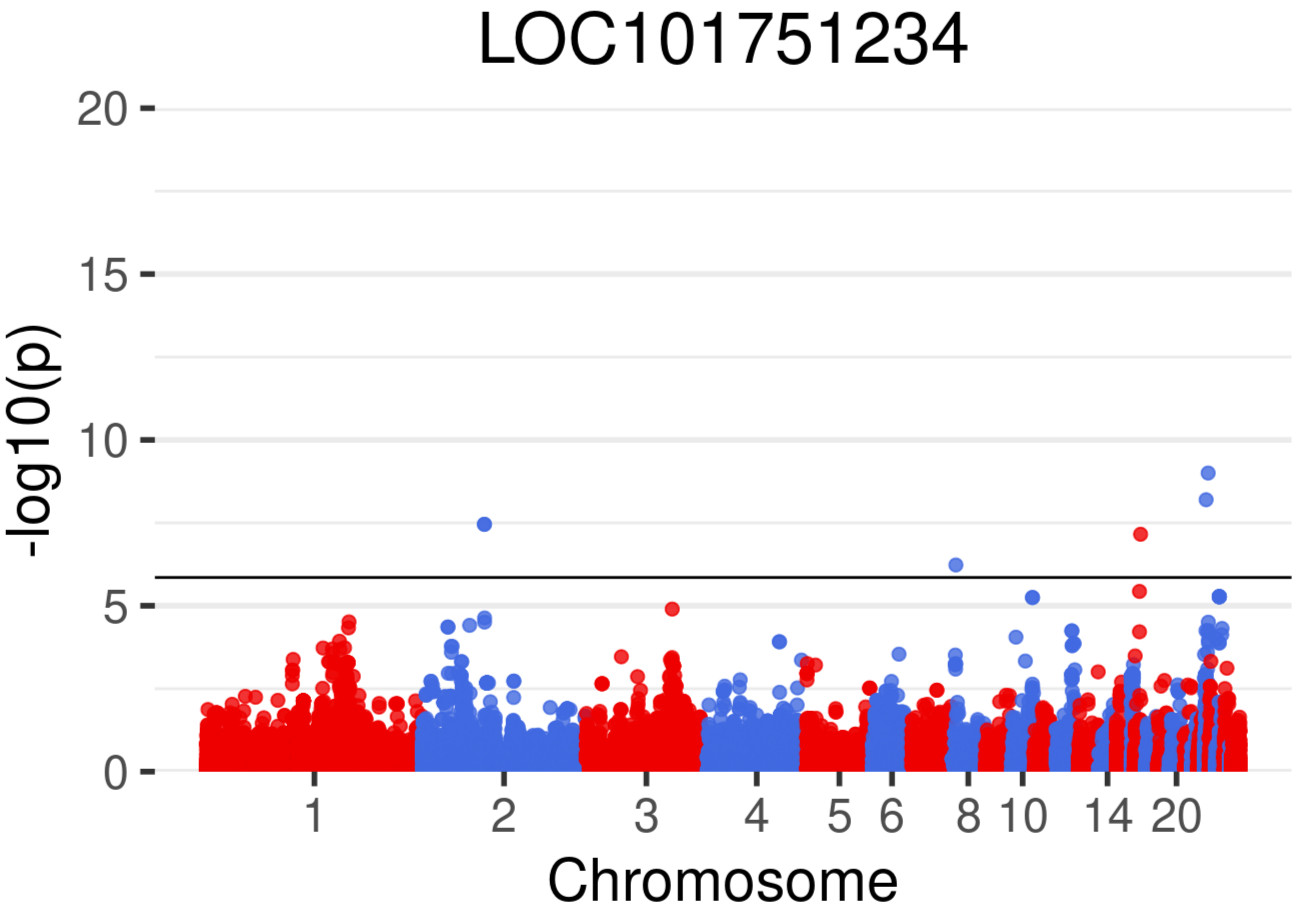
LOC101751234
-log10(p)
0
5
10
15
20
1
2
3
4
5
6
8
10
14
20
Chromosome

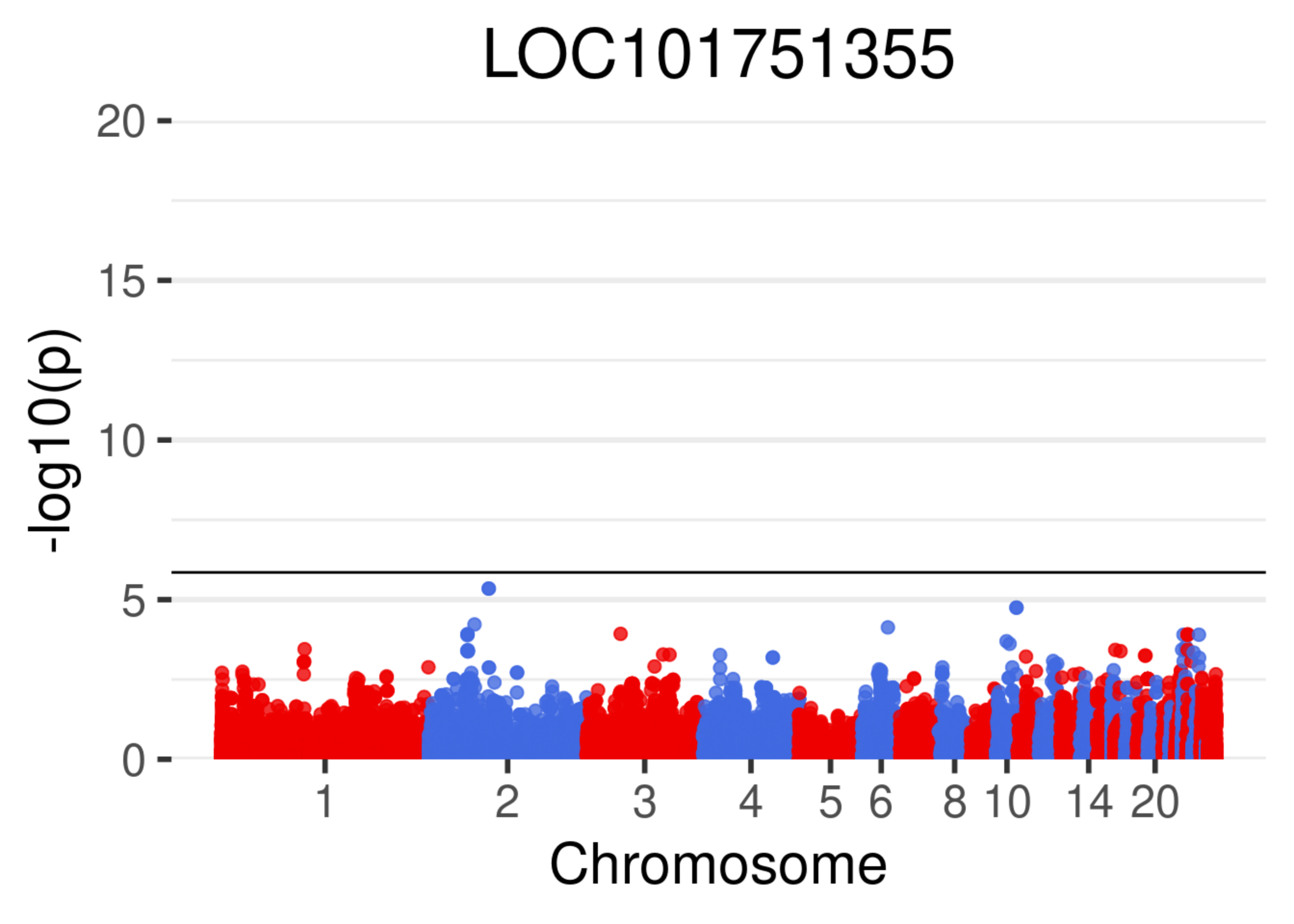

LOC101751355
-log10(p)
20
15
10
5
0
1
2
3
4
5
6
8
10
14
20
Chromosome

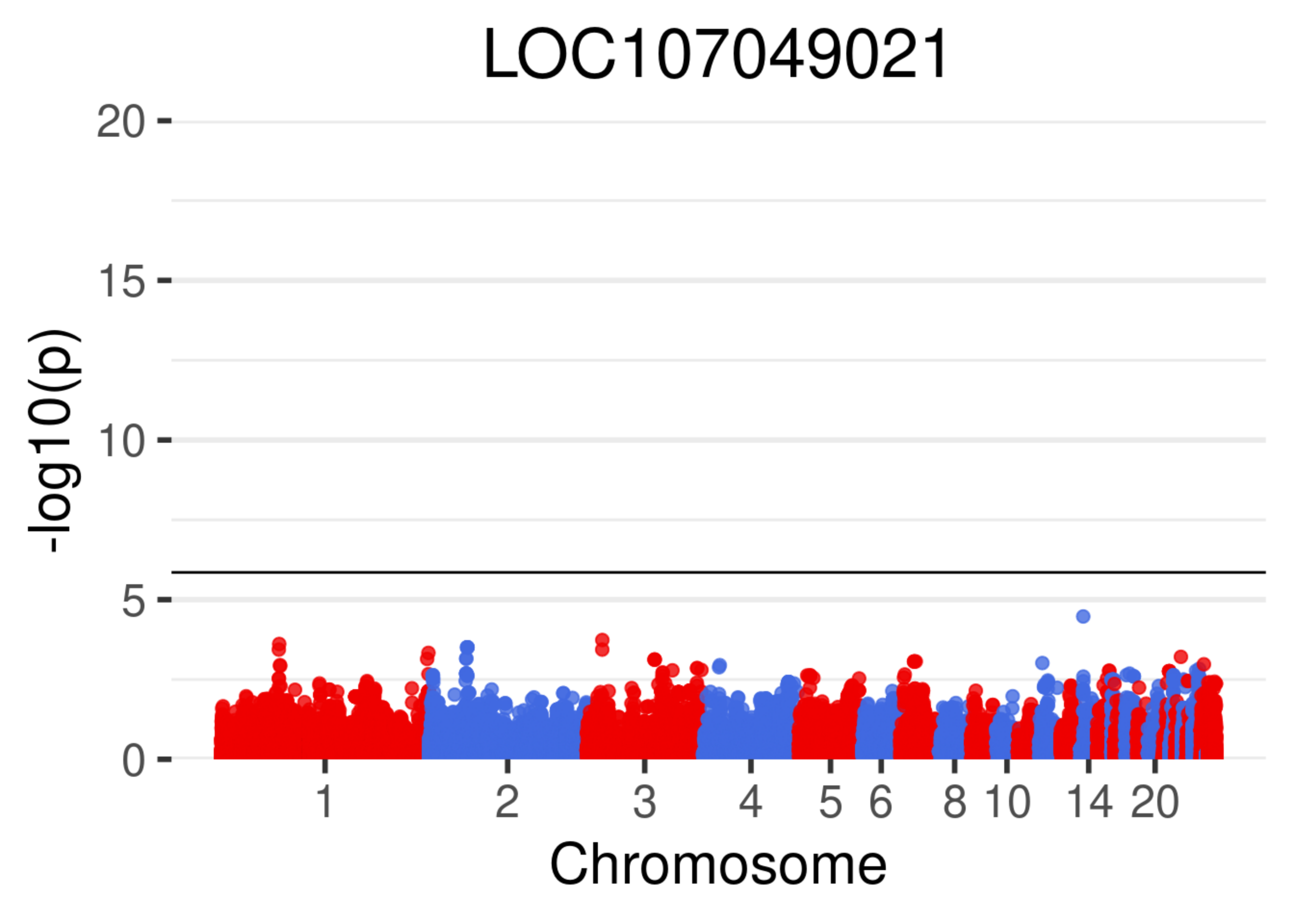

LOC107049021
-log10(p)
20
15
10
5
0
1
2
3
4
5
6
8
10
14
20
Chromosome

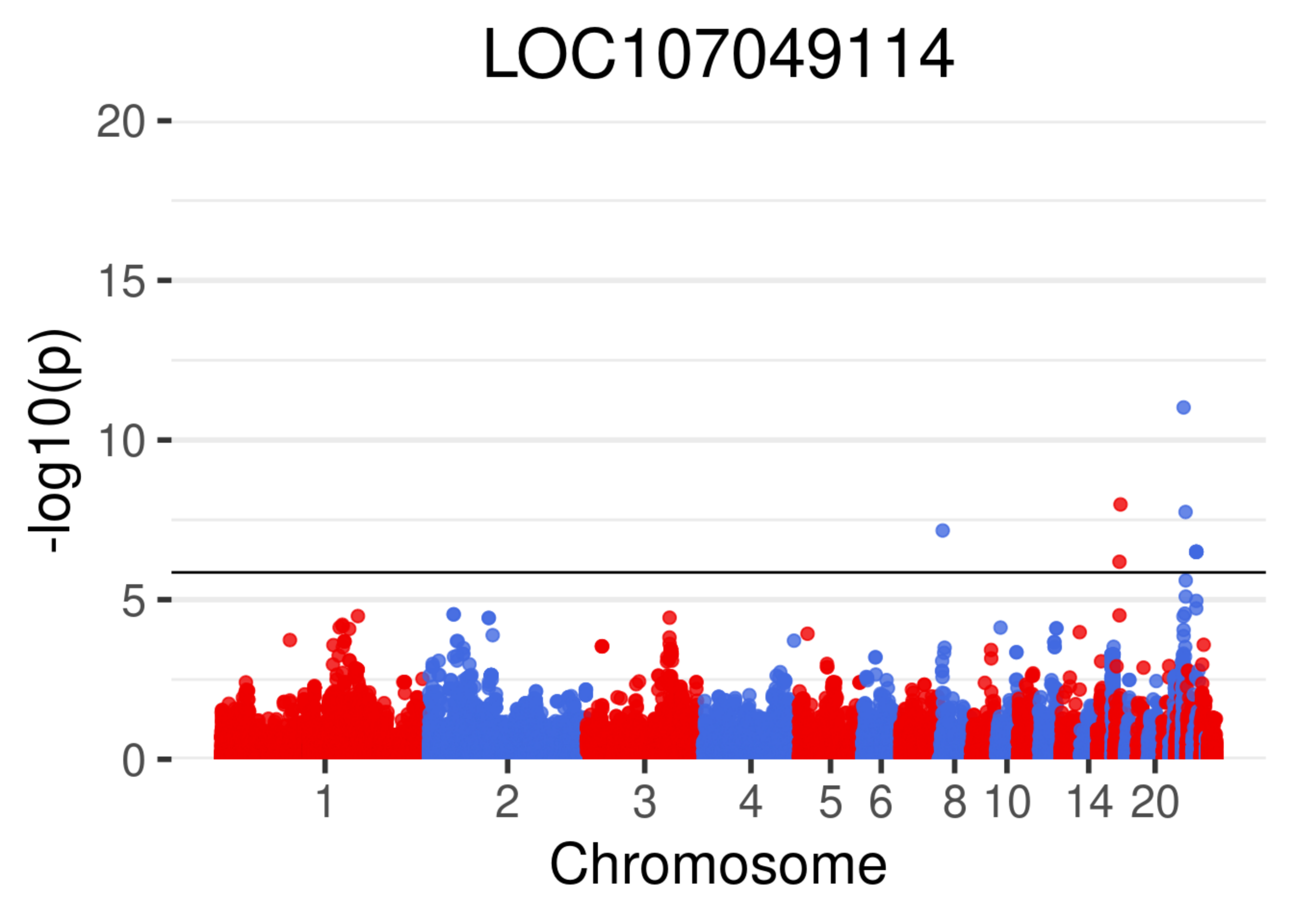
LOC107049114
-log10(p)
20
15
10
5
0
1
2
3
4
5
6
8
10
14
20
Chromosome

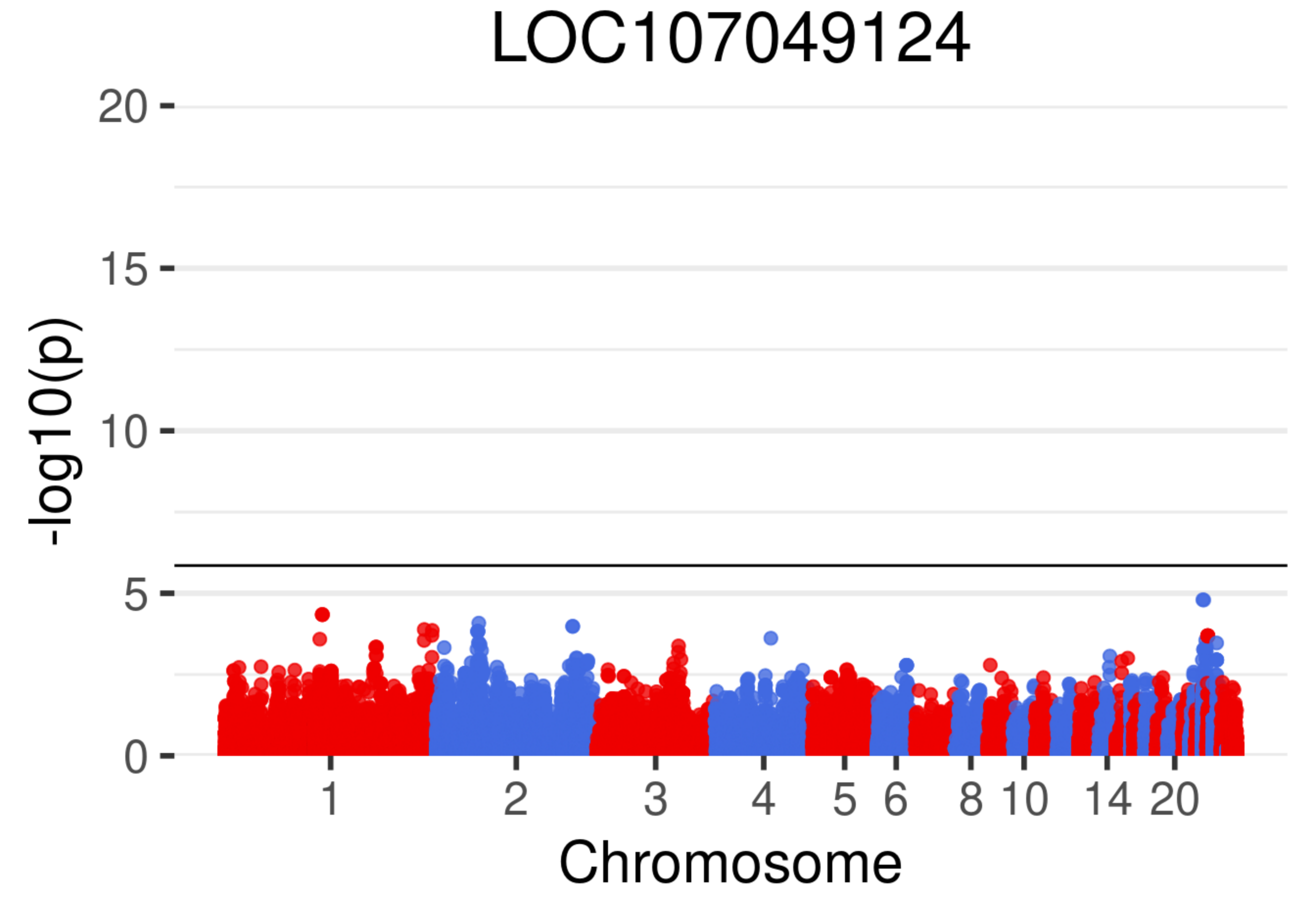

LOC107049124
-log10(p)
20
15
10
5
0
1
2
3
4
5
6
8
10
14
20
Chromosome

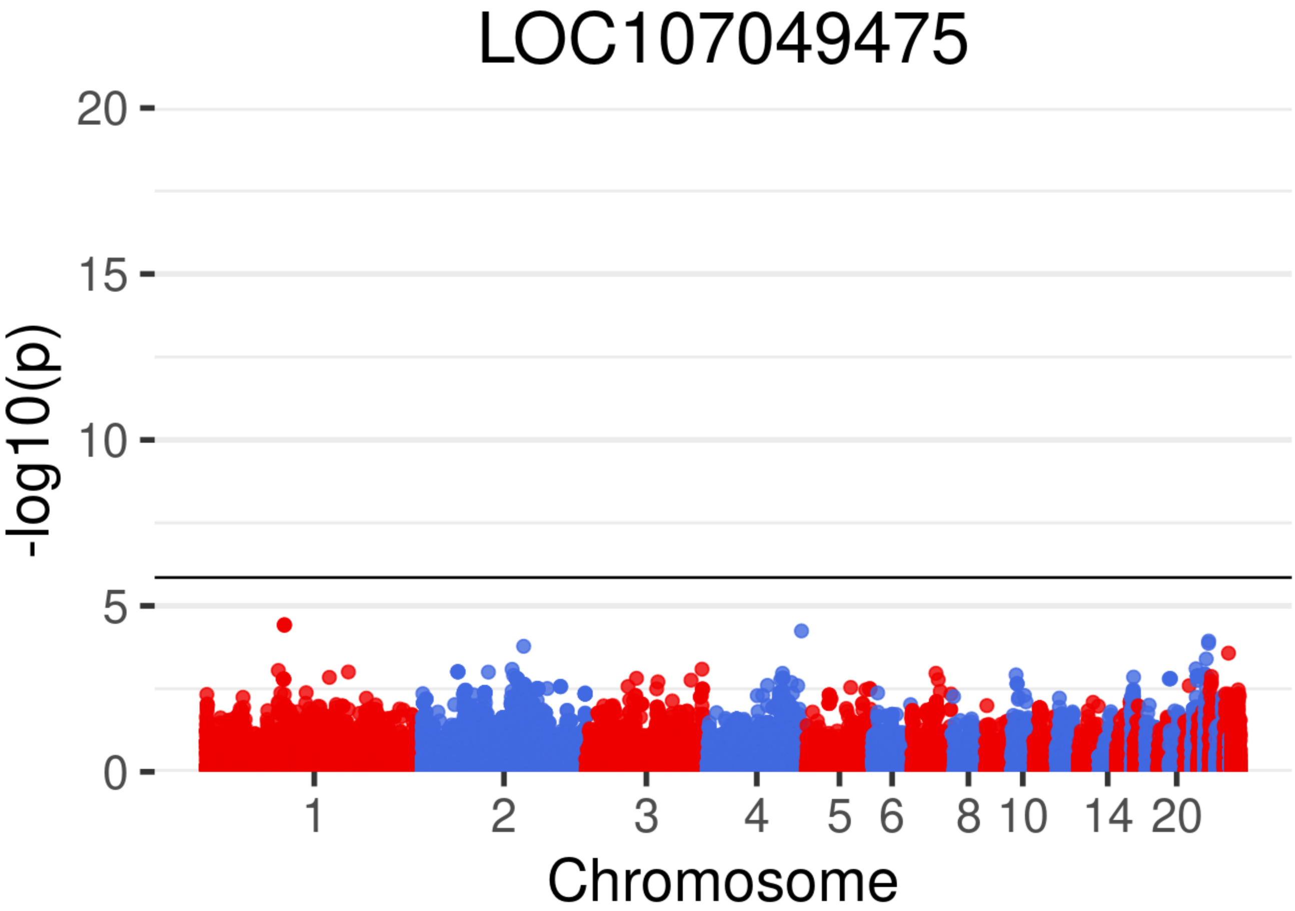

LOC107049475
-log10(p)
20
15
10
5
0
1
2
3
4
5
6
8
10
14
20
Chromosome

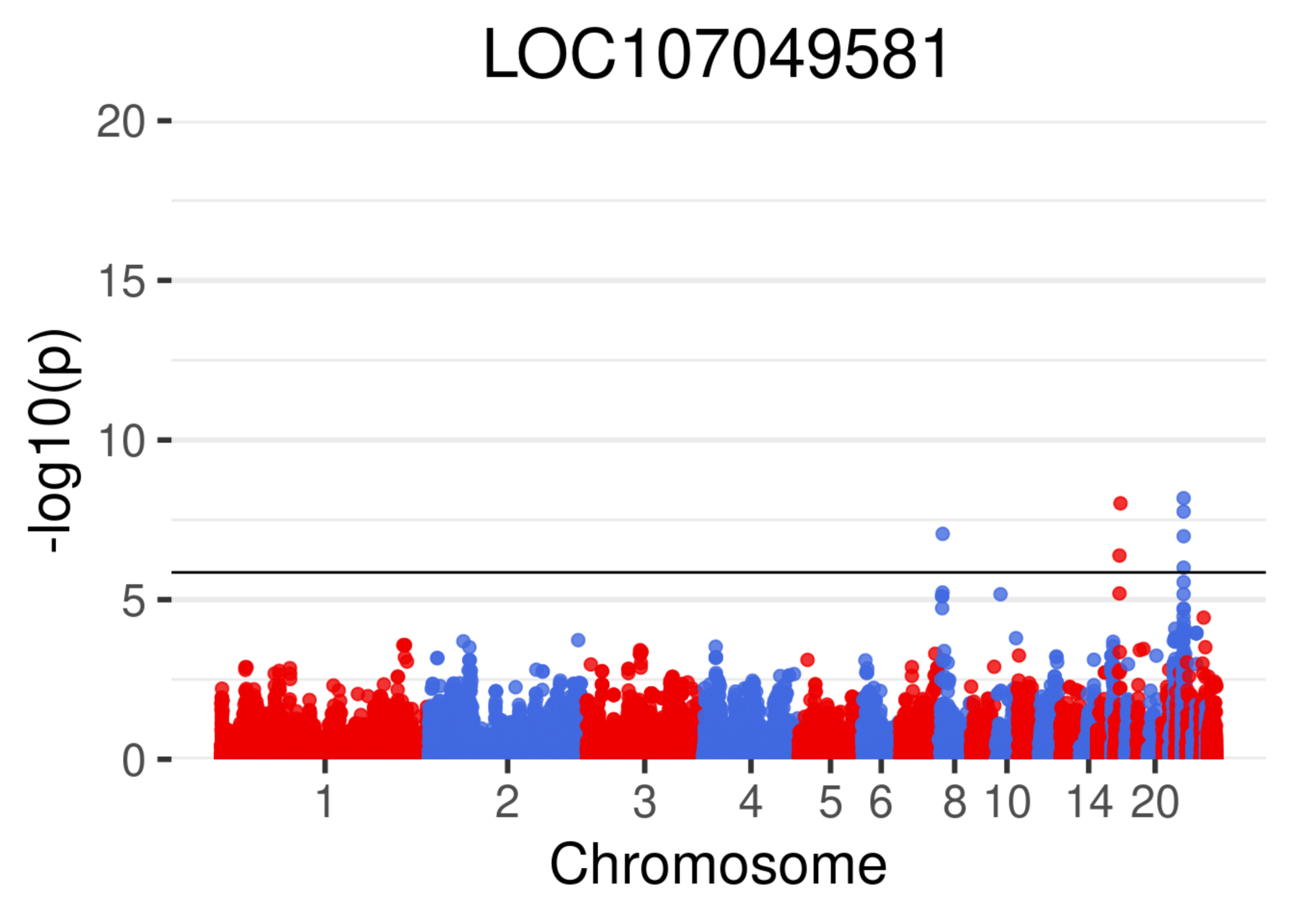
LOC107049581
20
15
10
5
0
-log10(p)
1
2
3
4
5
6
8
10
14
20
Chromosome

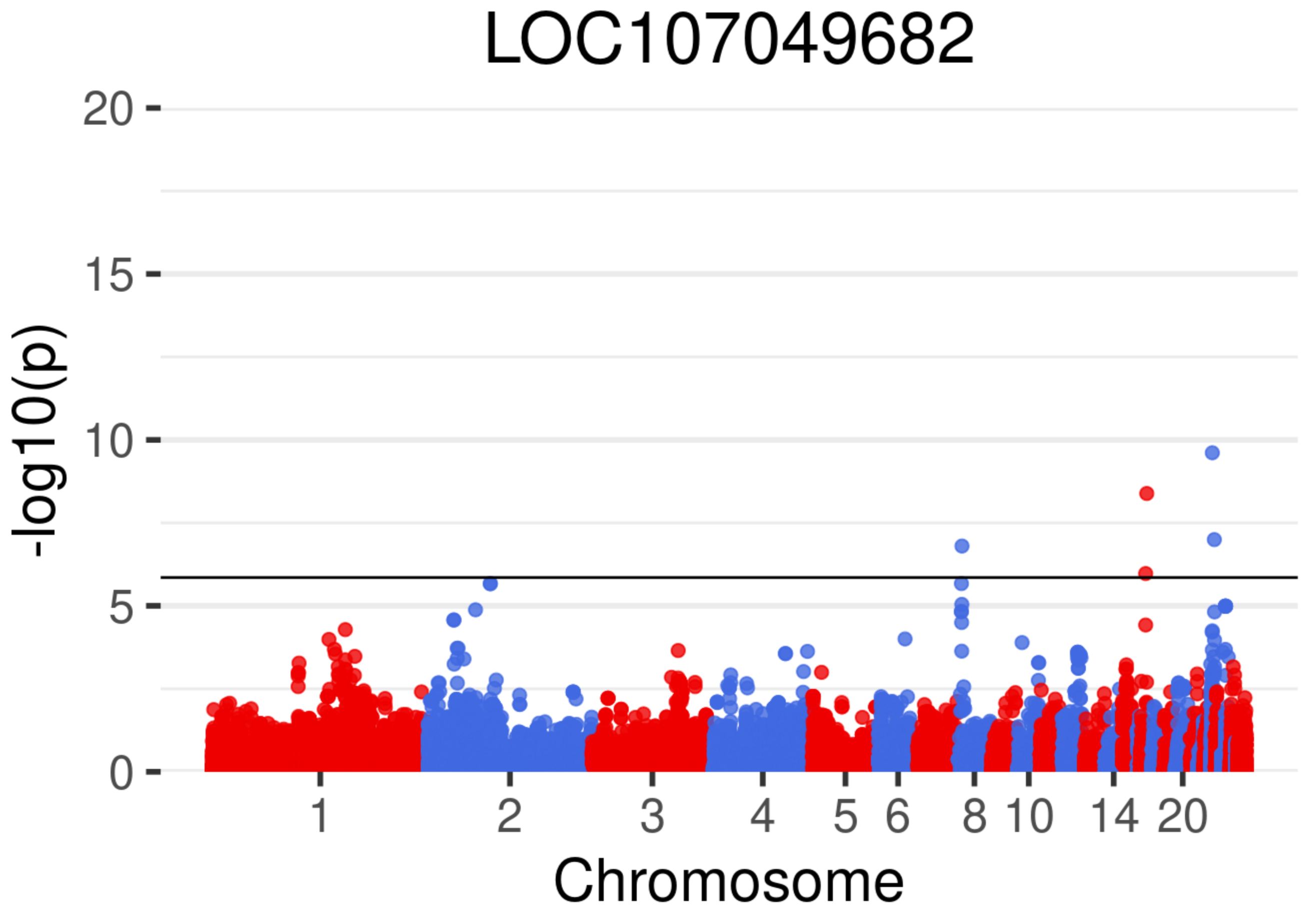
LOC107049682
-log10(p)
0
5
10
15
20
1
2
3
4
5
6
8
10
14
20
Chromosome

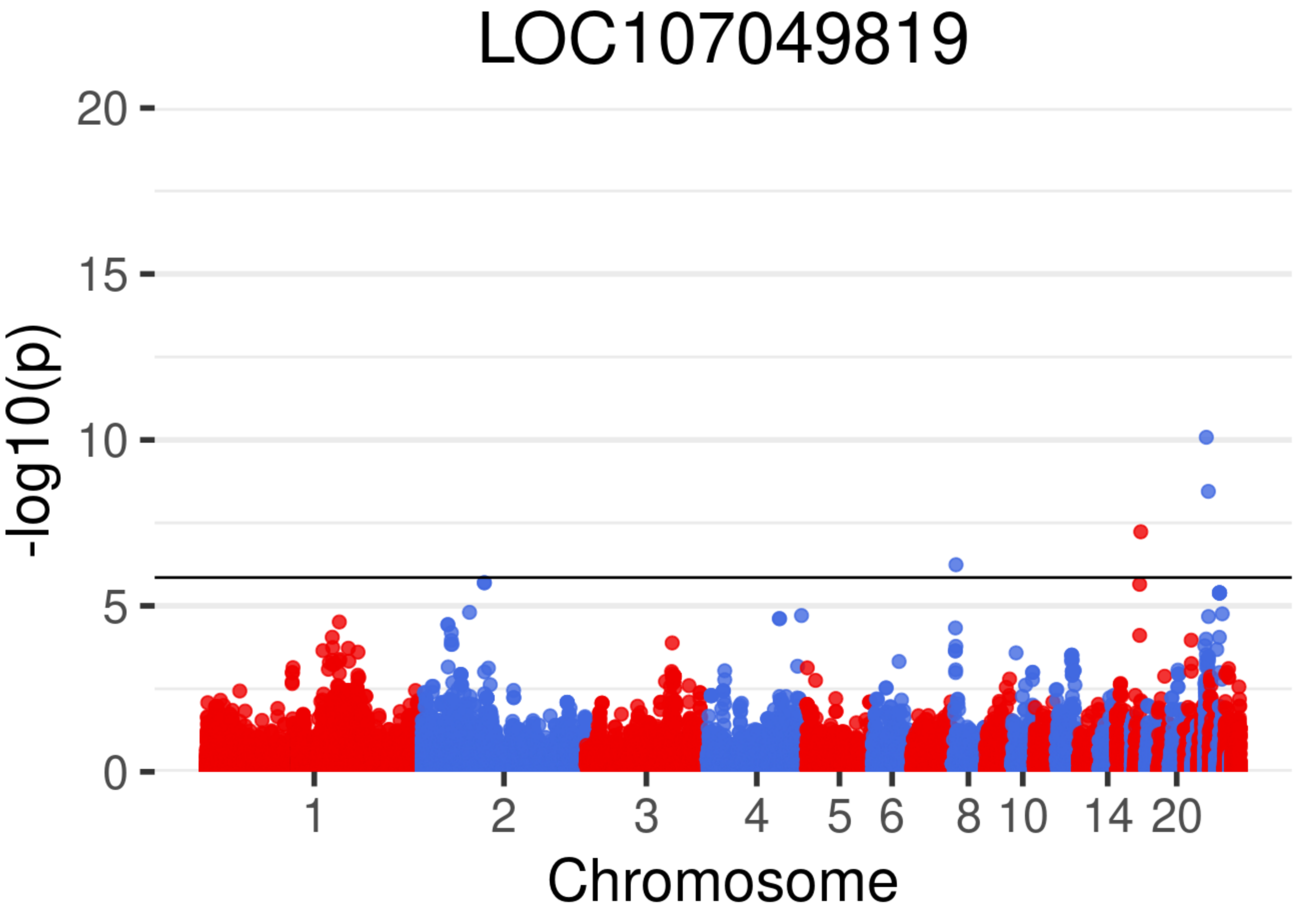

LOC107049819
-log10(p)
0
5
10
15
20
1
2
3
4
5
6
8
10
14
20
Chromosome

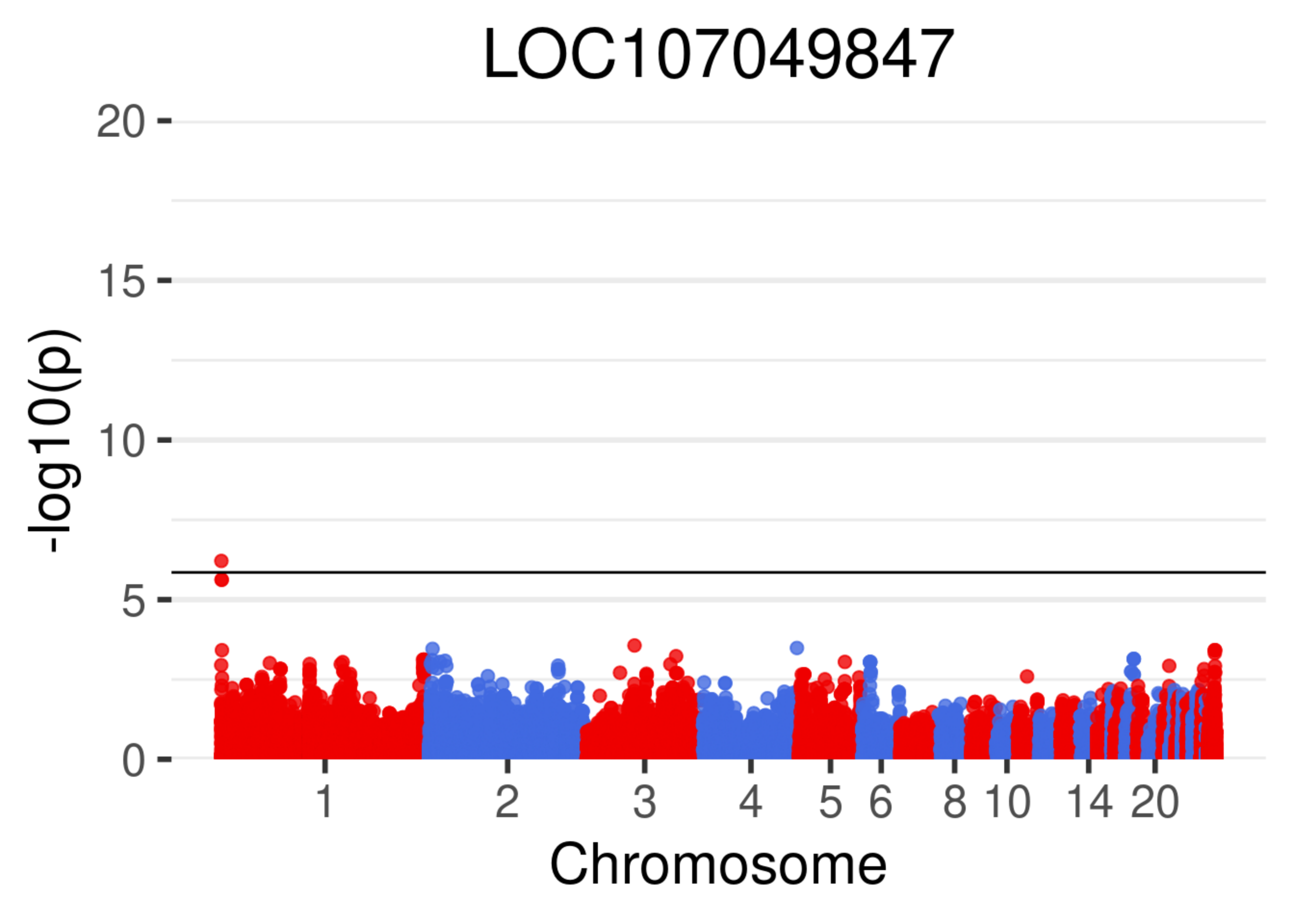

LOC107049847
20
15
10
5
0
-log10(p)
1
2
3
4
5
6
8
10
14
20
Chromosome

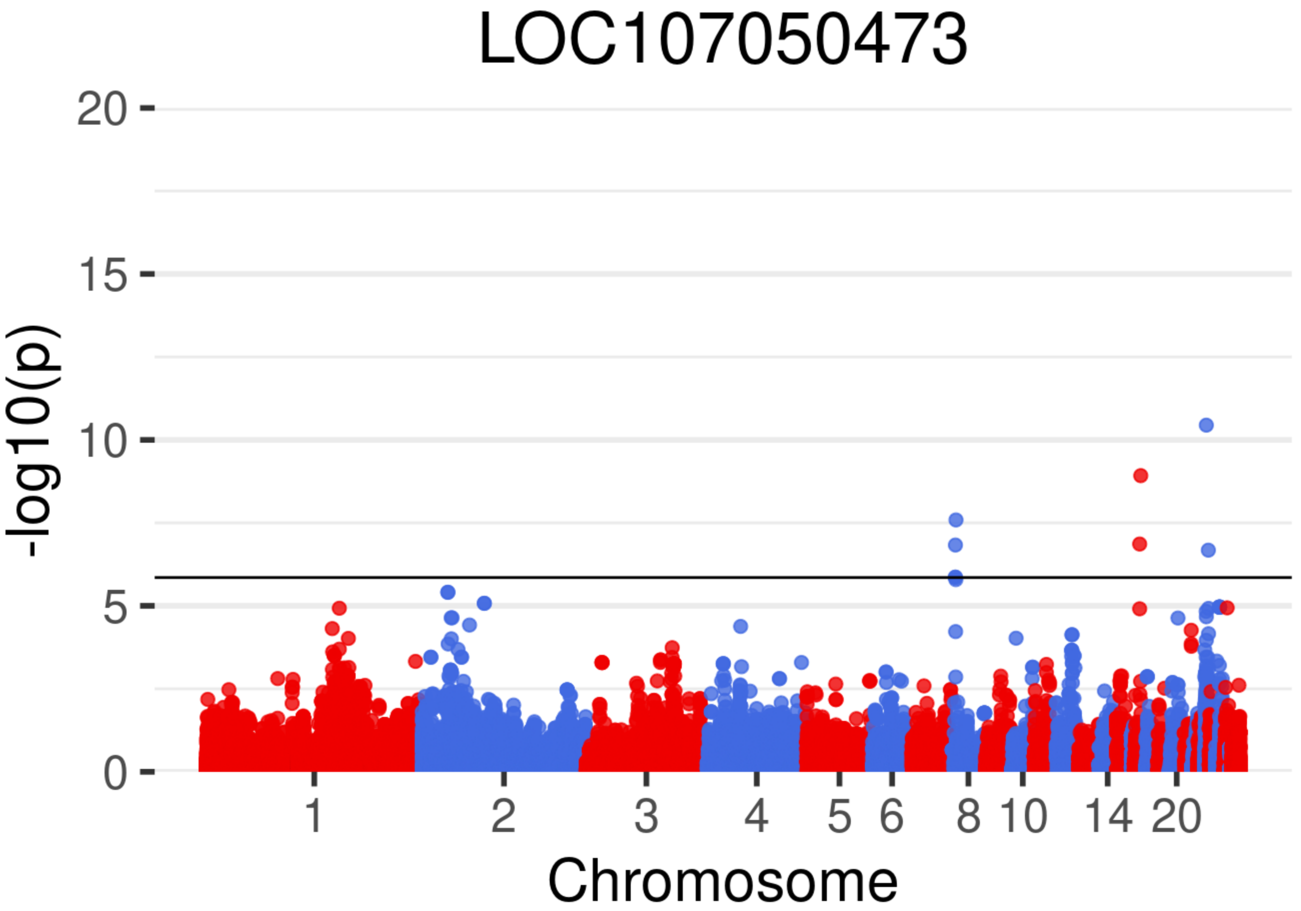
LOC107050473
-log10(p)
0
5
10
15
20
Chromosome
1
2
3
4
5
6
8
10
14
20

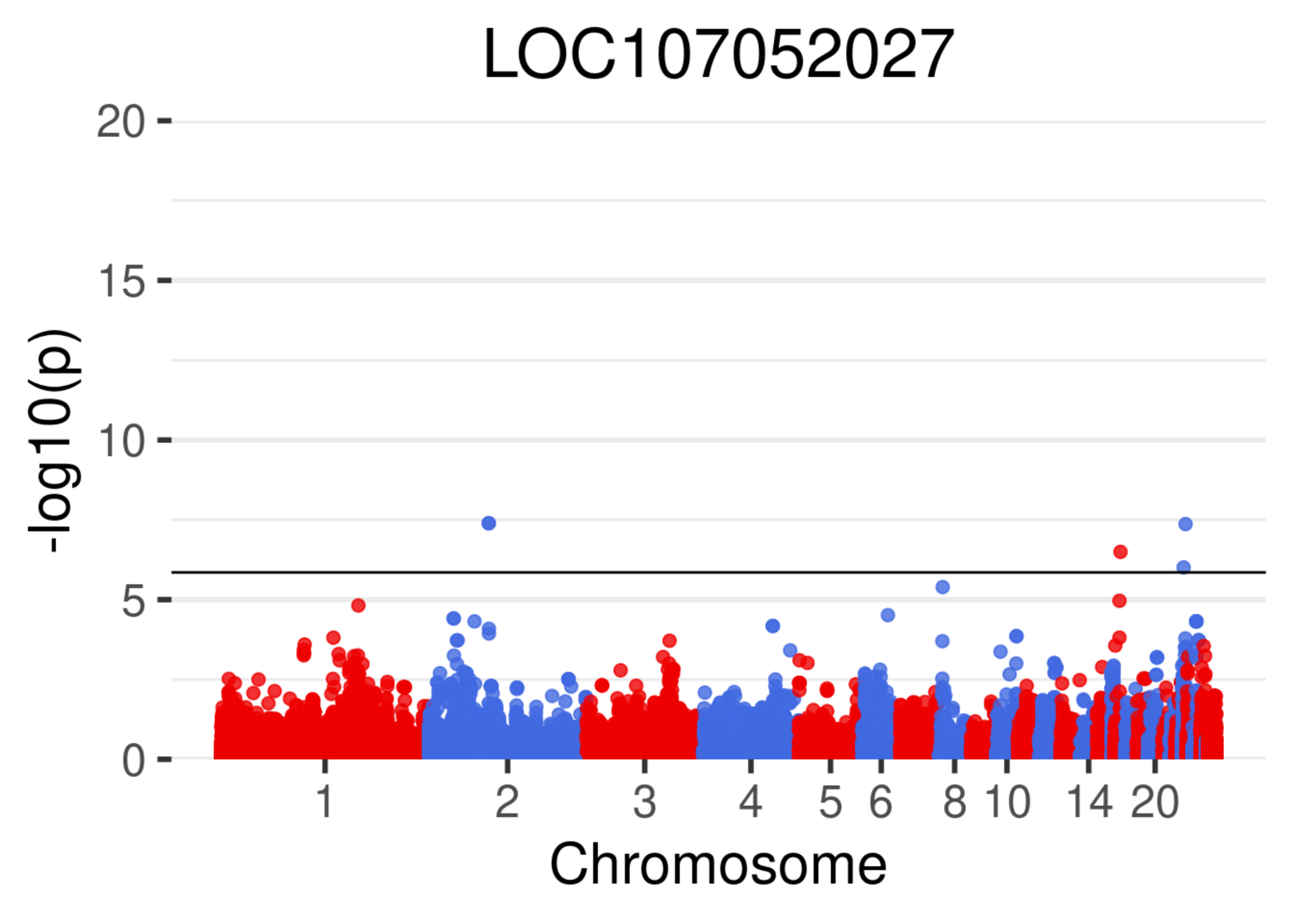
LOC107052027
-log10(p)
20
15
10
5
0
1
2
3
4
5
6
8
10
14
20
Chromosome

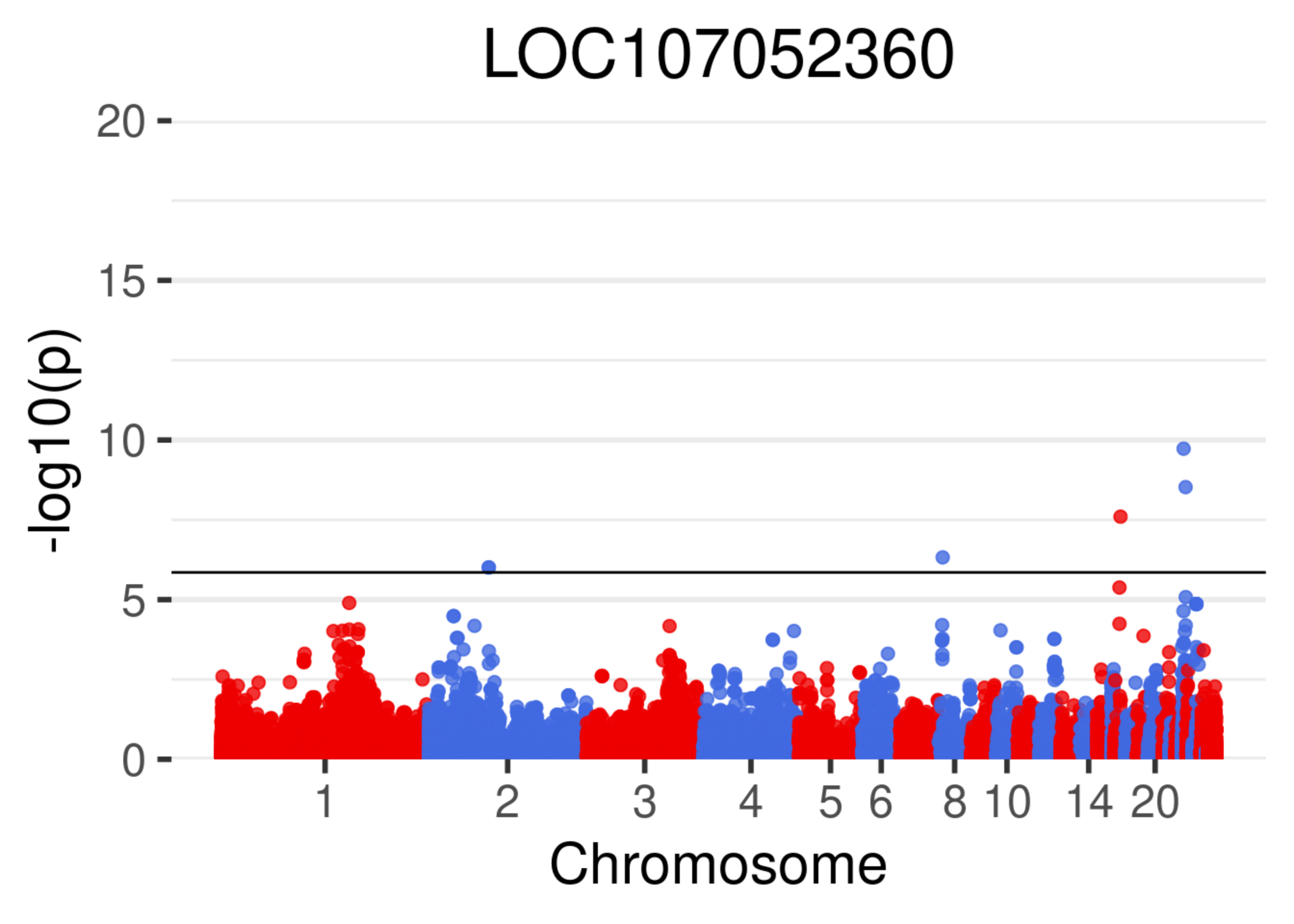
LOC107052360
-log10(p)
0
5
10
15
20
1
2
3
4
5
6
8
10
14
20
Chromosome

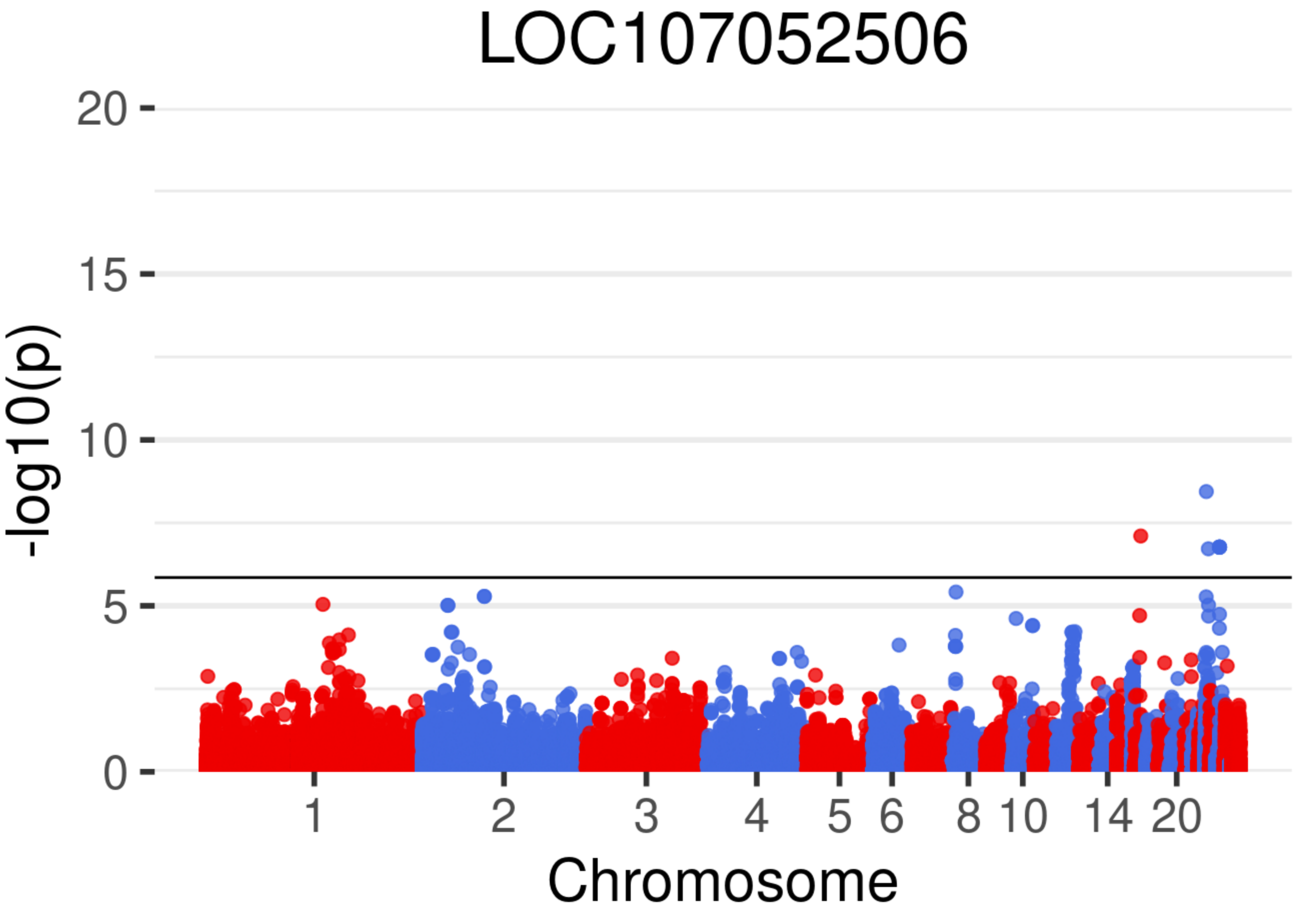
LOC107052506
-log10(p)
20
15
10
5
0
1
2
3
4
5
6
8
10
14
20
Chromosome

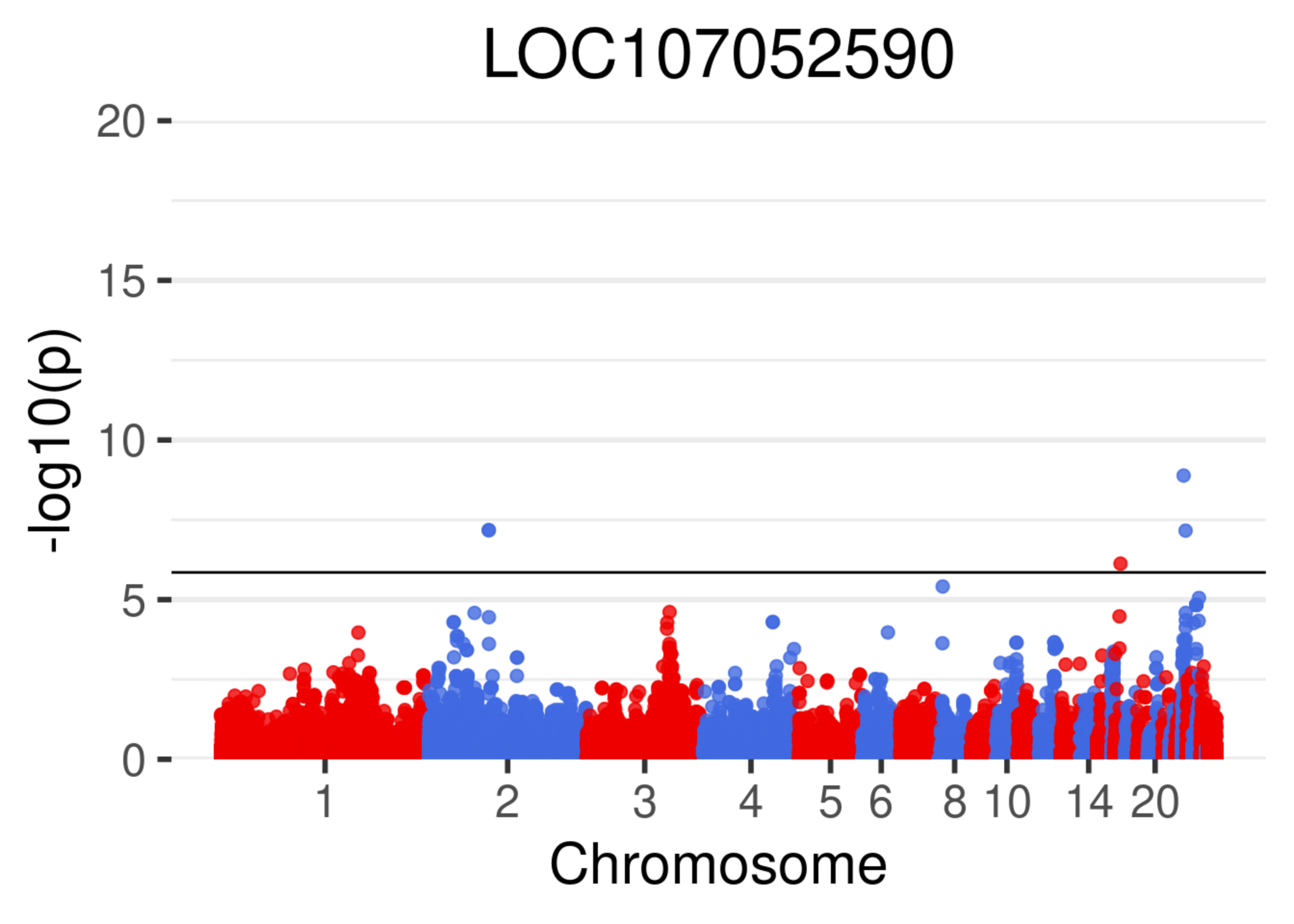
LOC107052590
-log10(p)
20
15
10
5
0
1
2
3
4
5
6
8
10
14
20
Chromosome

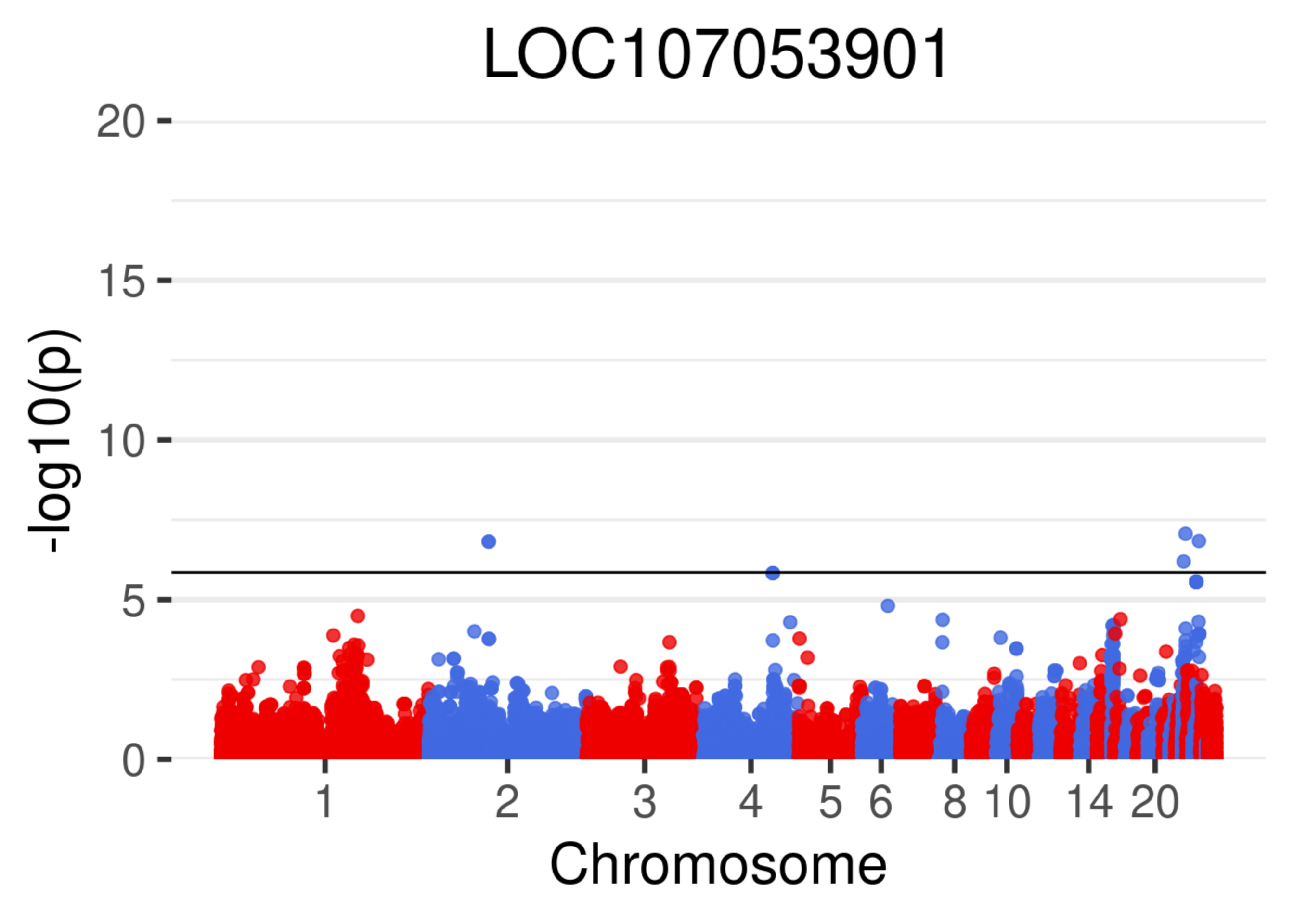
LOC107053901
-log10(p)
20
15
10
5
0
1
2
3
4
5
6
8
10
14
20
Chromosome

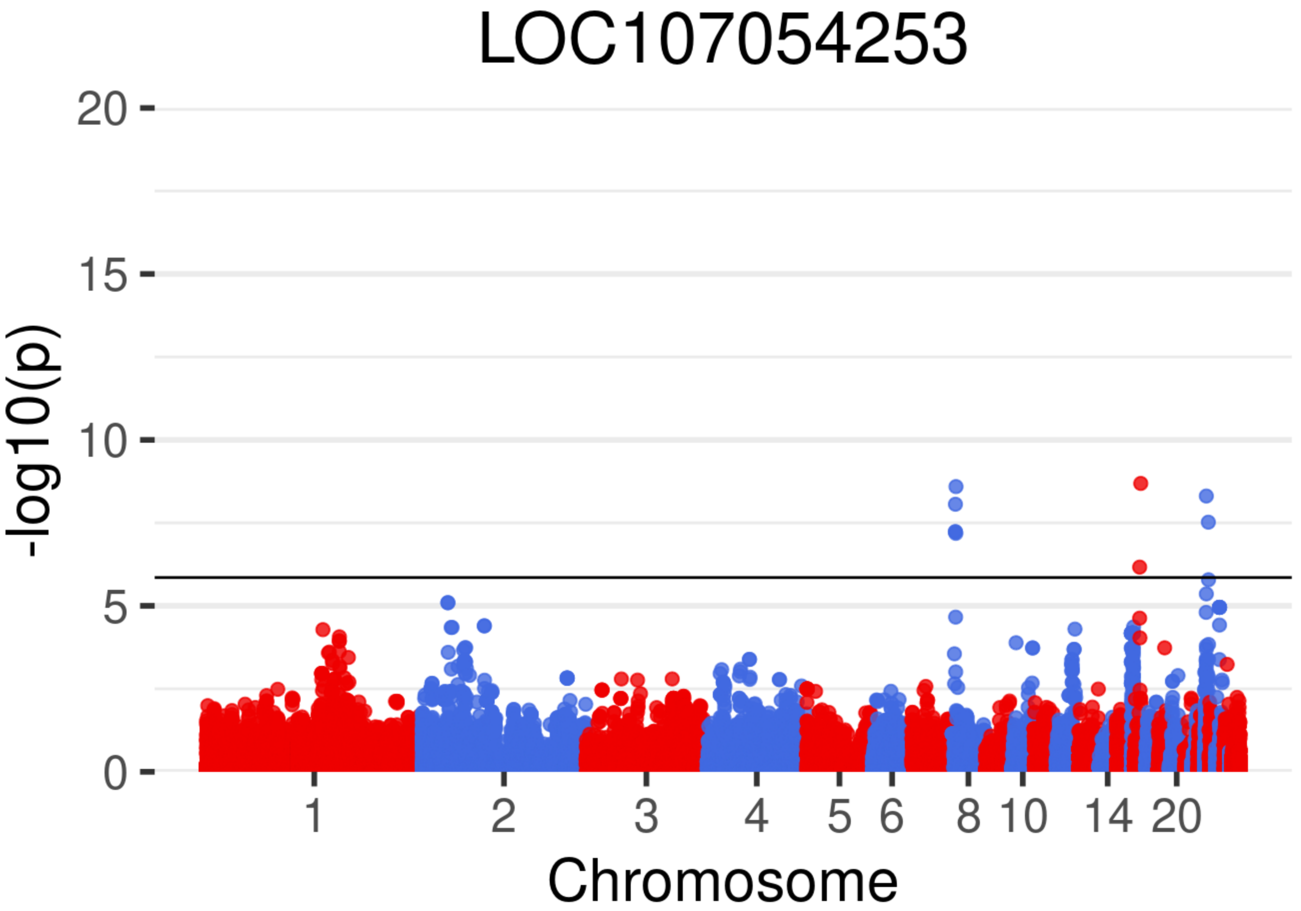
LOC107054253
-log10(p)
20
15
10
5
0
1
2
3
4
5
6
8
10
14
20
Chromosome

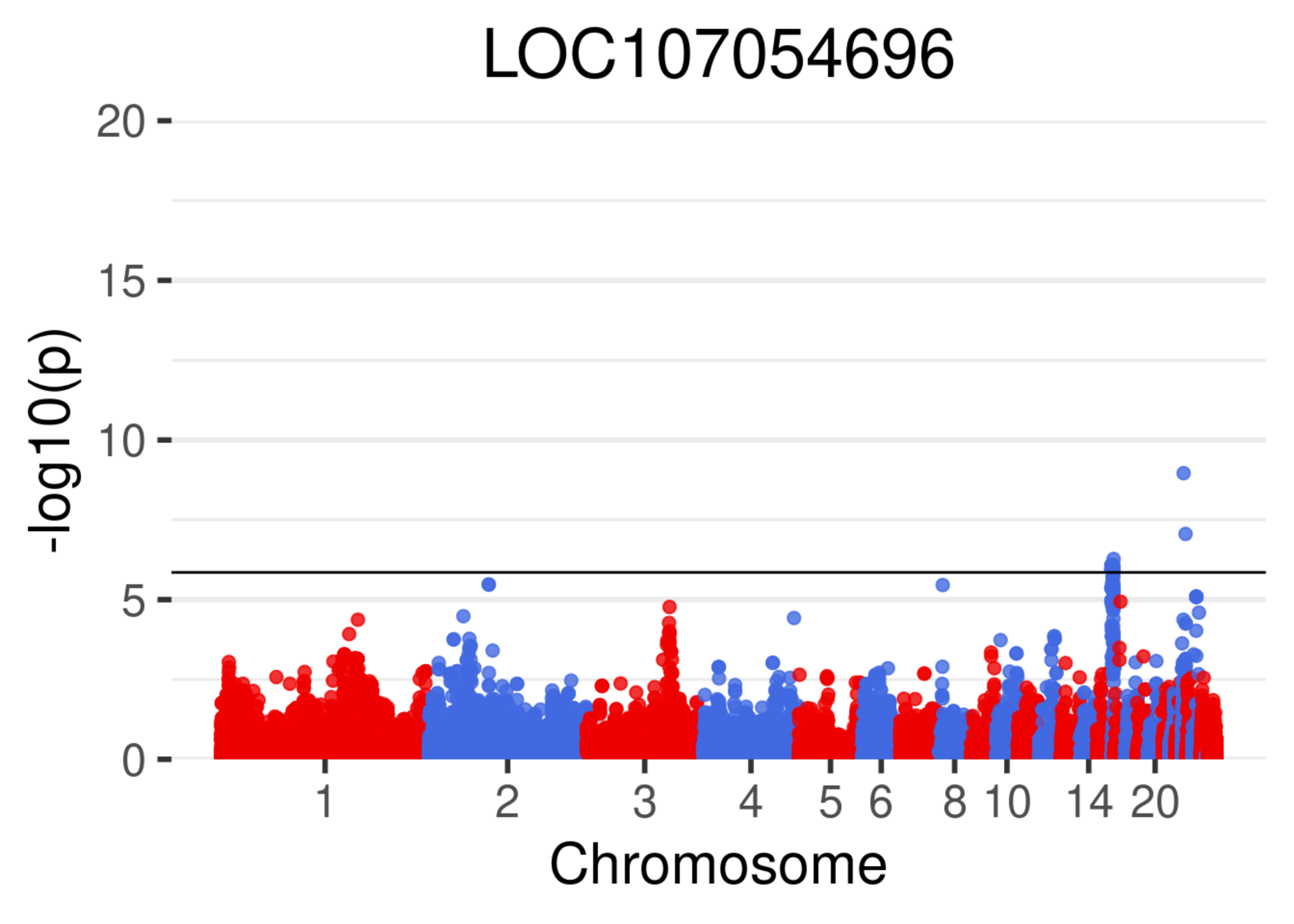
LOC107054696
-log10(p)
20
15
10
5
0
1
2
3
4
5
6
8
10
14
20
Chromosome

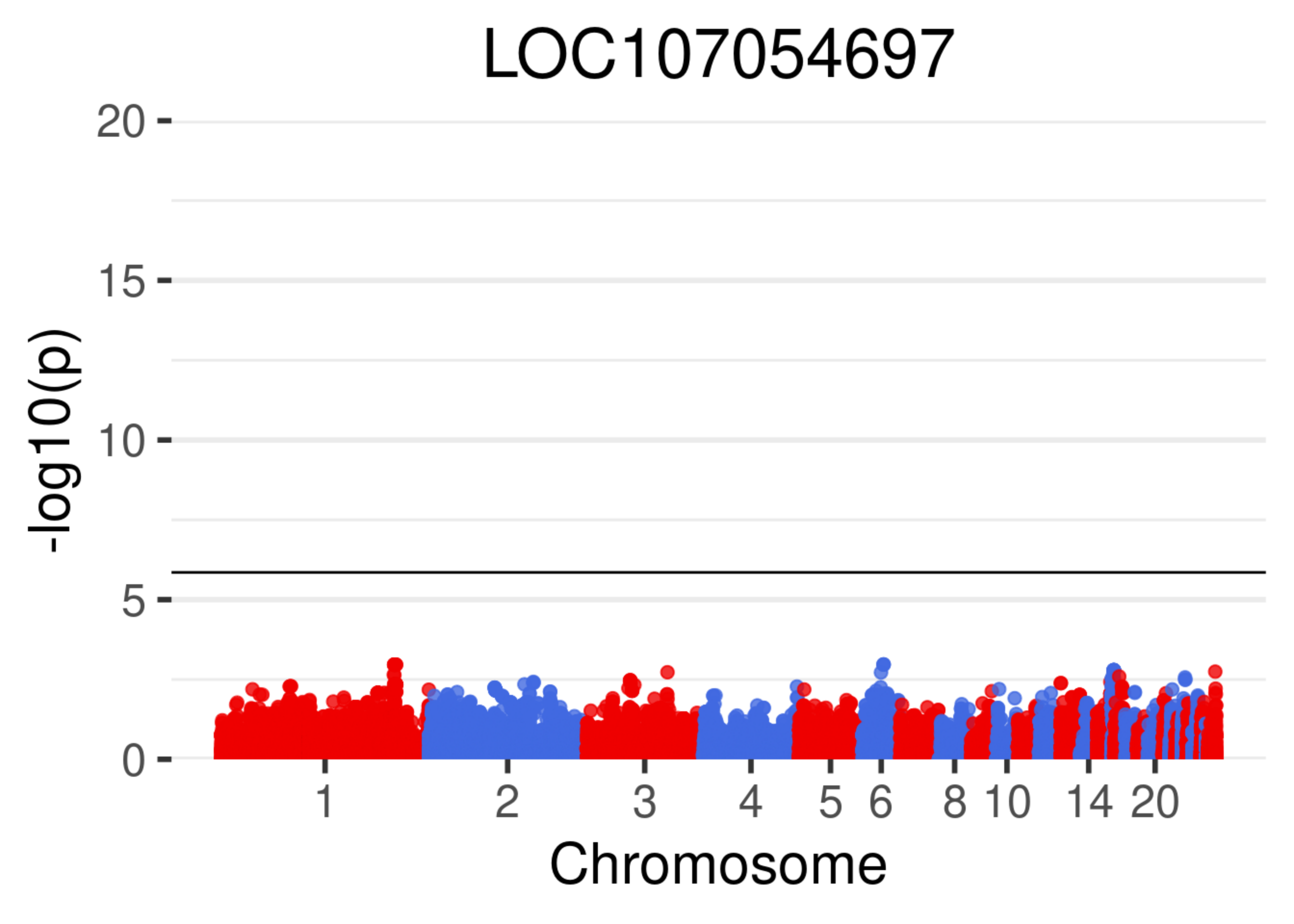
LOC107054697
20
15
10
5
0
-log10(p)
1
2
3
4
5
6
8
10
14
20
Chromosome

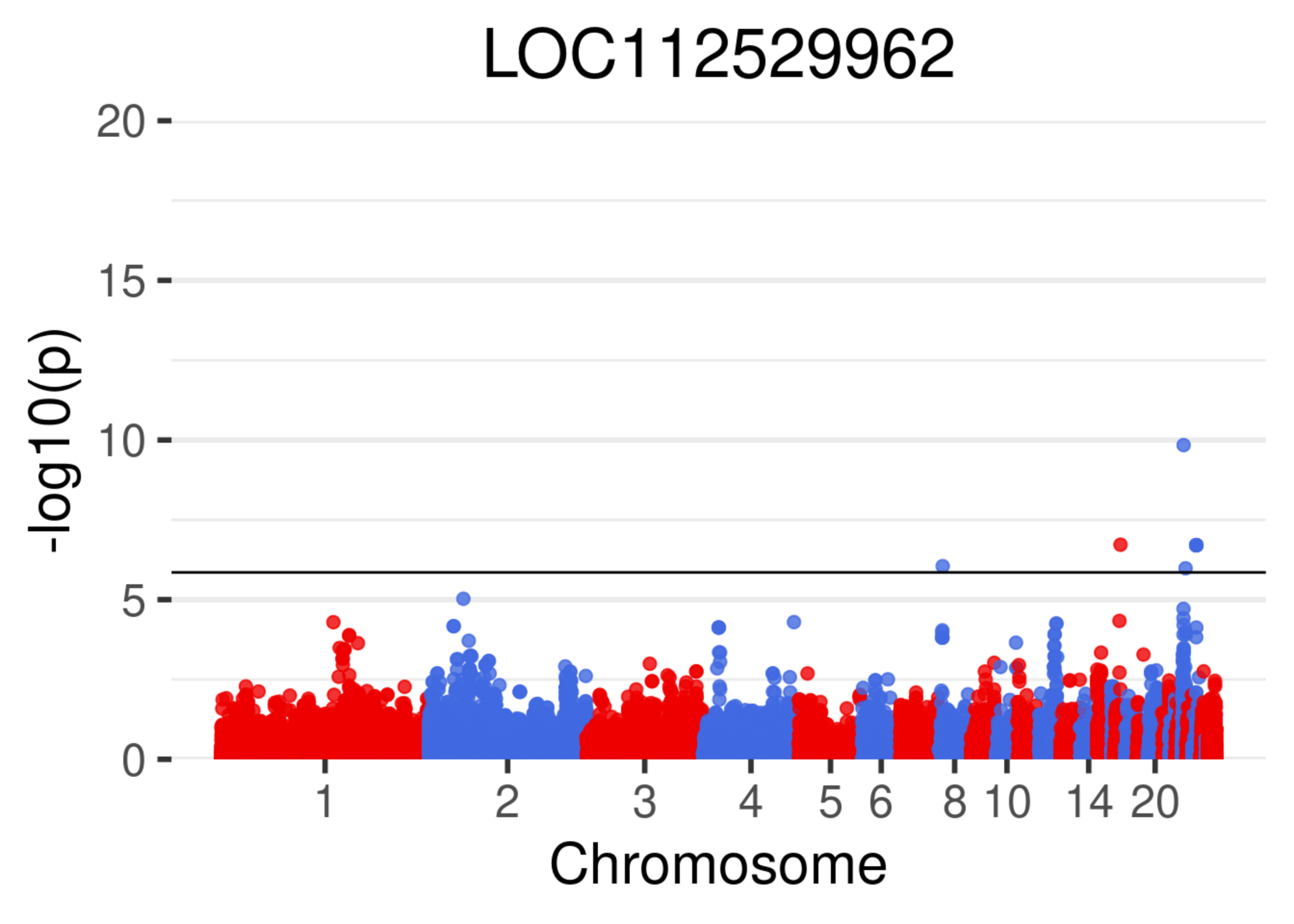
LOC112529962
-log10(p)
20
15
10
5
0
1
2
3
4
5
6
8
10
14
20
Chromosome

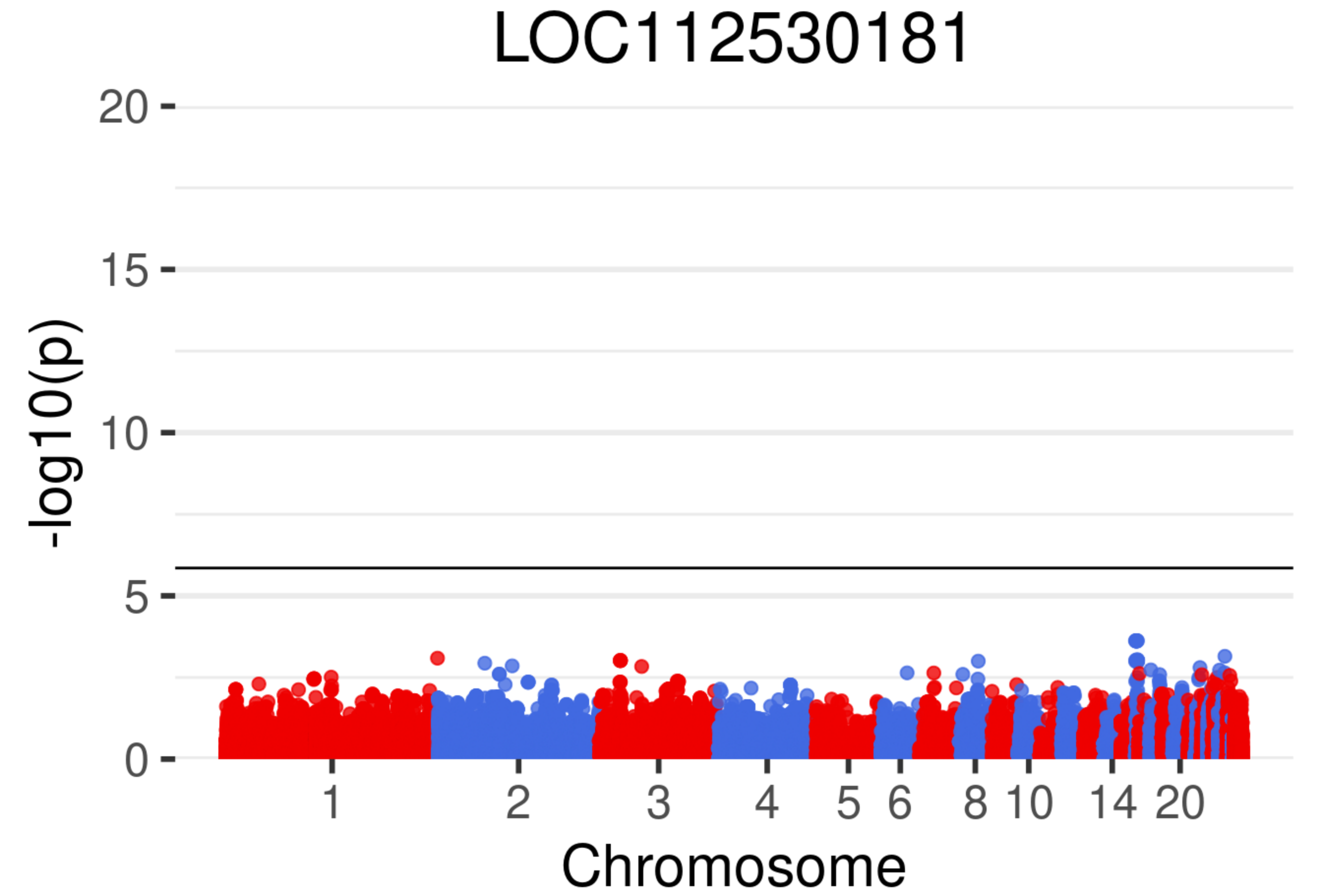
LOC112530181
20
15
10
5
0
-log10(p)
1
2
3
4
5
6
8
10
14
20
Chromosome

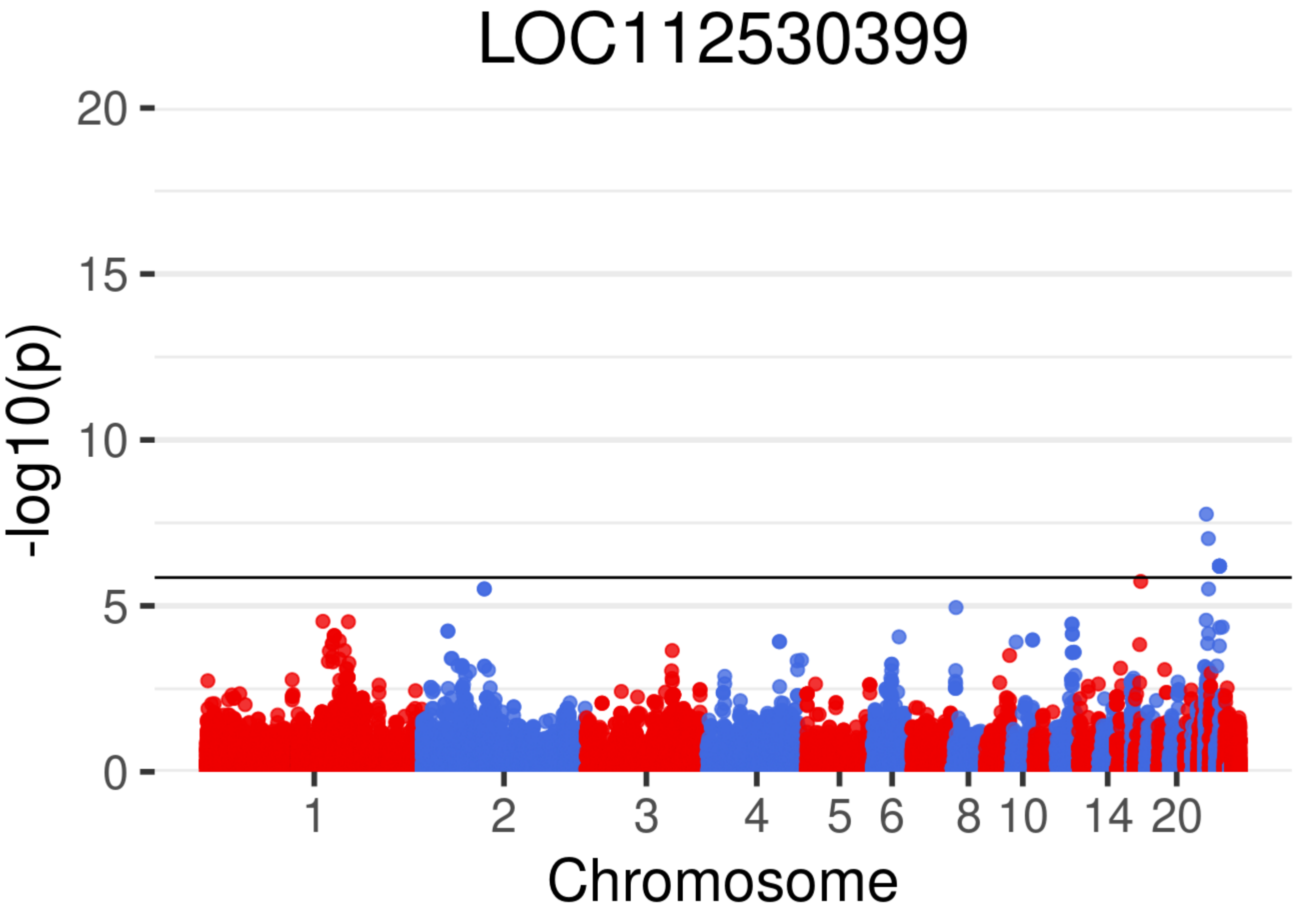
LOC112530399
-log10(p)
20
15
10
5
0
1
2
3
4
5
6
8
10
14
20
Chromosome

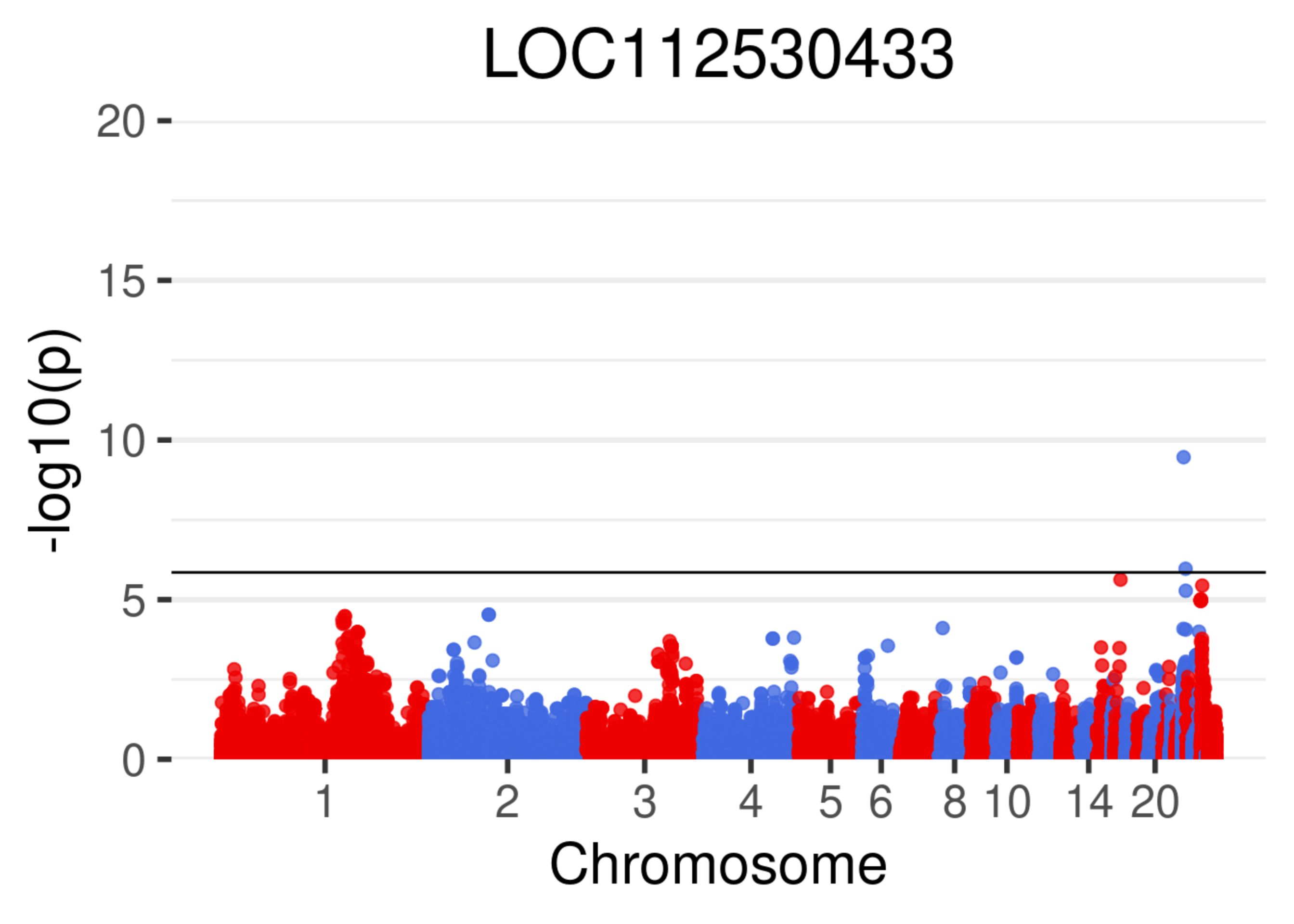
LOC112530433
20
15
10
5
0
-log10(p)
1
2
3
4
5
6
8
10
14
20
Chromosome

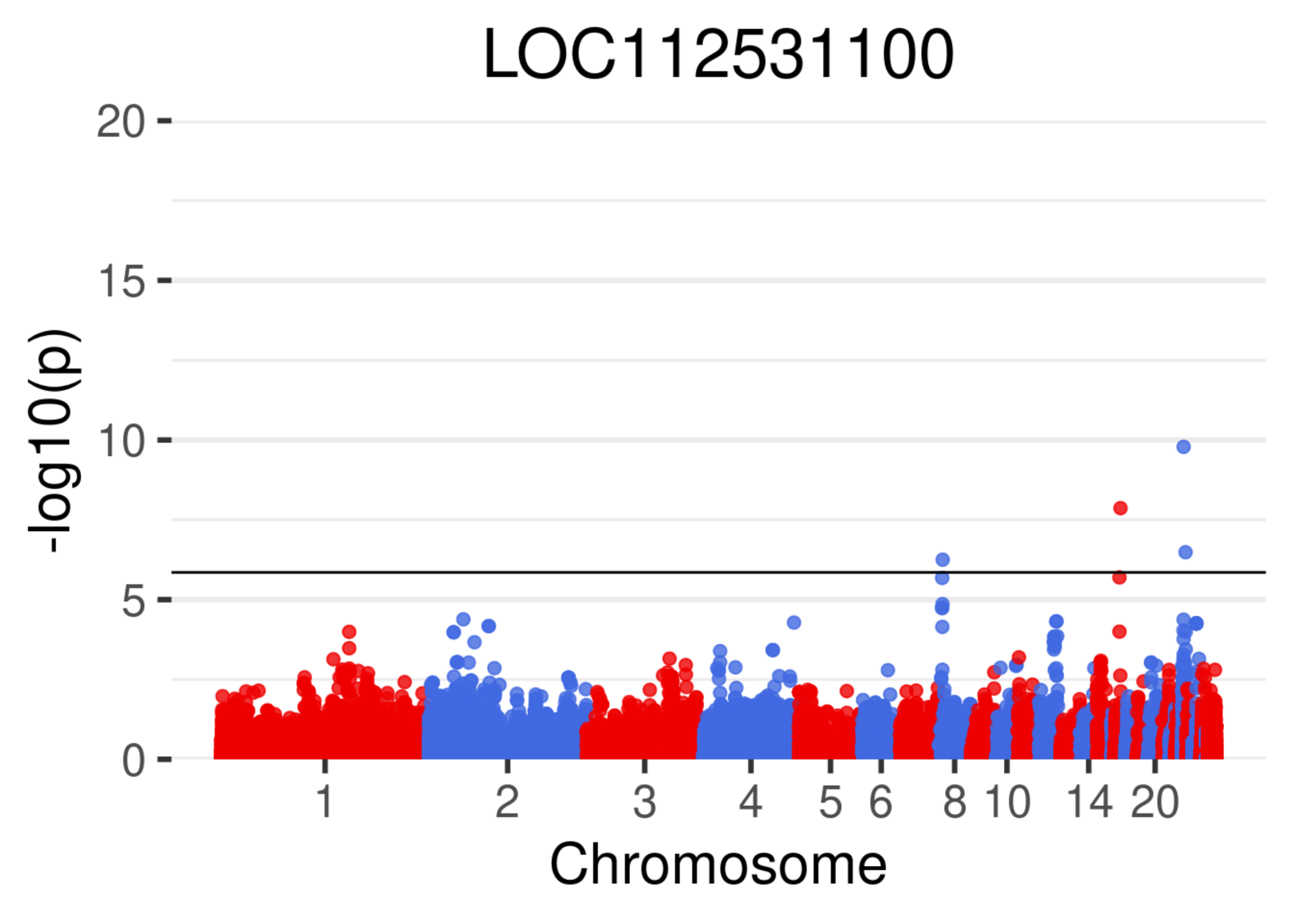

LOC112531100
-log10(p)
20
15
10
5
0
1
2
3
4
5
6
8
10
14
20
Chromosome

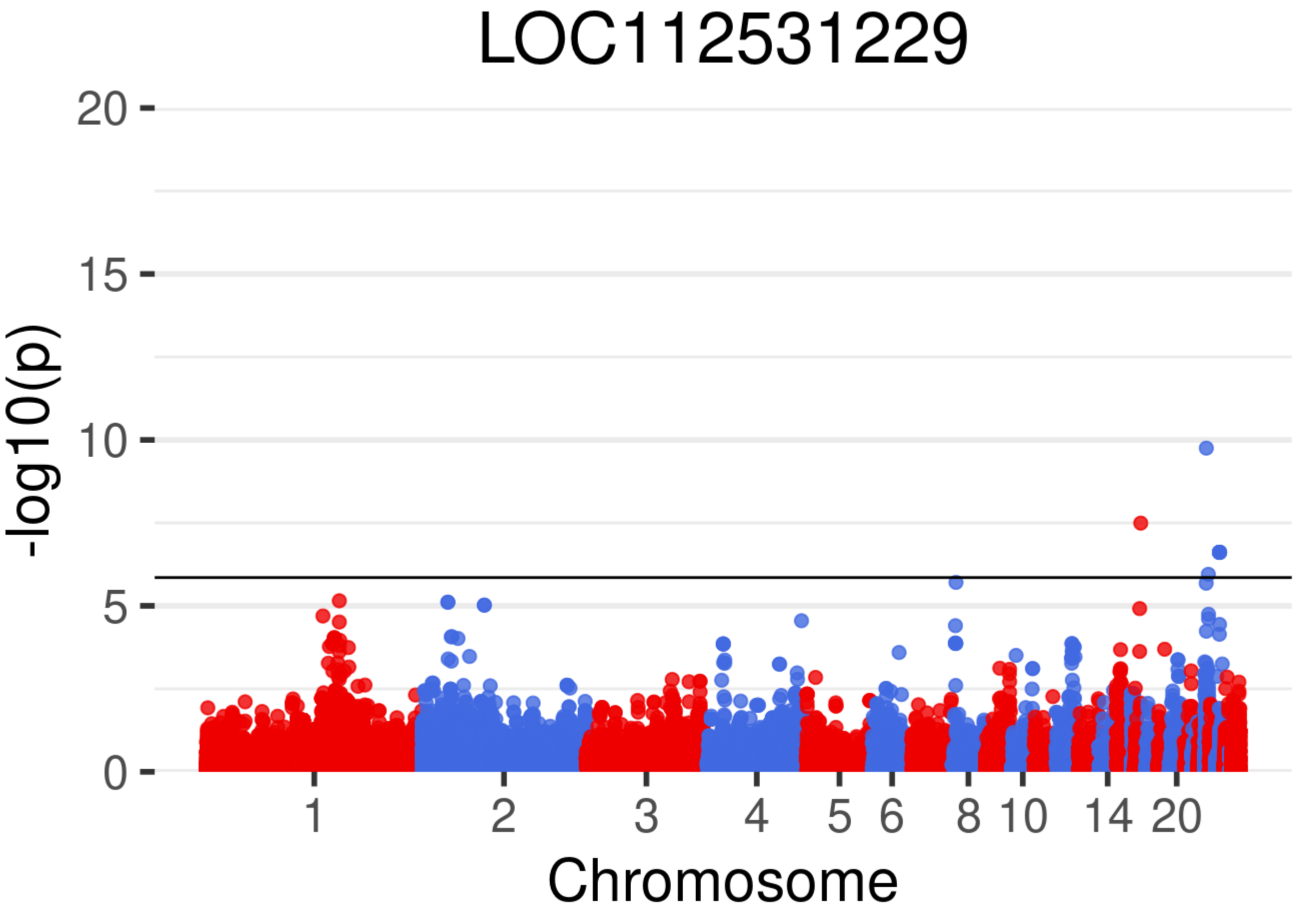

LOC112531229
-log10(p)
20
15
10
5
0
1
2
3
4
5
6
8
10
14
20
Chromosome

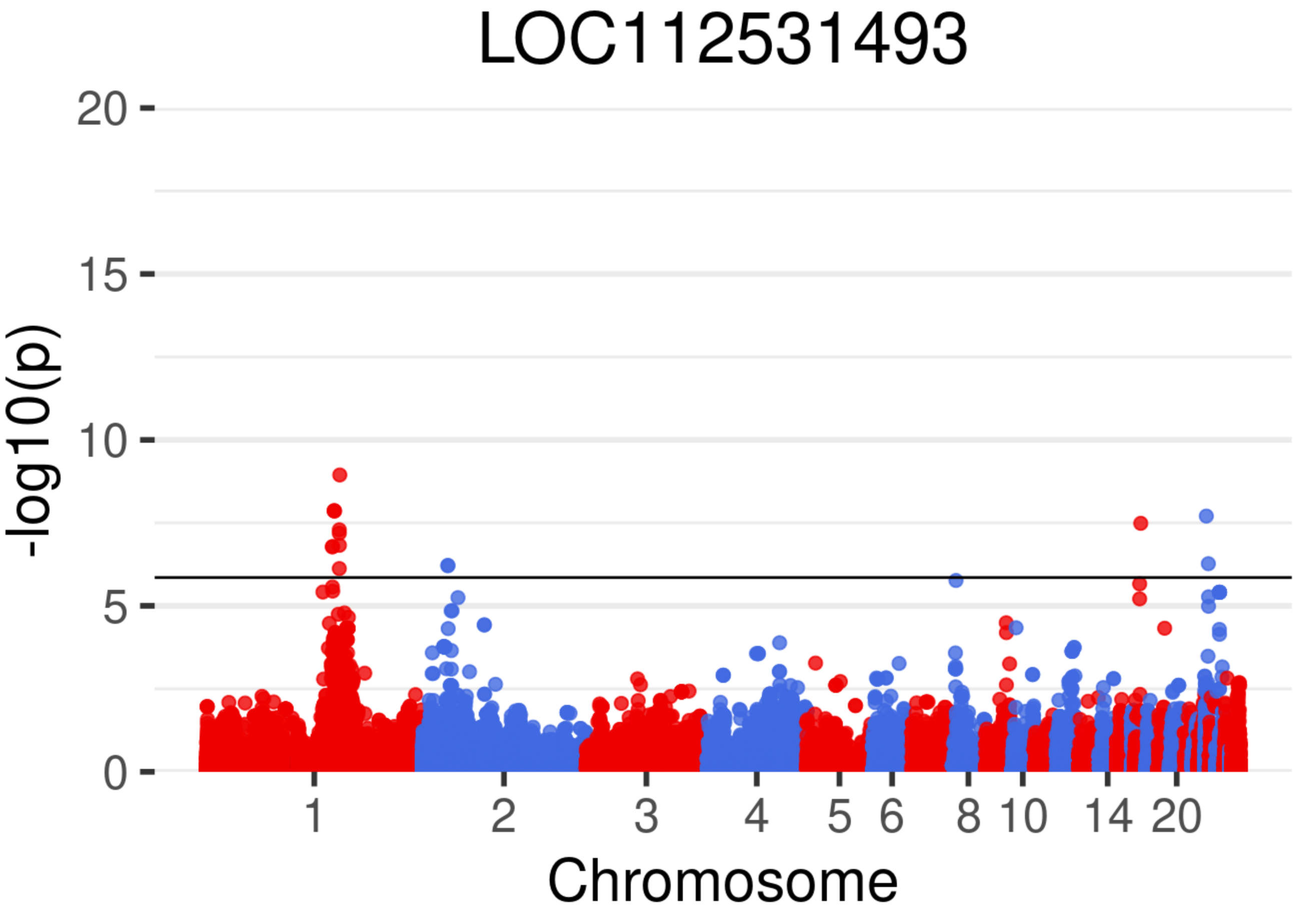
LOC112531493
-log10(p)
20
15
10
5
0
1
2
3
4
5
6
8
10
14
20
Chromosome

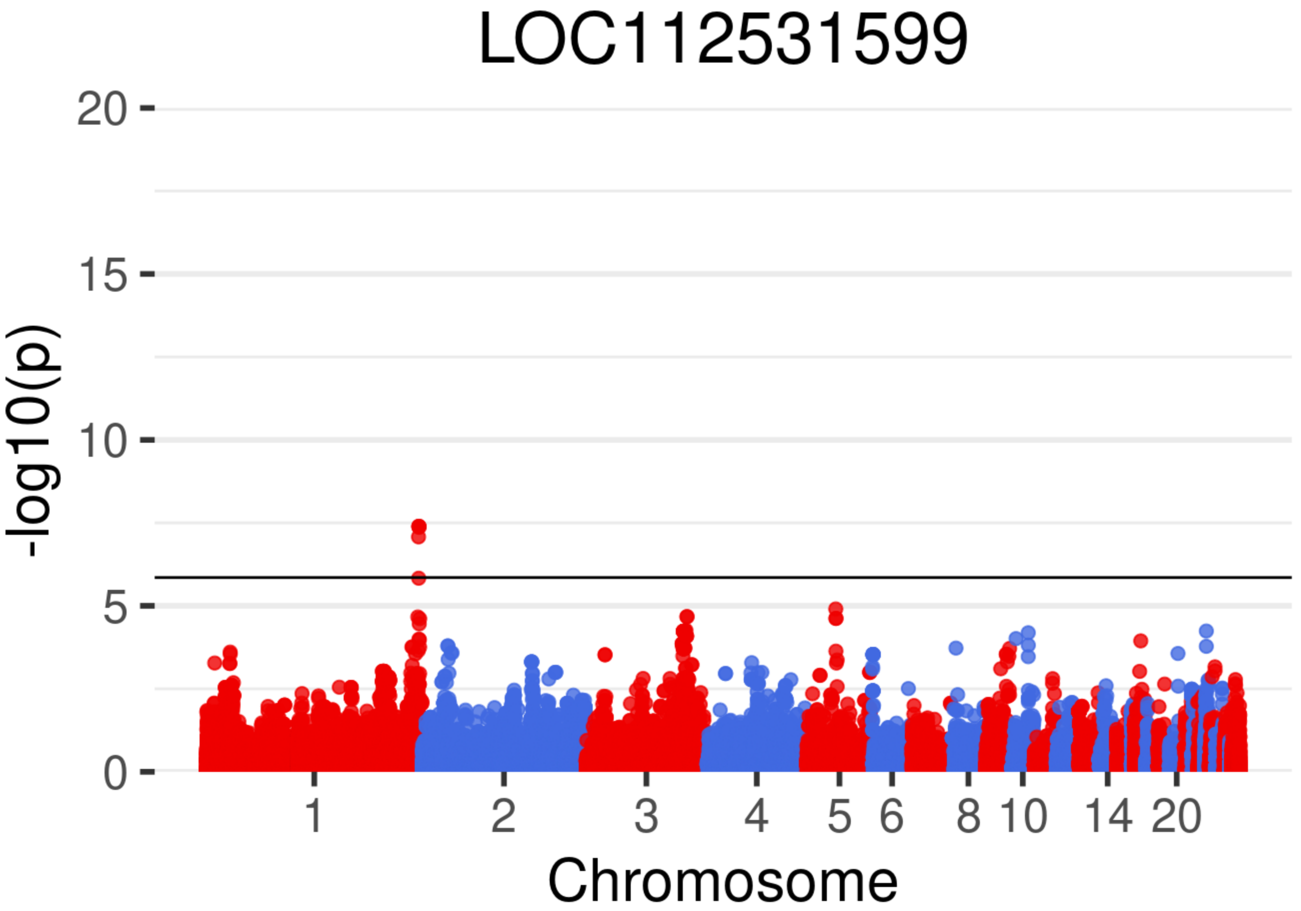

LOC112531599
-log10(p)
20
15
10
5
0
1
2
3
4
5
6
8
10
14
20
Chromosome

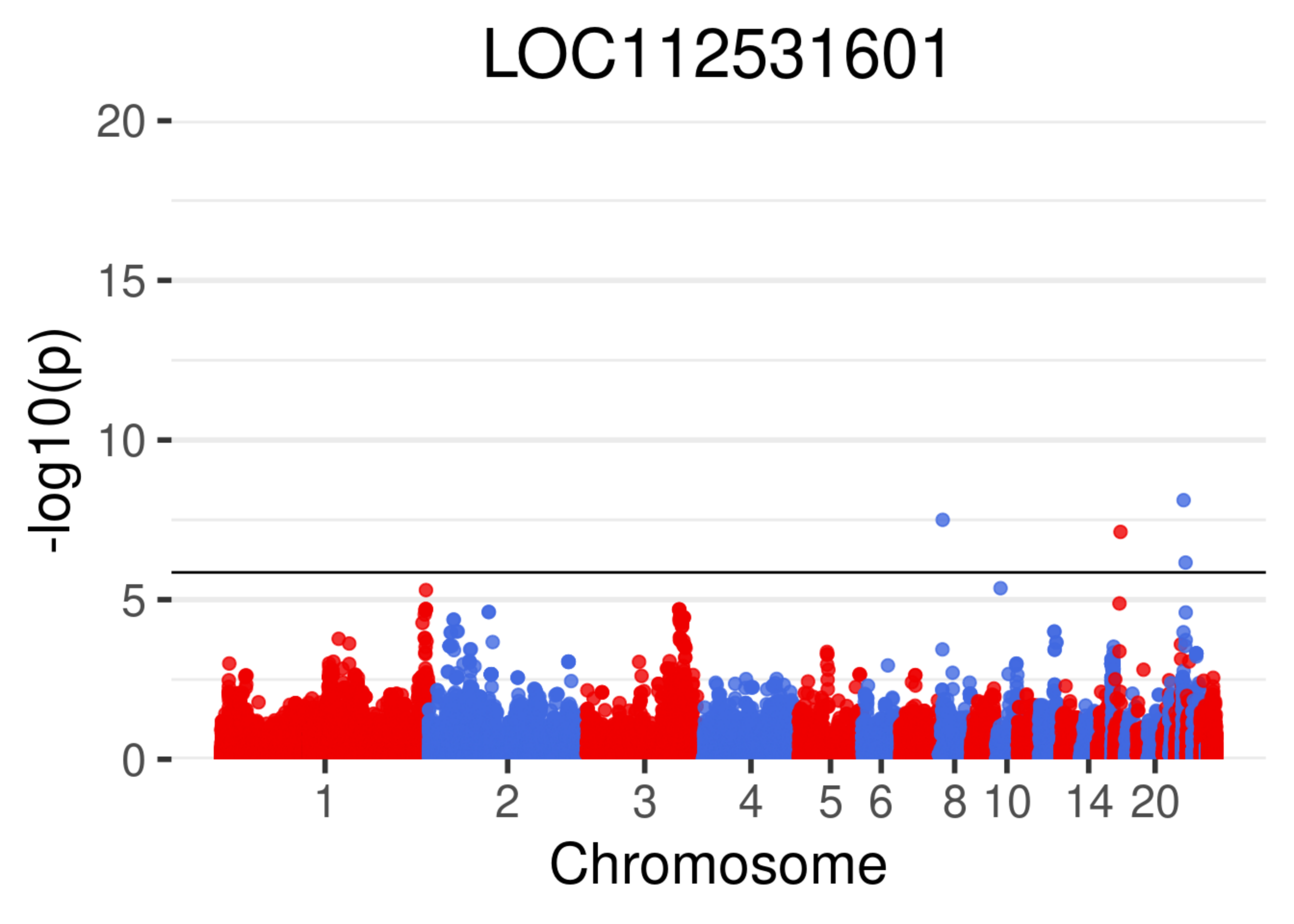

LOC112531601
20
15
10
5
0
-log10(p)
1
2
3
4
5
6
8
10
14
20
Chromosome

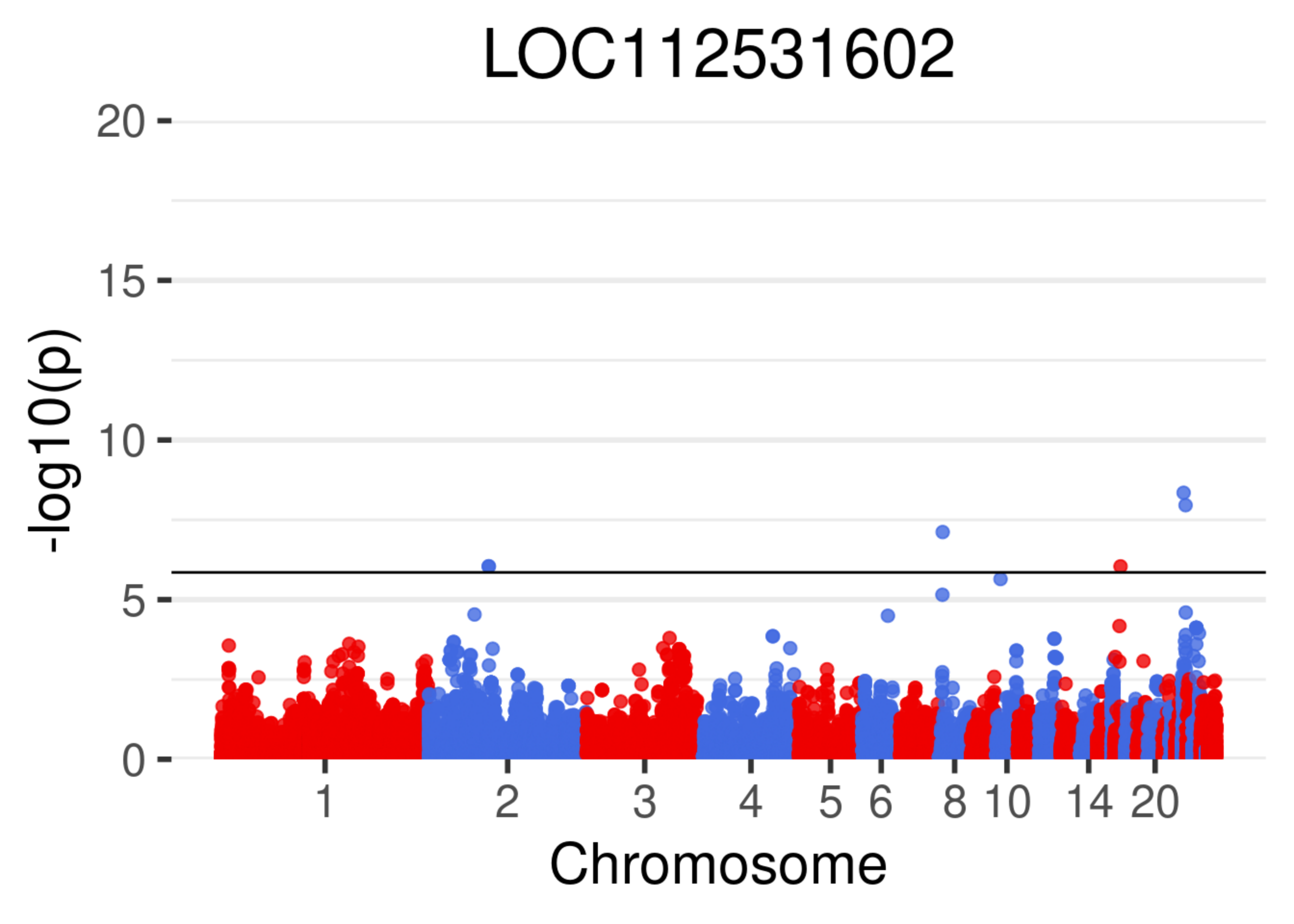
LOC112531602
-log10(p)
20
15
10
5
0
1
2
3
4
5
6
8
10
14
20
Chromosome

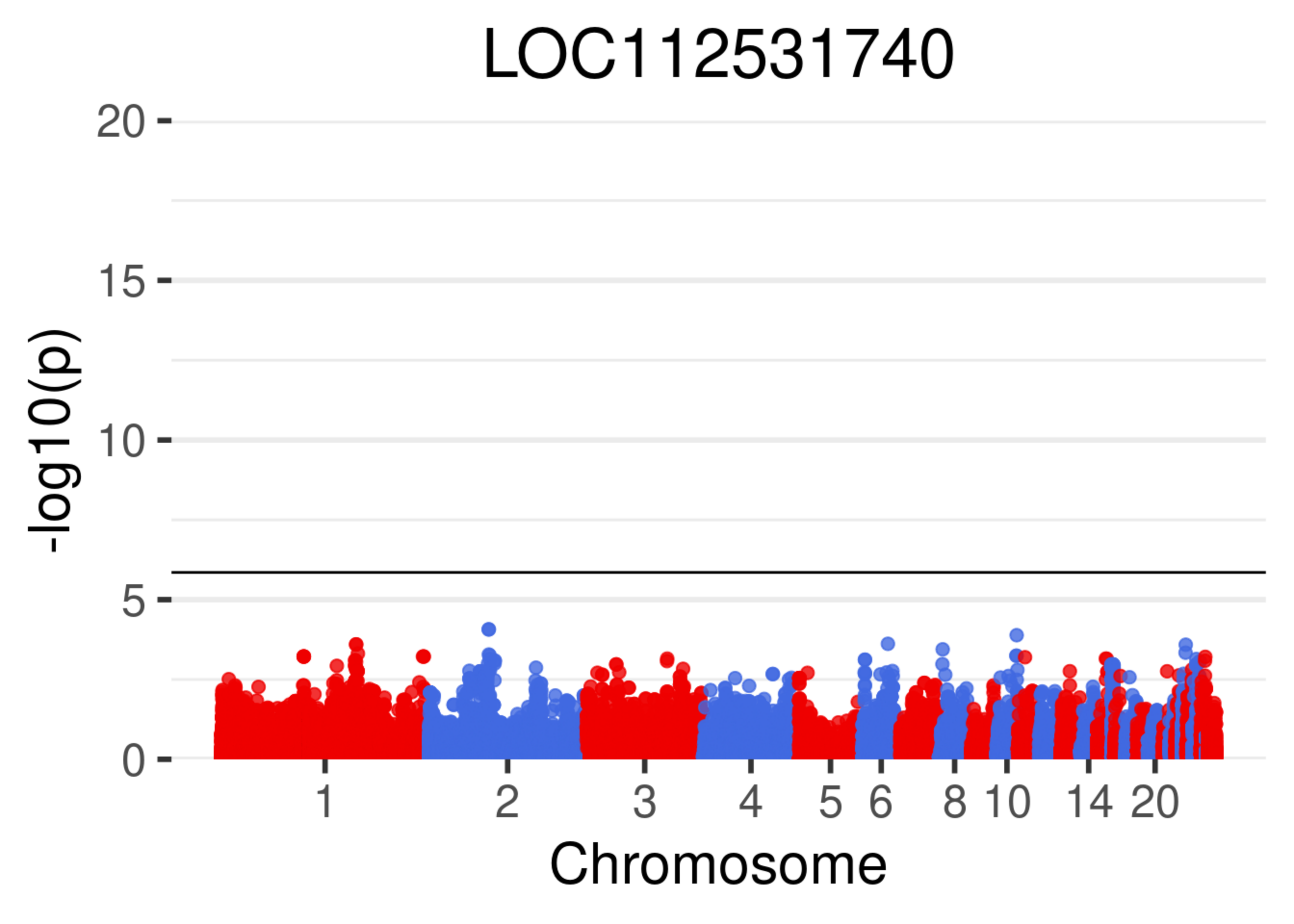

LOC112531740
-log10(p)
20
15
10
5
0
1
2
3
4
5
6
8
10
14
20
Chromosome

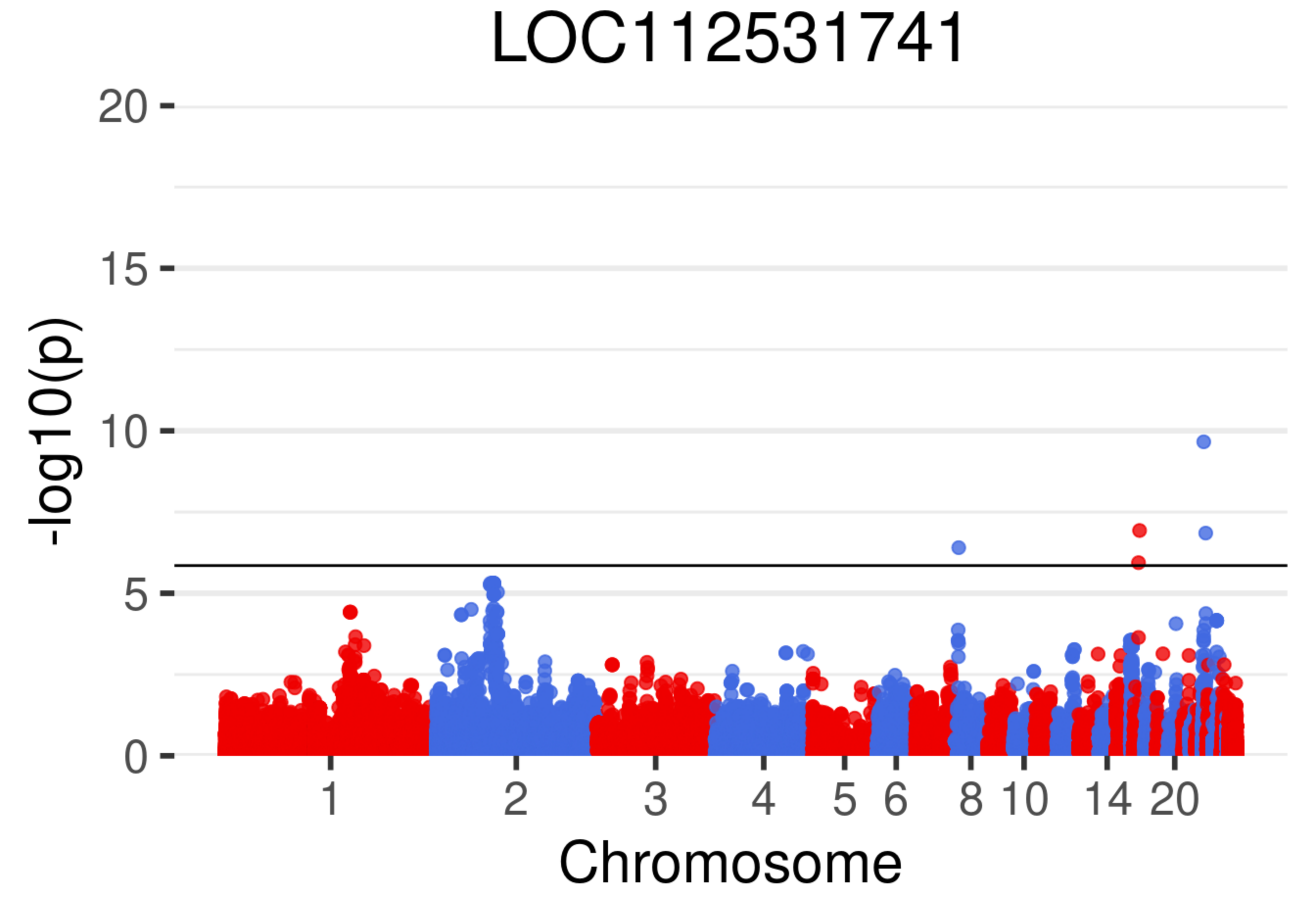
LOC112531741
-log10(p)
20
15
10
5
0
1
2
3
4
5
6
8
10
14
20
Chromosome

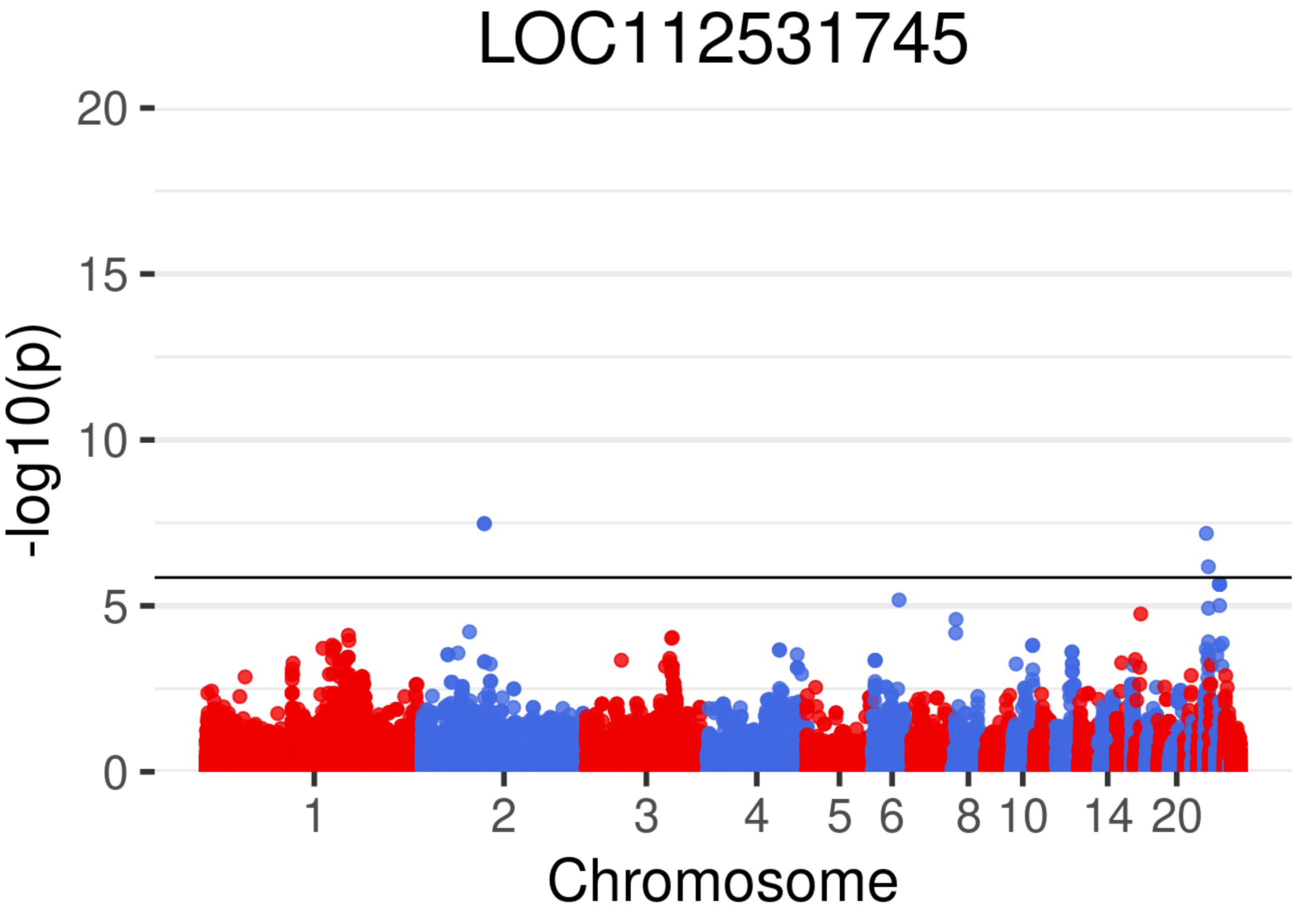
LOC112531745
-log10(p)
20
15
10
5
0
1
2
3
4
5
6
8
10
14
20
Chromosome

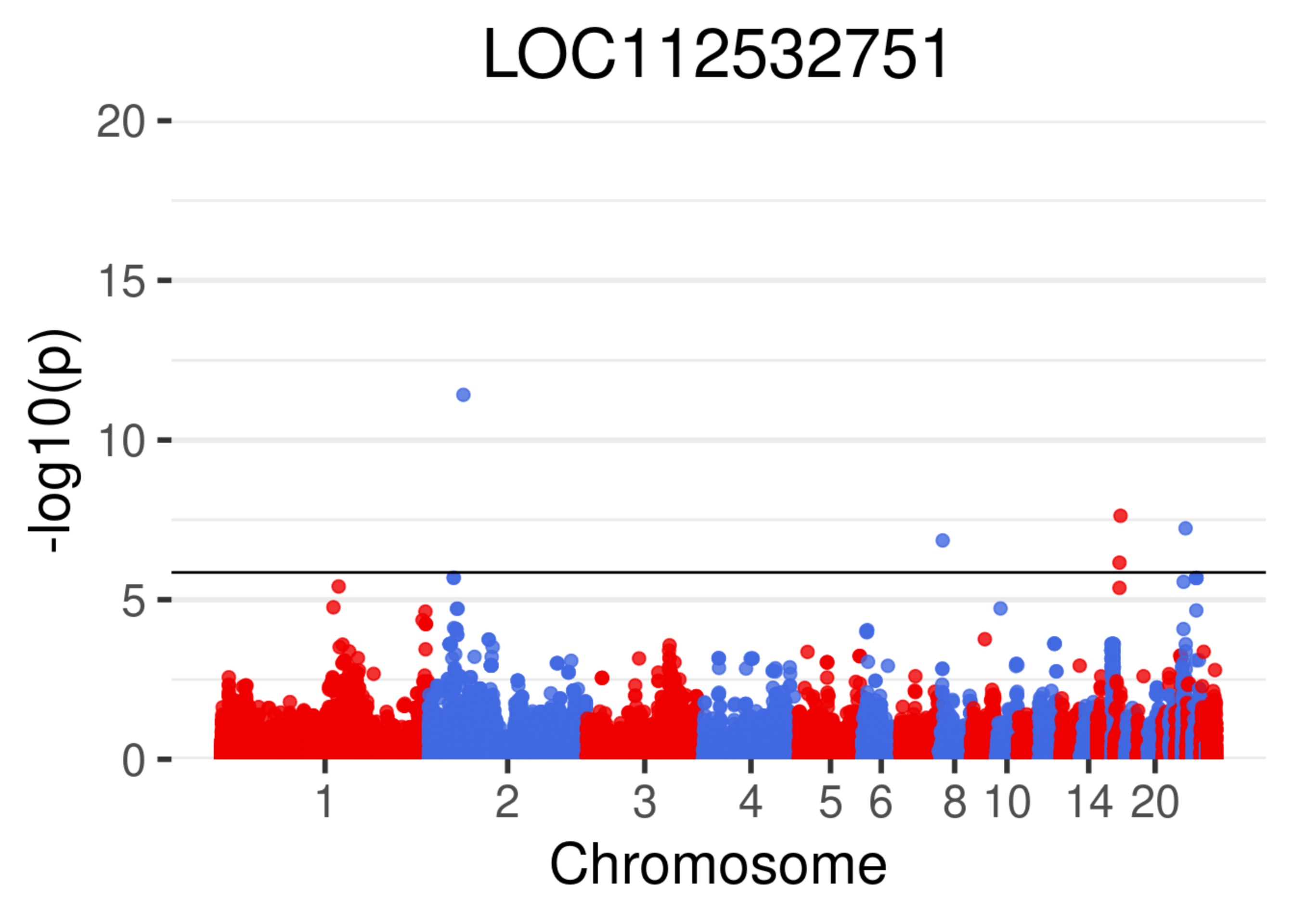
LOC112532751
-log10(p)
20
15
10
5
0
1
2
3
4
5
6
8
10
14
20
Chromosome

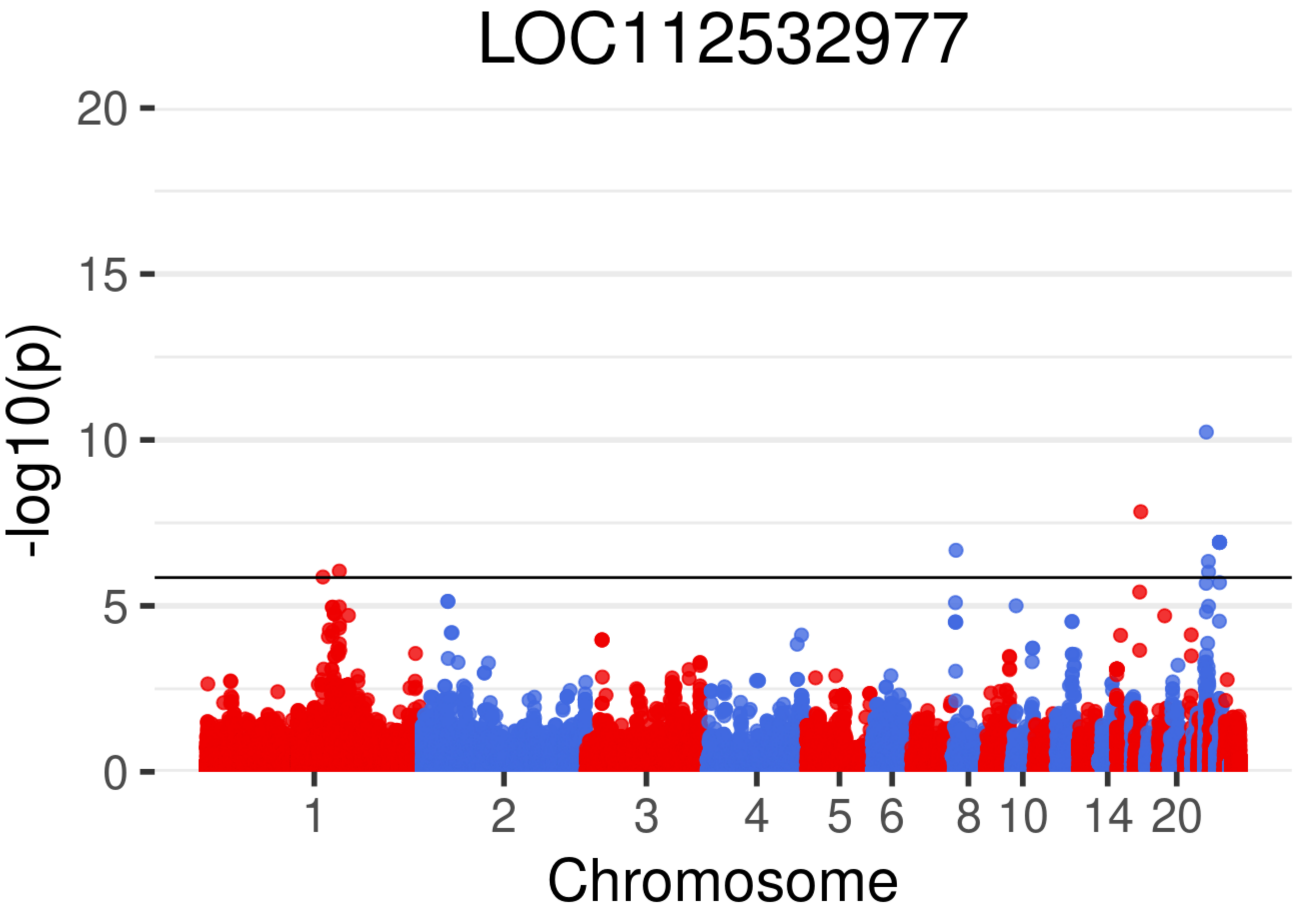

LOC112532977
-log10(p)
20
15
10
5
0
1
2
3
4
5
6
8
10
14
20
Chromosome

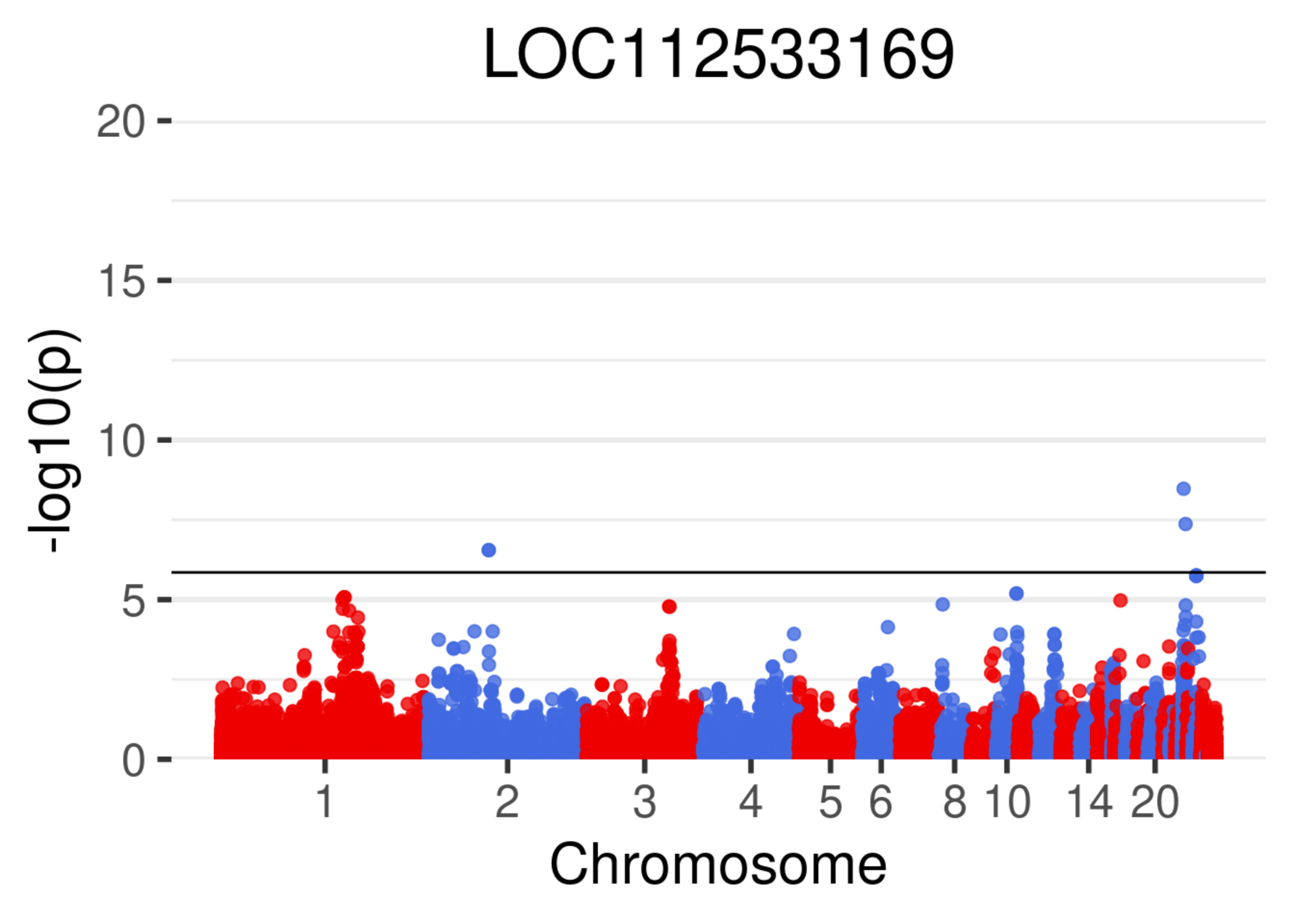

LOC112533169
-log10(p)
20
15
10
5
0
1
2
3
4
5
6
8
10
14
20
Chromosome

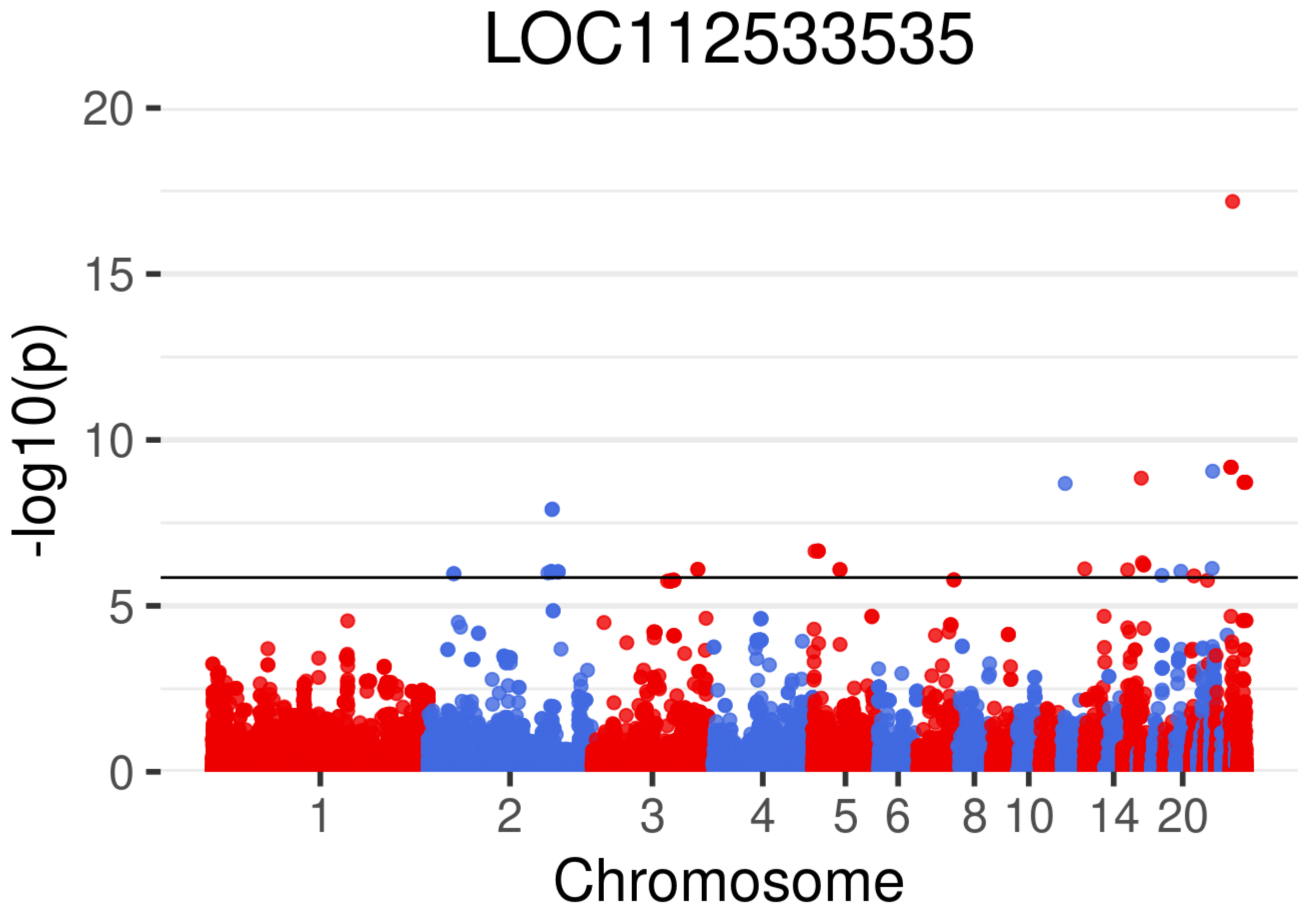
LOC112533535
-log10(p)
0
5
10
15
20
1
2
3
4
5
6
8
10
14
20
Chromosome

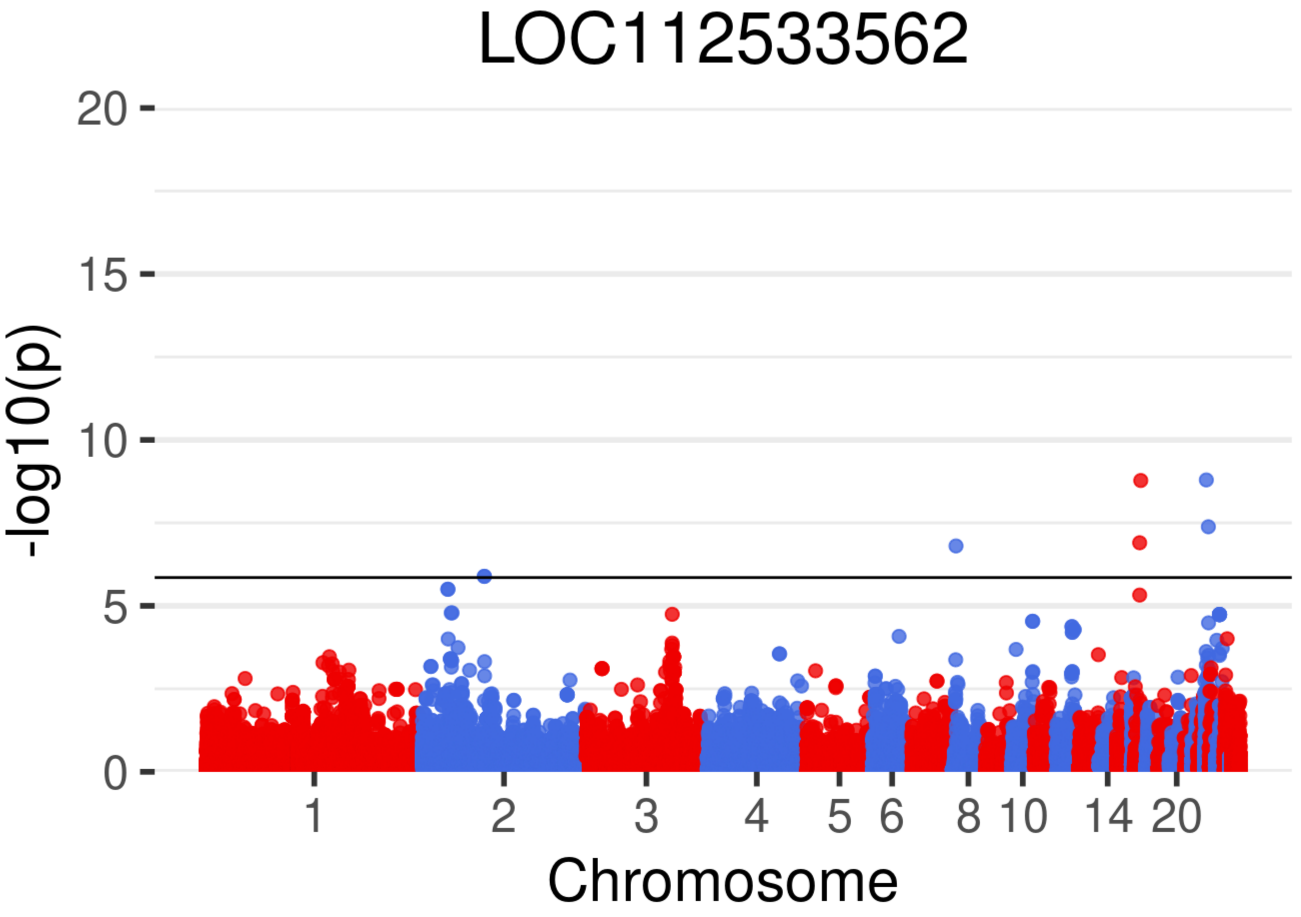
LOC112533562
-log10(p)
20
15
10
5
0
1
2
3
4
5
6
8
10
14
20
Chromosome

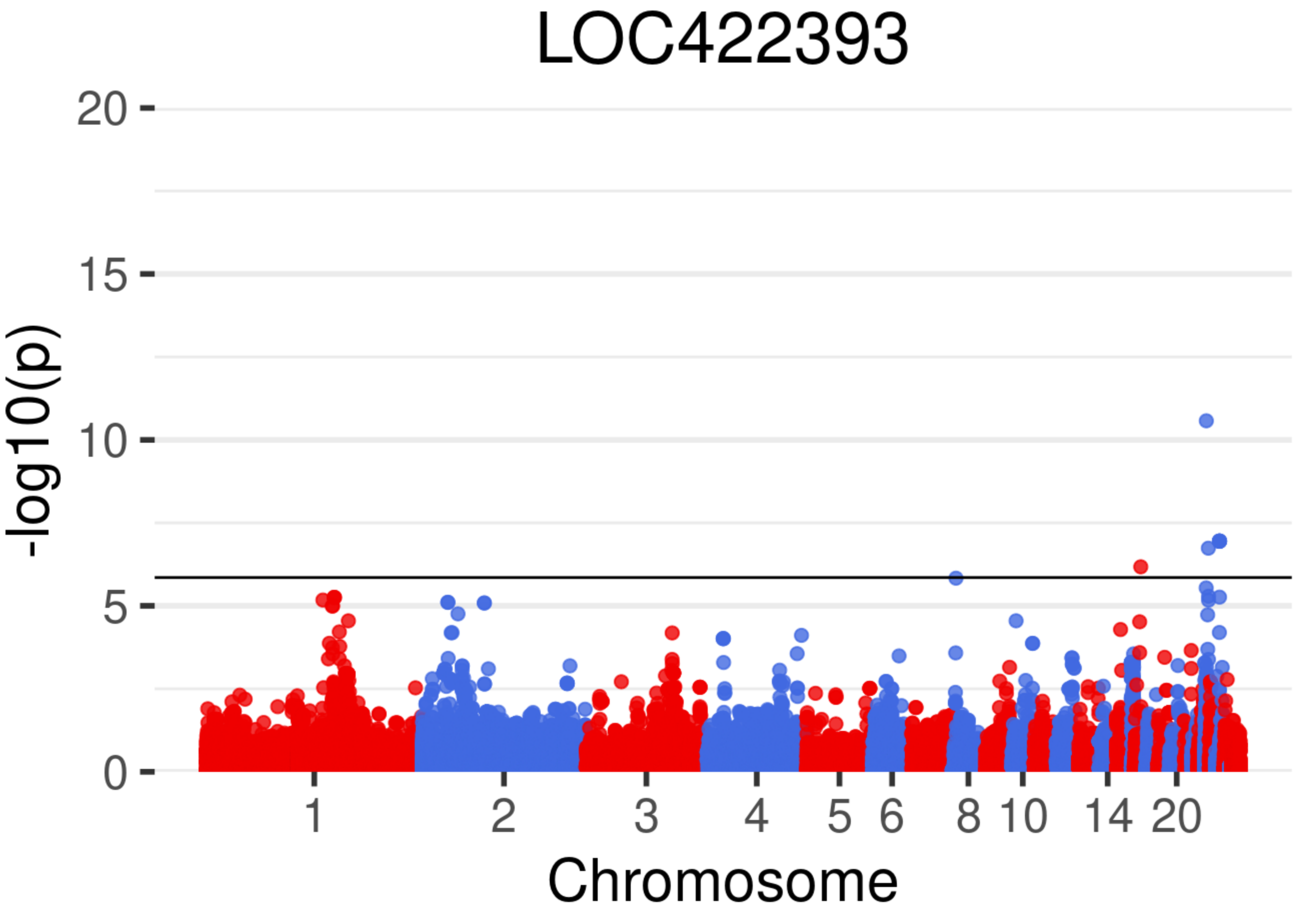
LOC422393
-log10(p)
0
5
10
15
20
1
2
3
4
5
6
8
10
14
20
Chromosome

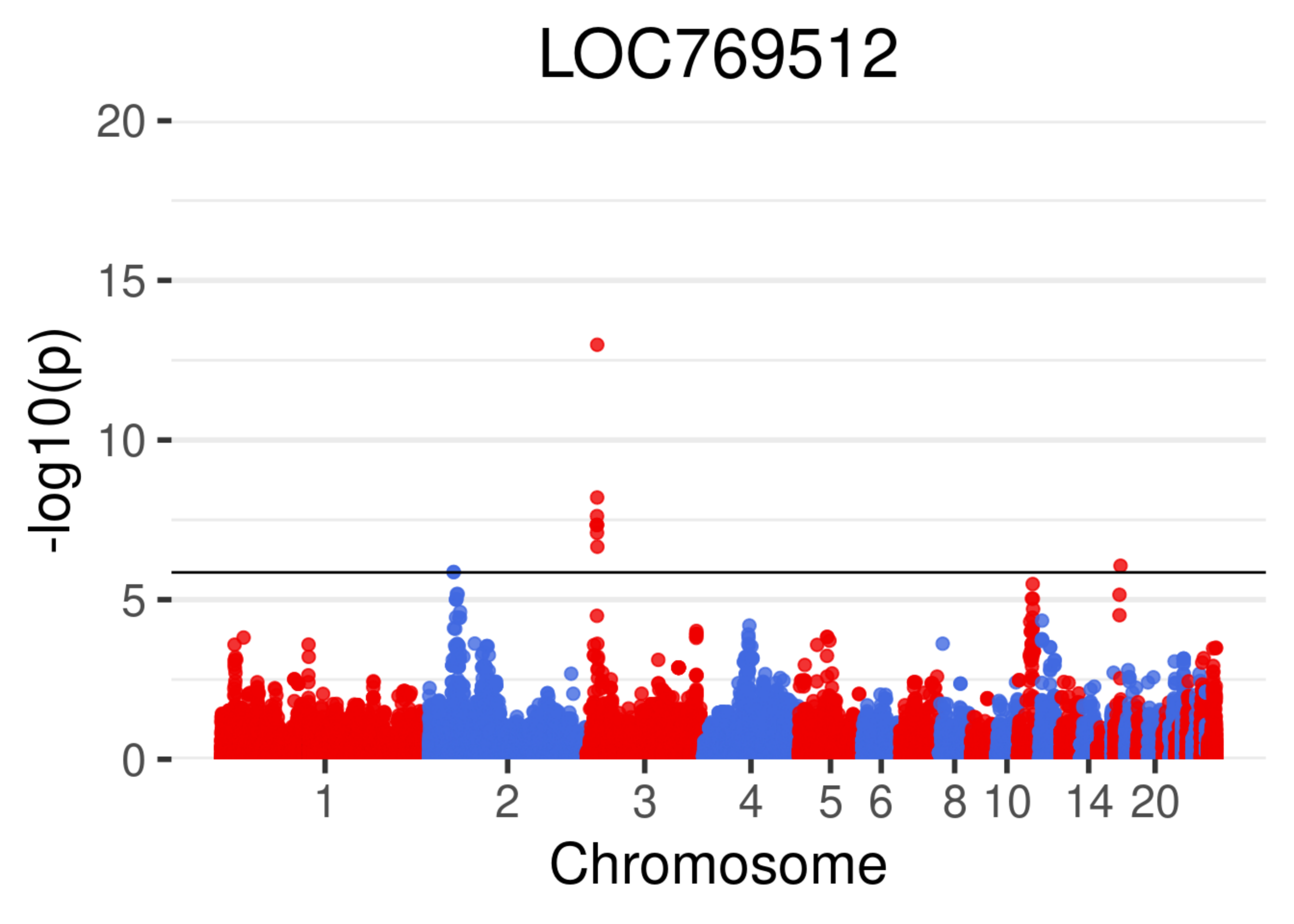
LOC769512
-log10(p)
20
15
10
5
0
1
2
3
4
5
6
8
10
14
20
Chromosome

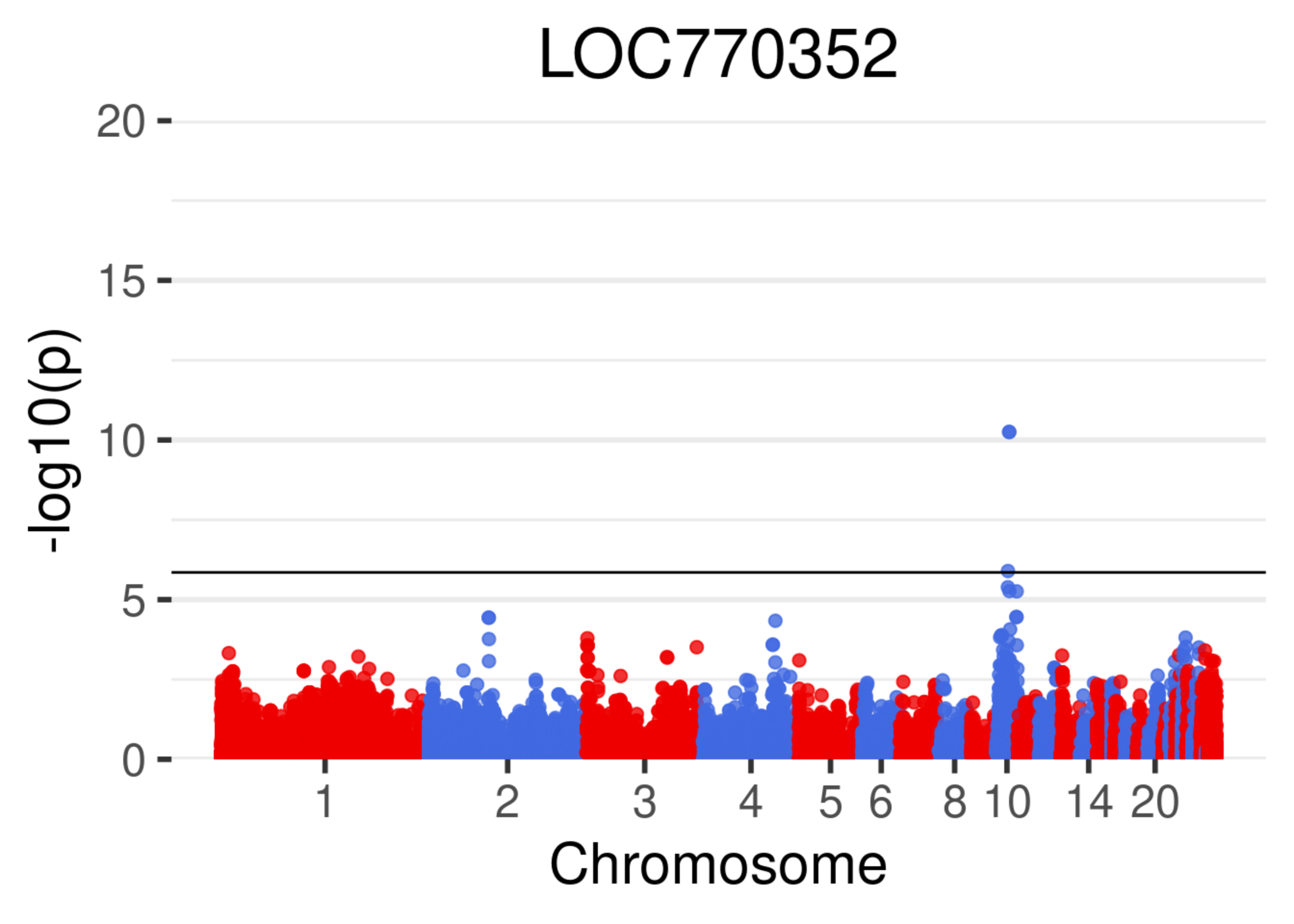

LOC770352
-log10(p)
20
15
10
5
0
1
2
3
4
5
6
8
10
14
20
Chromosome

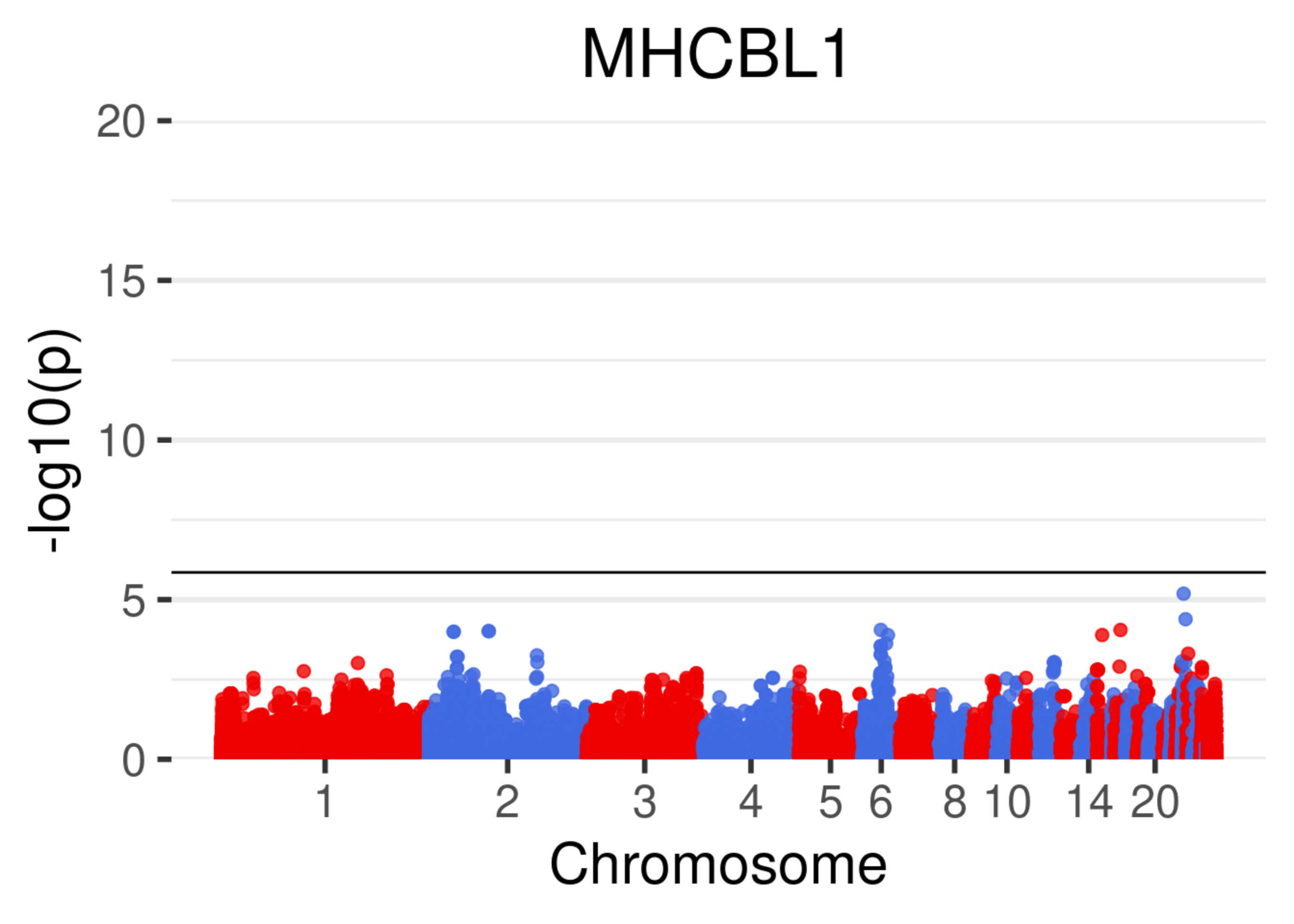
MHCBL1
-log10(p)
20
15
10
5
0
1
2
3
4
5
6
8
10
14
20
Chromosome

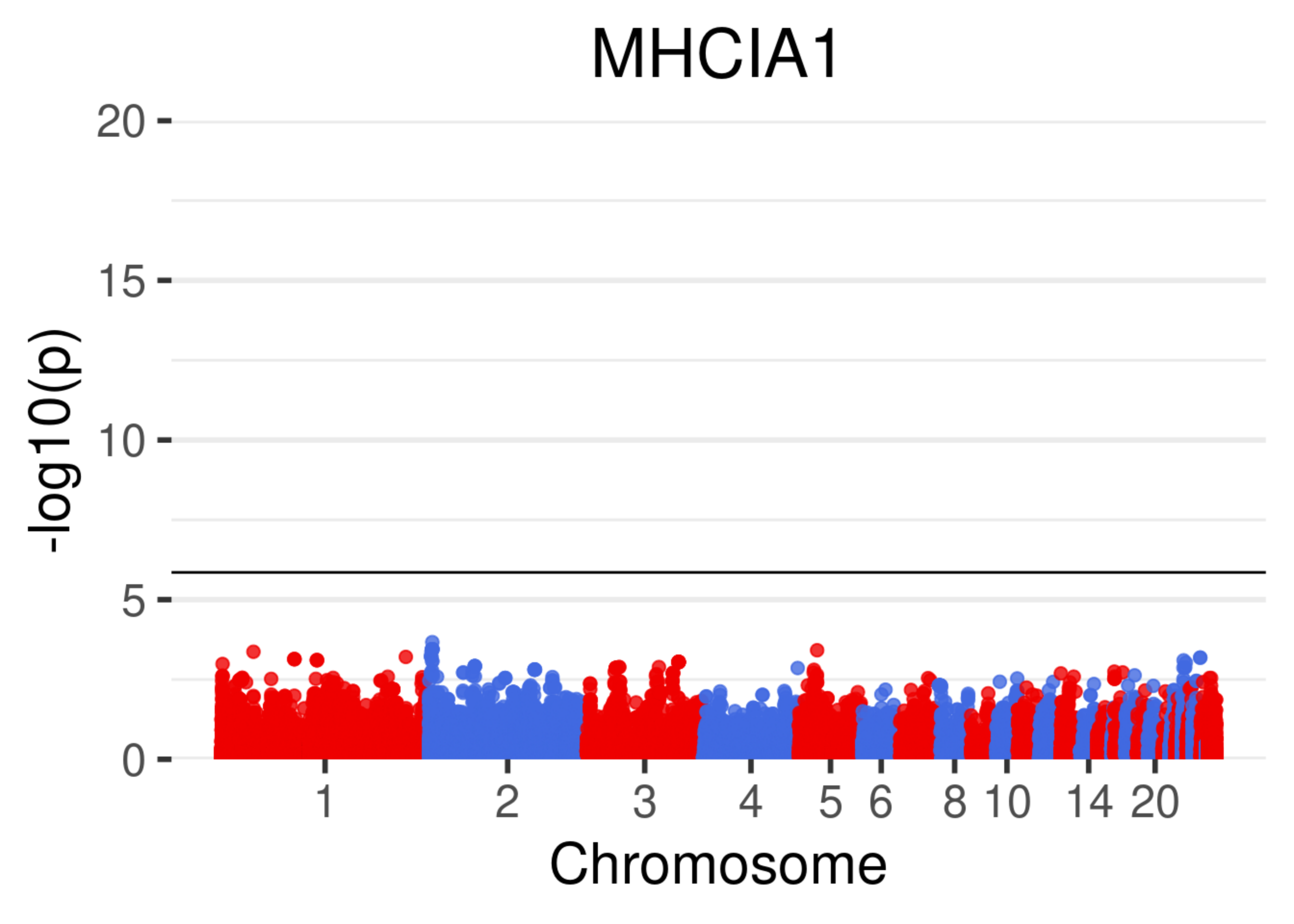
MHCIA1
20
15
10
5
0
-log10(p)
1
2
3
4
5
6
8
10
14
20
Chromosome

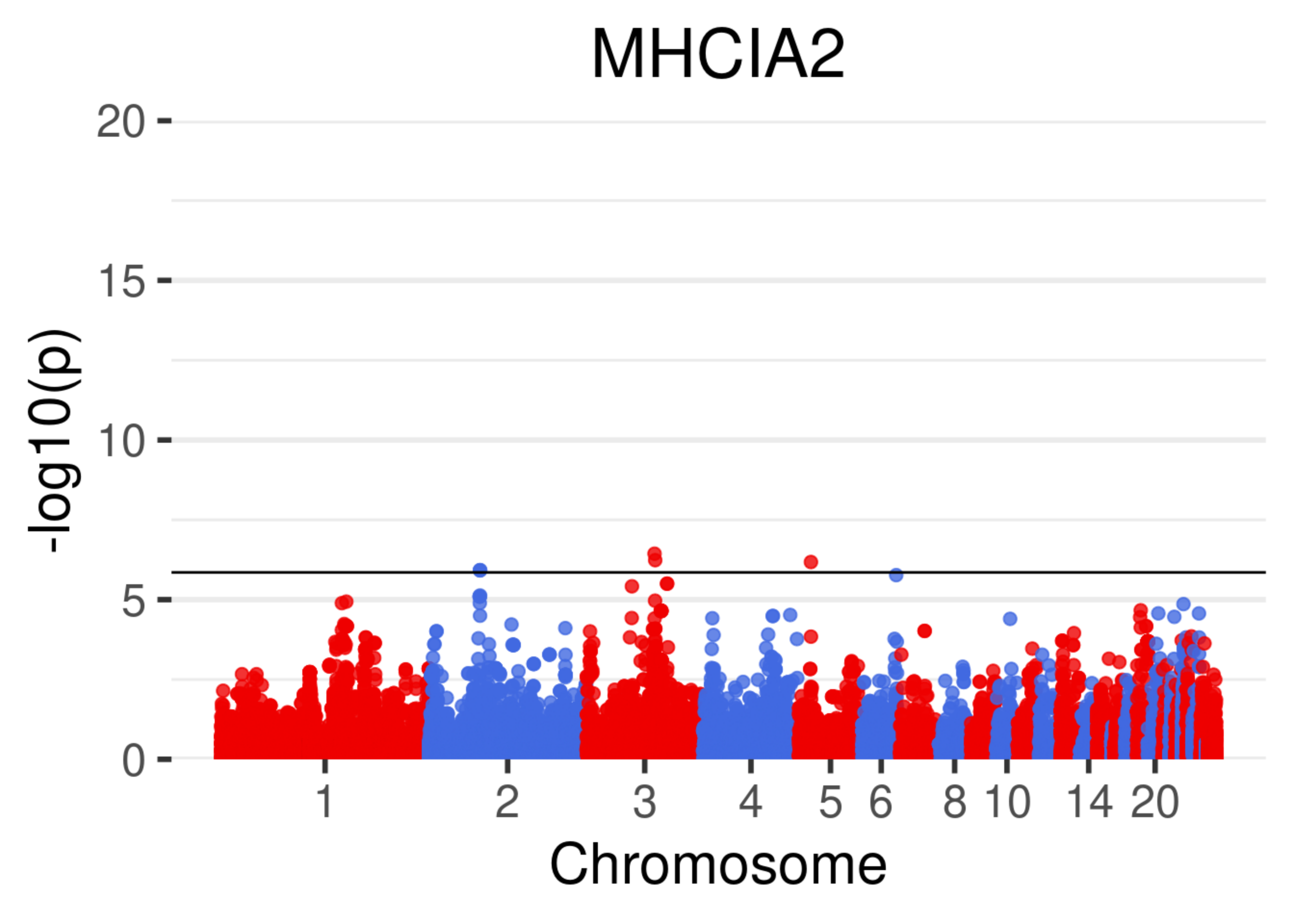
MHCIA2
-log10(p)
20
15
10
5
0
1
2
3
4
5
6
8
10
14
20
Chromosome

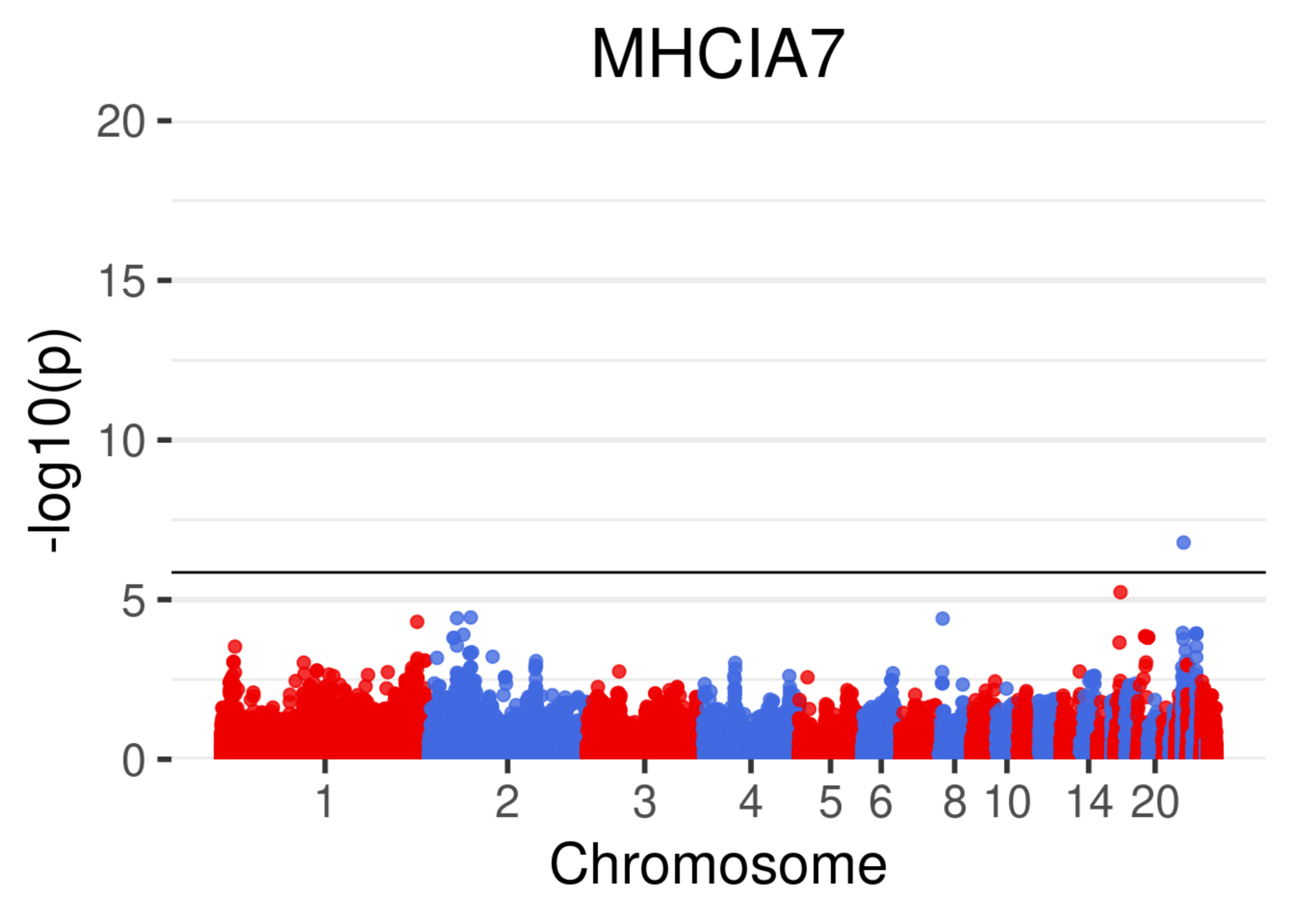
MHCIA7
-log10(p)
20
15
10
5
0
1
2
3
4
5
6
8
10
14
20
Chromosome

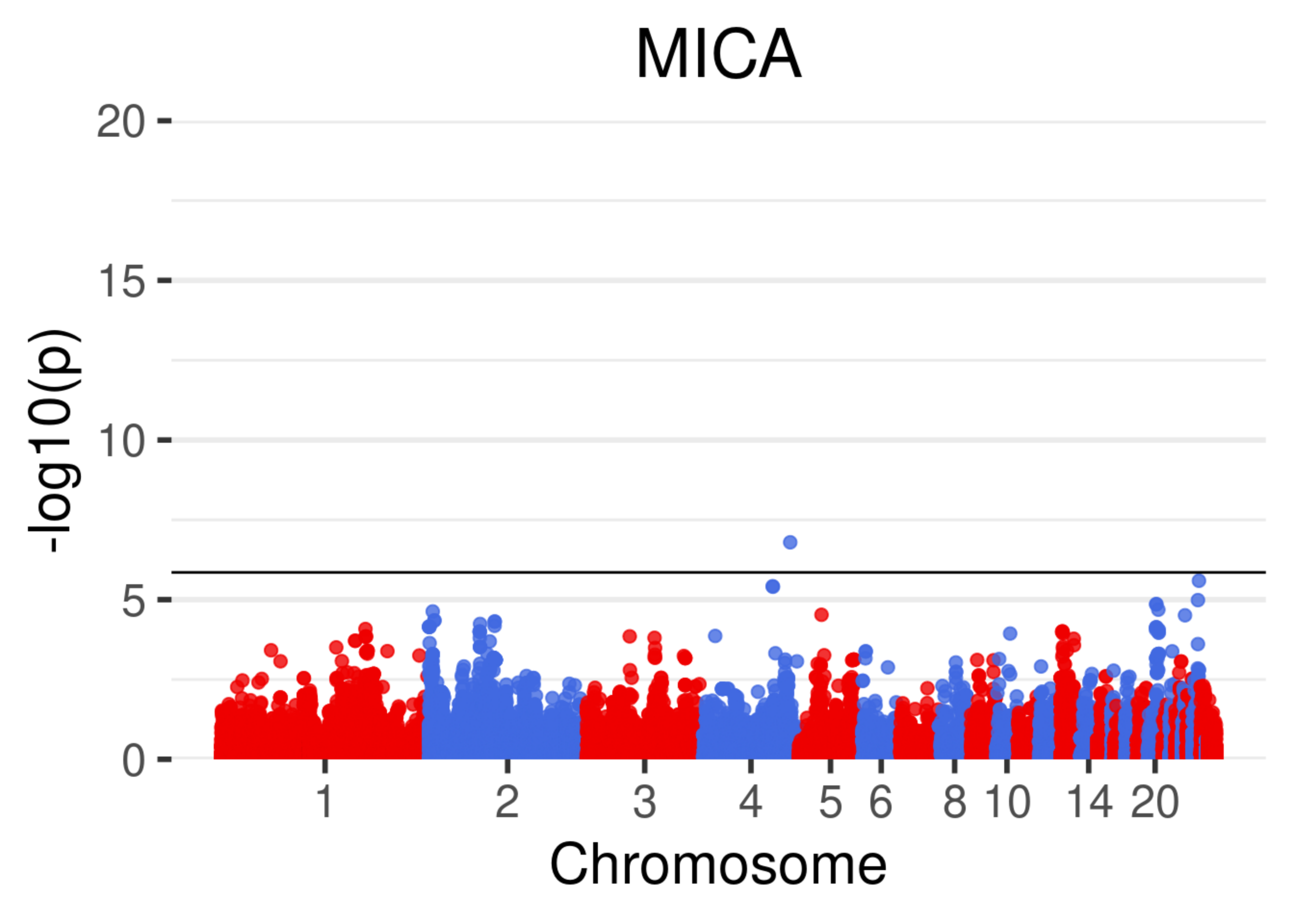
MICA
20
15
10
5
0
-log10(p)
1
2
3
4
5
6
8
10
14
20
Chromosome

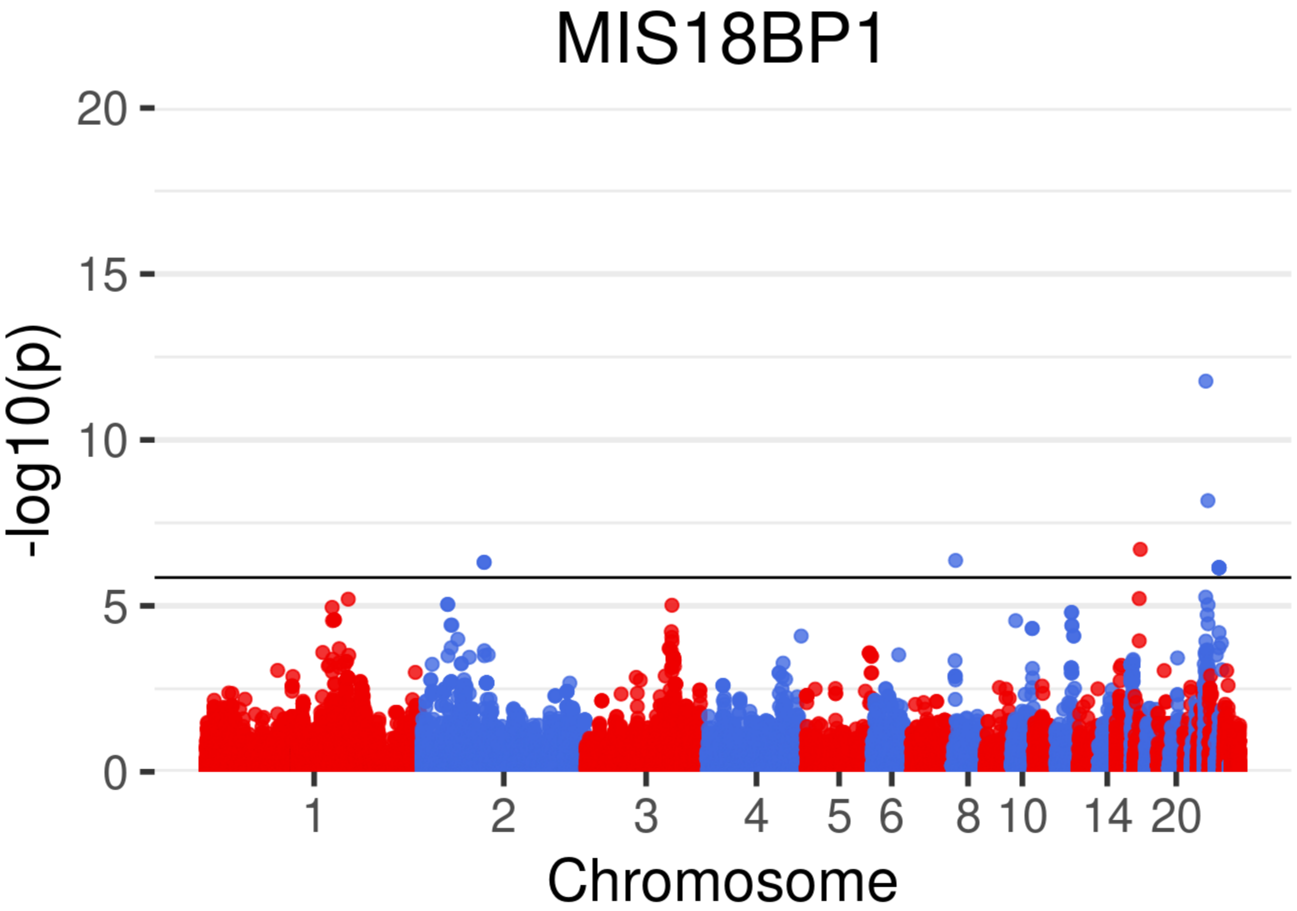

MIS18BP1
20
15
10
5
0
-log10(p)
1
2
3
4
5
6
8
10
14
20
Chromosome

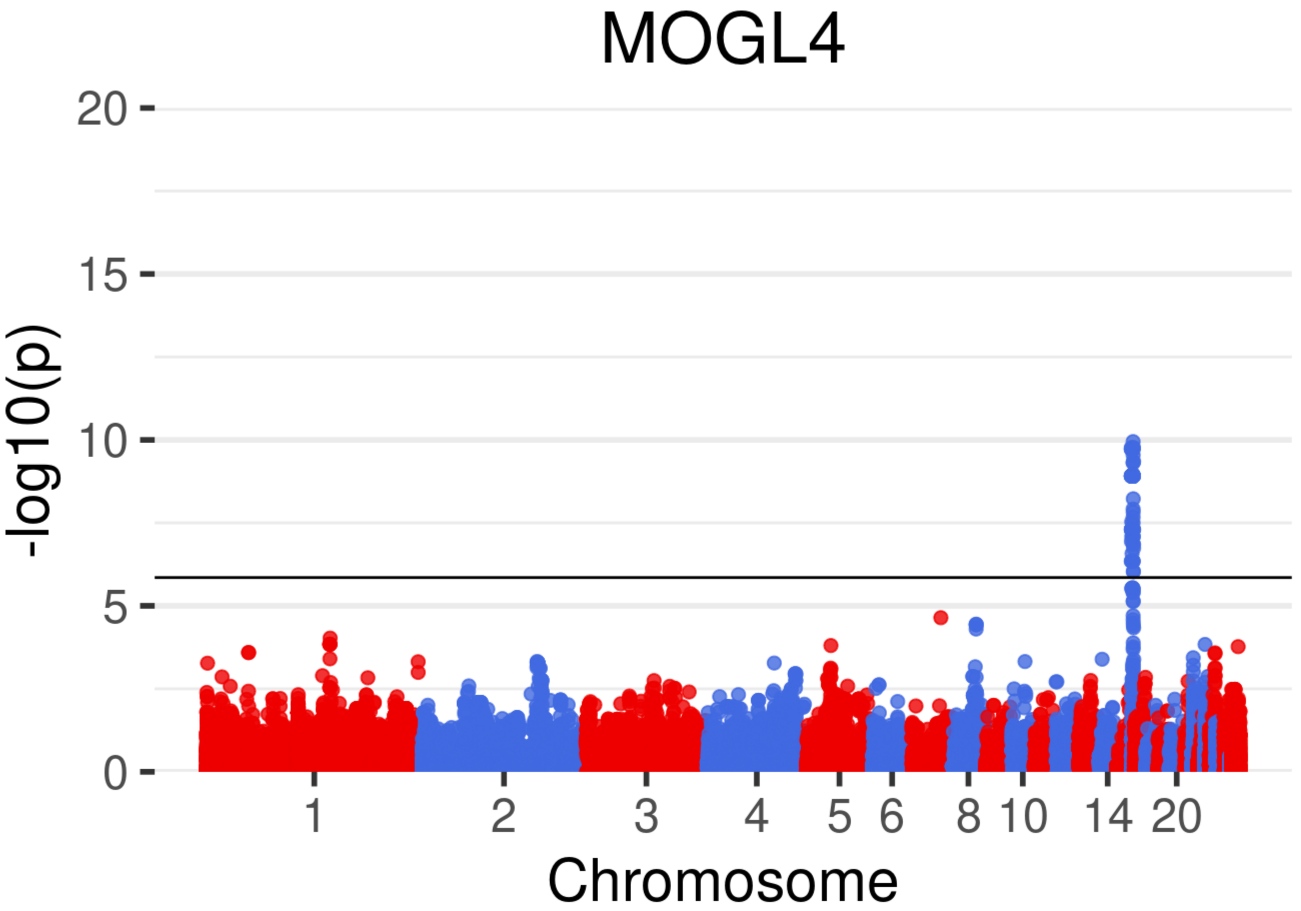
MOGL4
-log10(p)
20
15
10
5
0
1
2
3
4
5
6
8
10
14
20
Chromosome

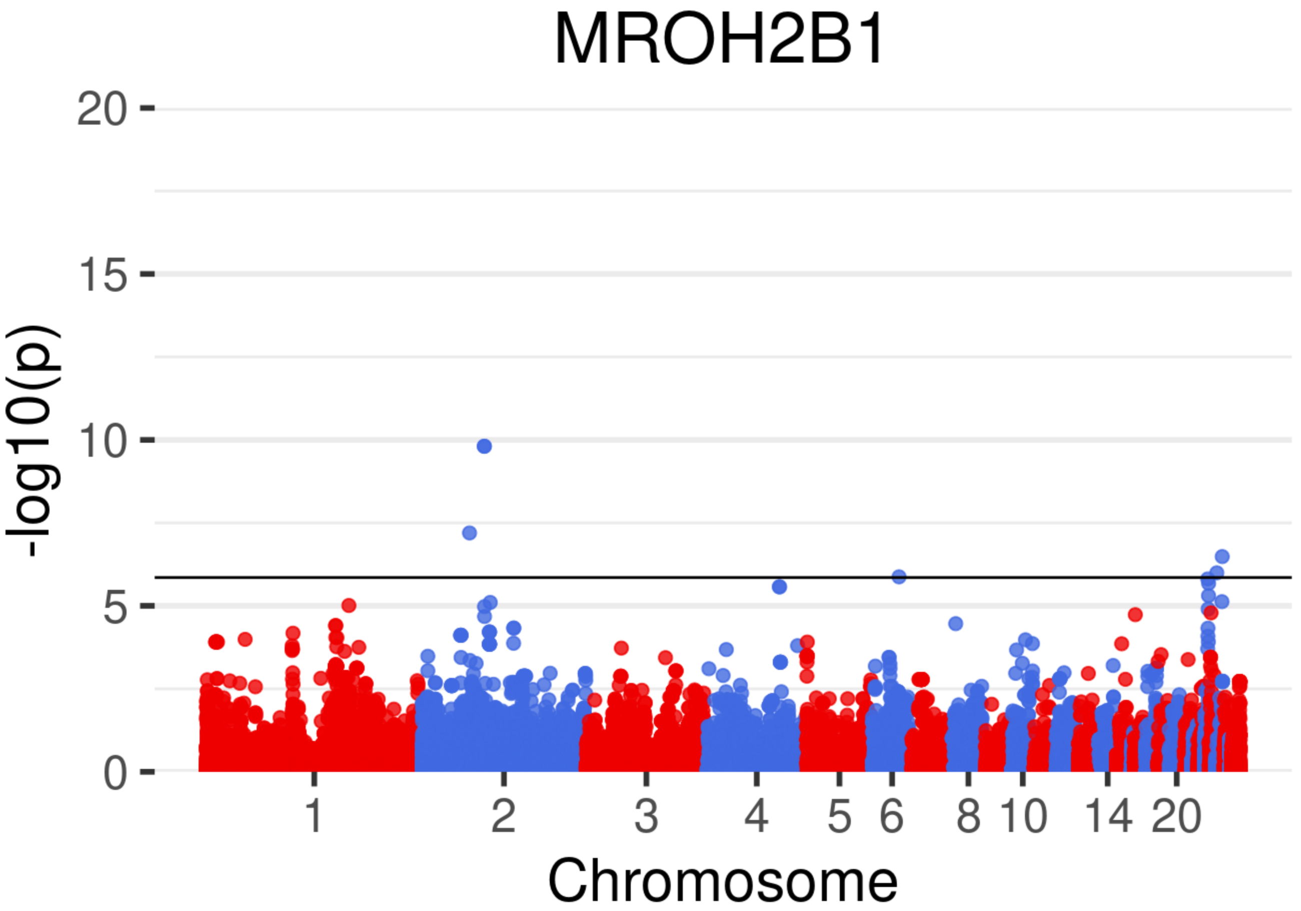

MROH2B1
-log10(p)
20
15
10
5
0
1
2
3
4
5
6
8
10
14
20
Chromosome

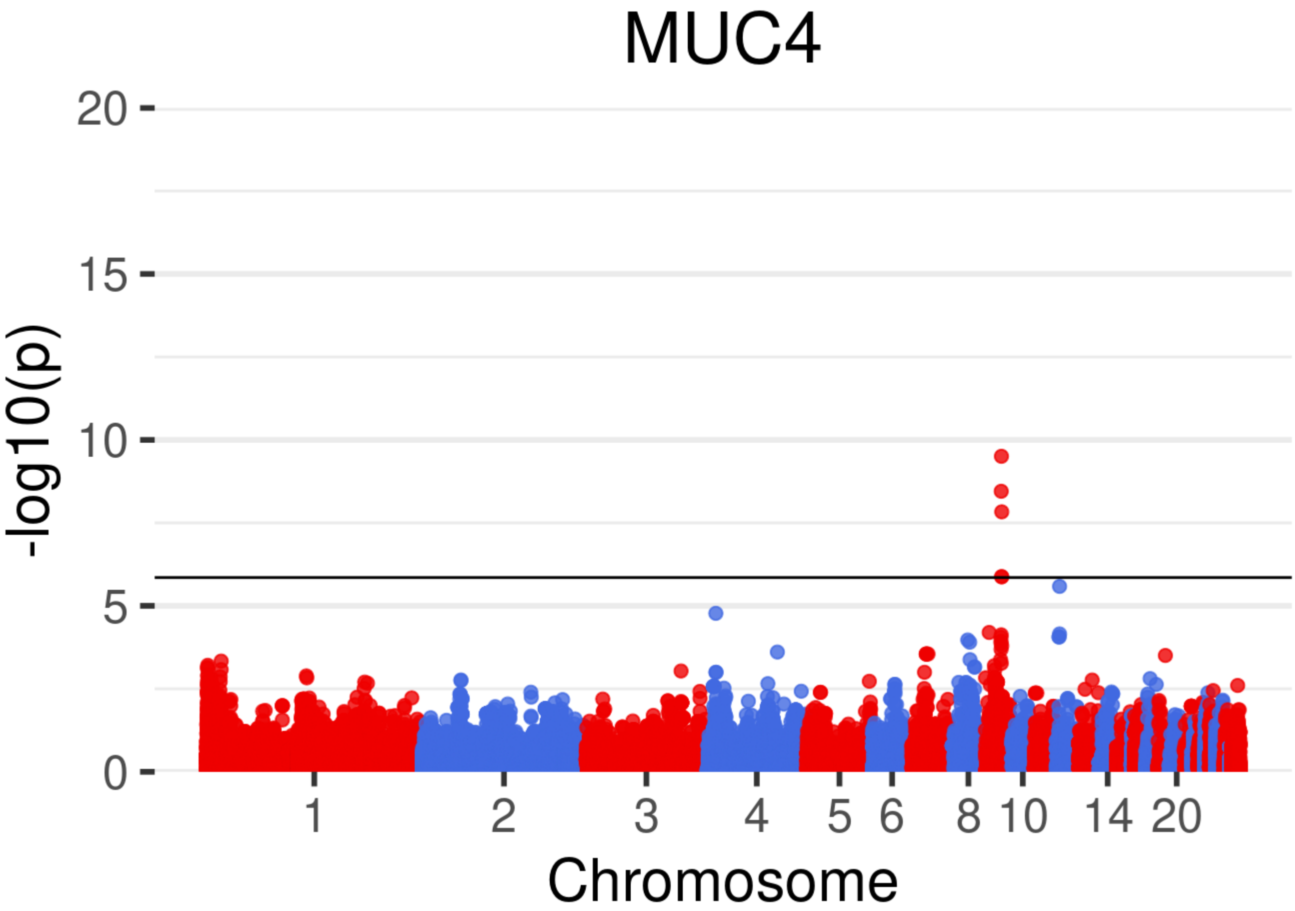
MUC4
-log10(p)
20
15
10
5
0
1
2
3
4
5
6
8
10
14
20
Chromosome

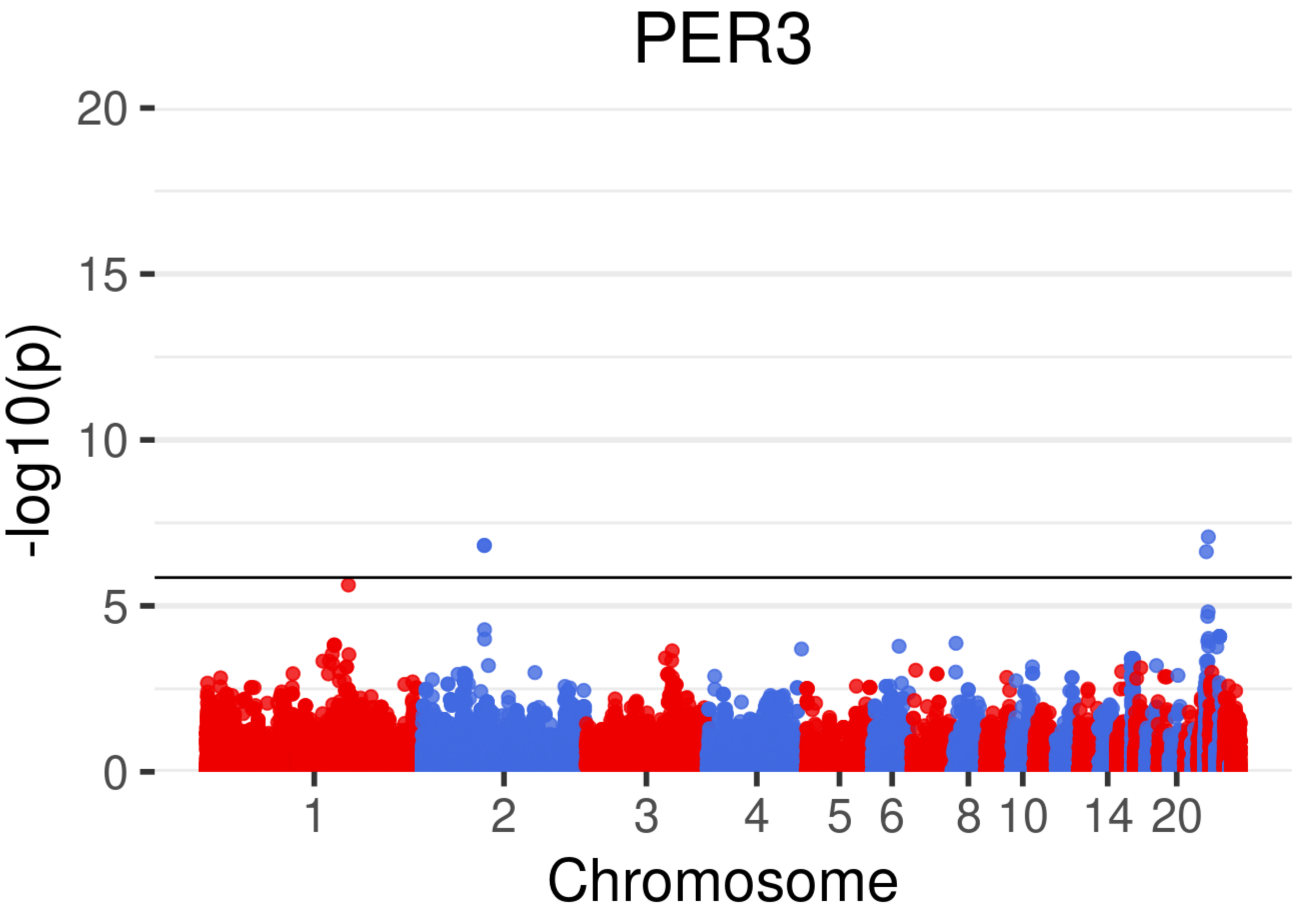
PER3
20
15
10
5
0
-log10(p)
1
2
3
4
5
6
8
10
14
20
Chromosome

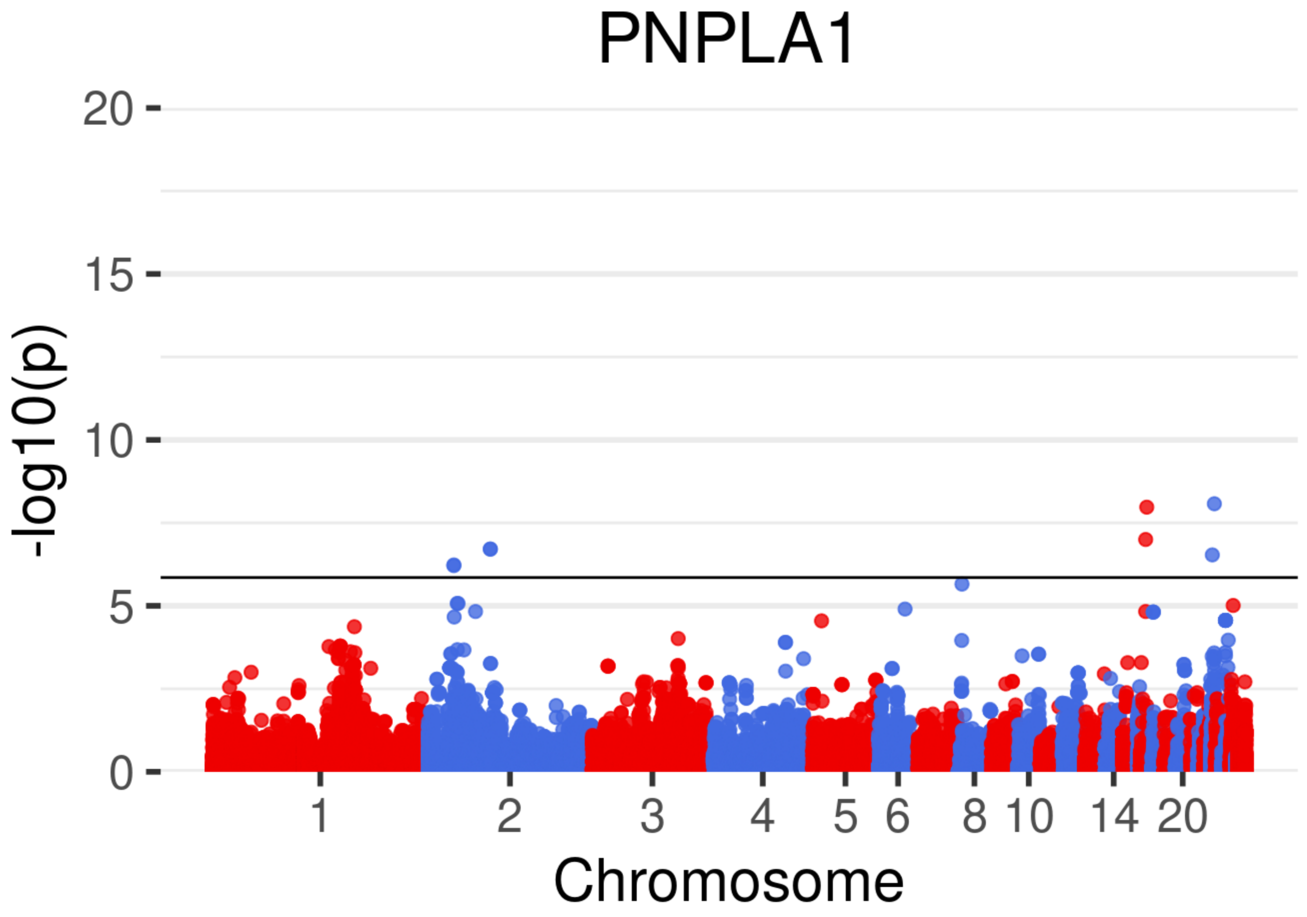

PNPLA1
-log10(p)
20
15
10
5
0
1
2
3
4
5
6
8
10
14
20
Chromosome

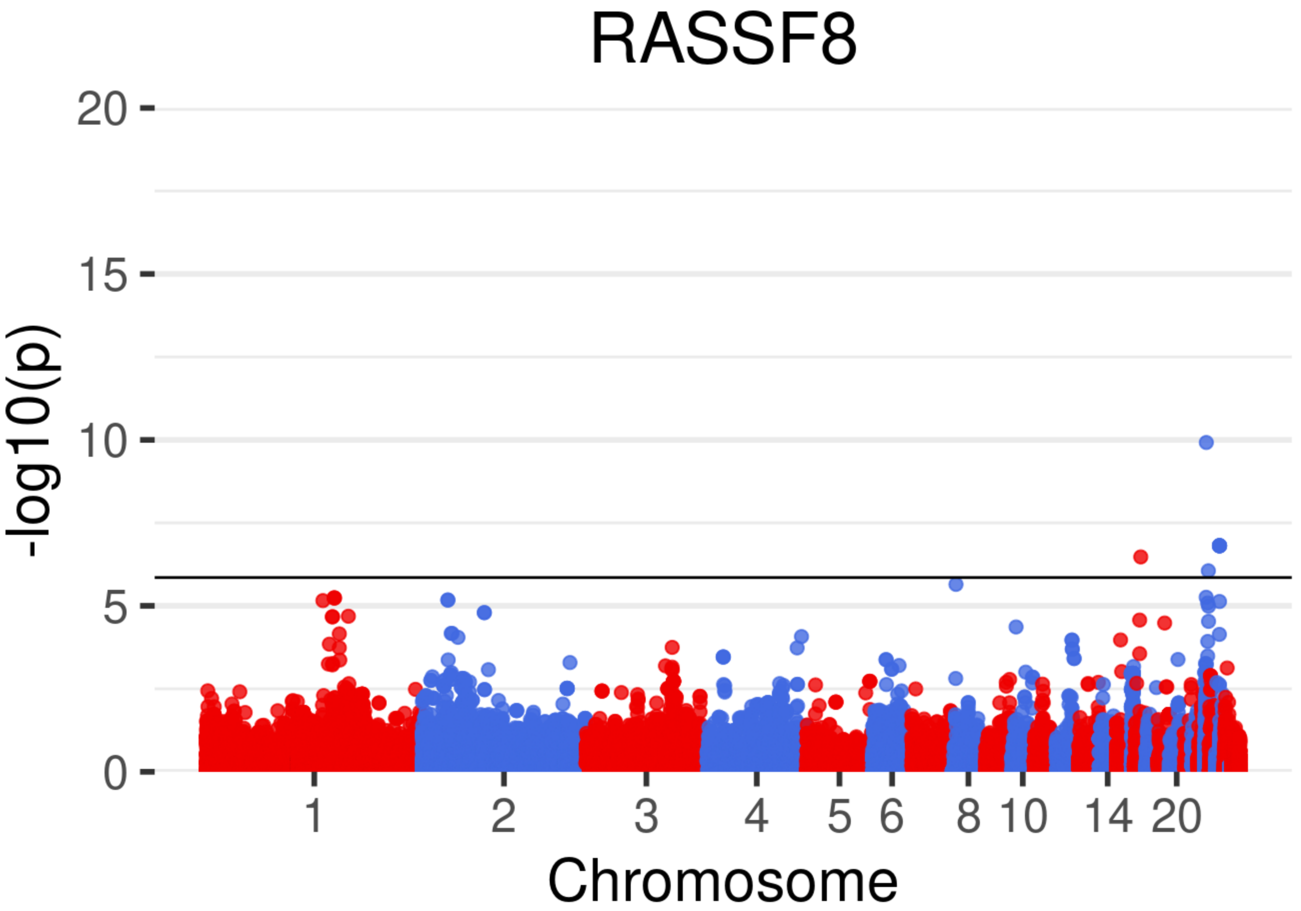

RASSF8
20
15
10
5
0
-log10(p)
1
2
3
4
5
6
8
10
14
20
Chromosome

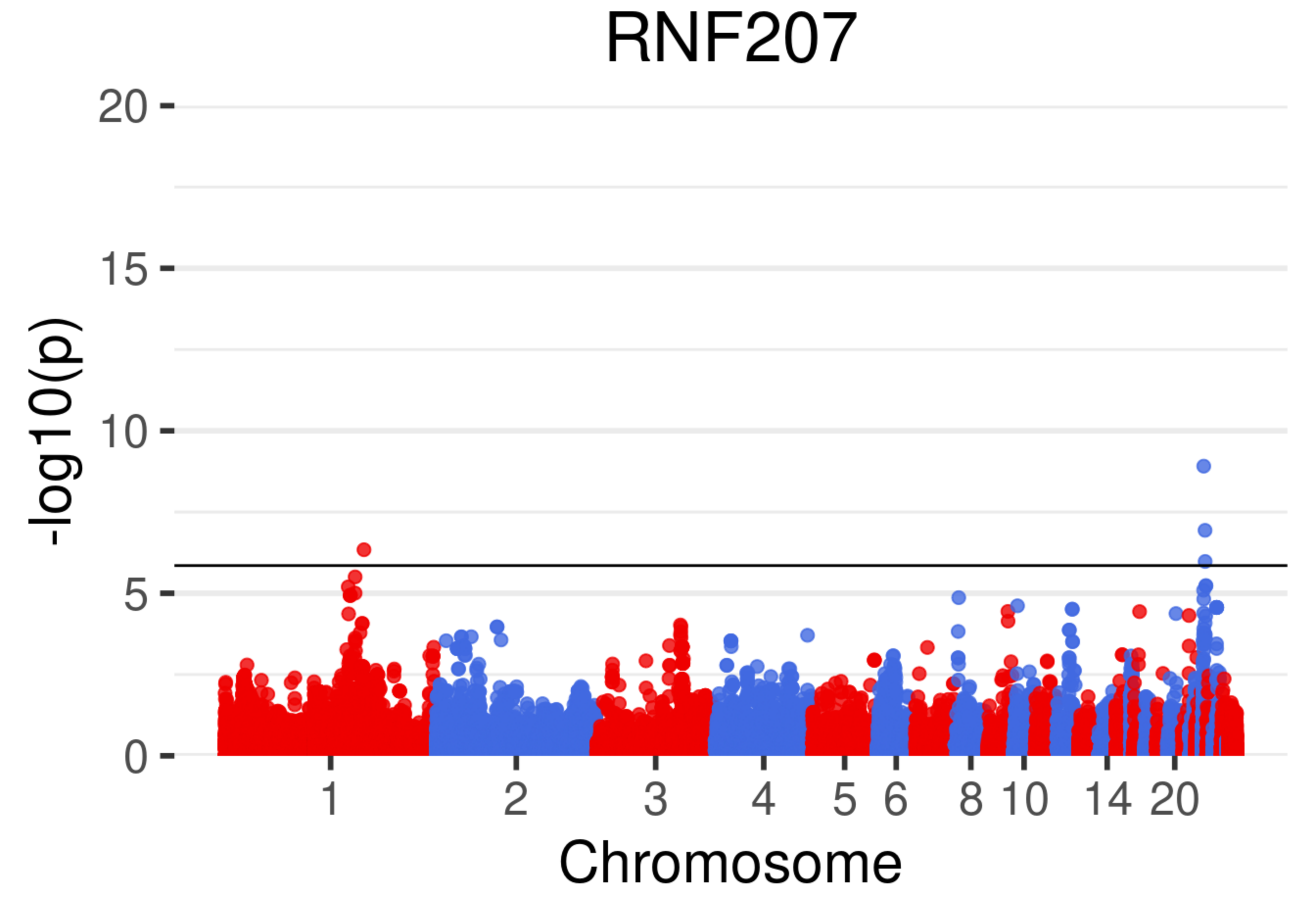
RNF207
-log10(p)
20
15
10
5
0
1
2
3
4
5
6
8
10
14
20
Chromosome

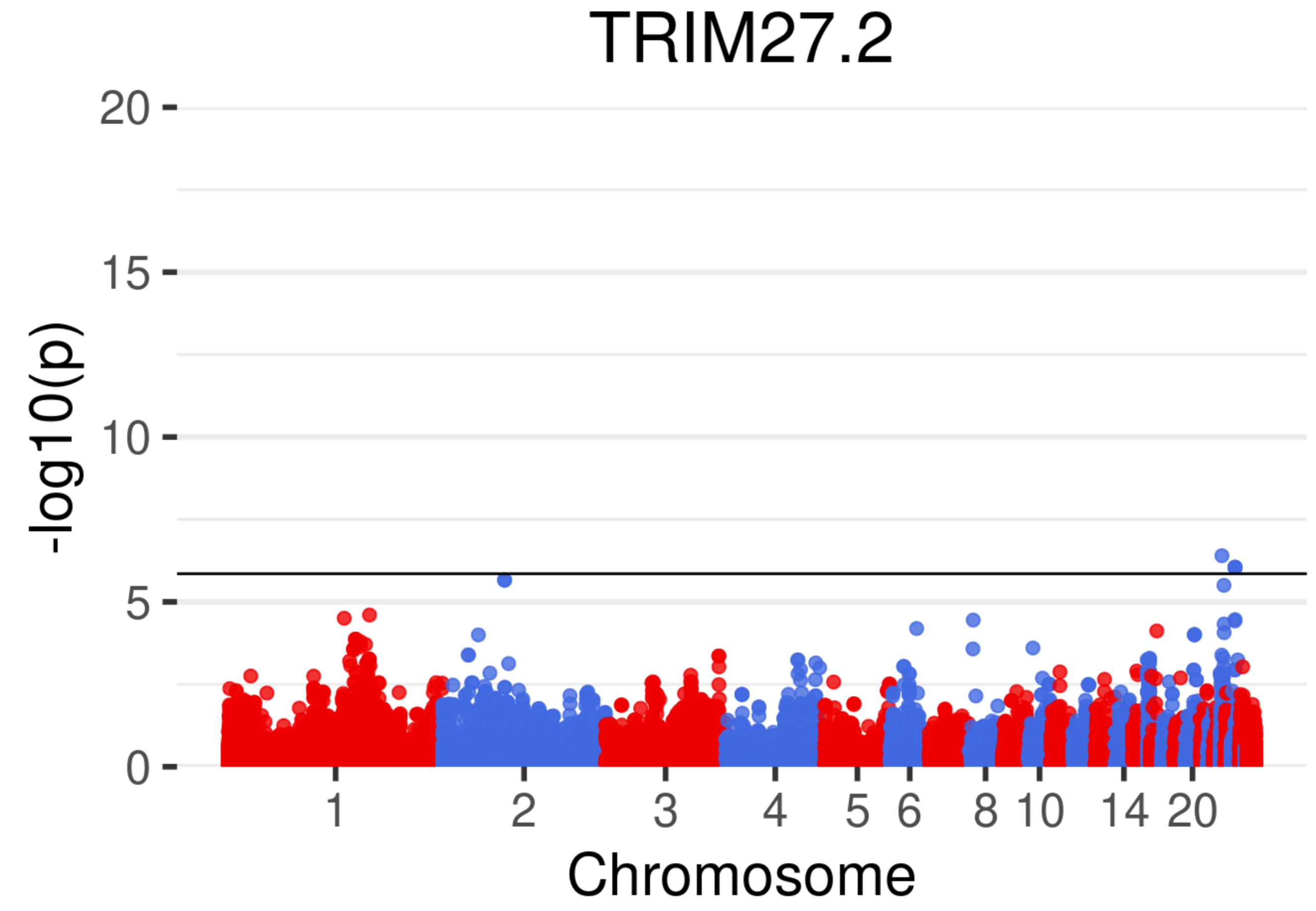

TRIM27.2
20
15
10
5
0
-log10(p)
1
2
3
4
5
6
8
10
14
20
Chromosome

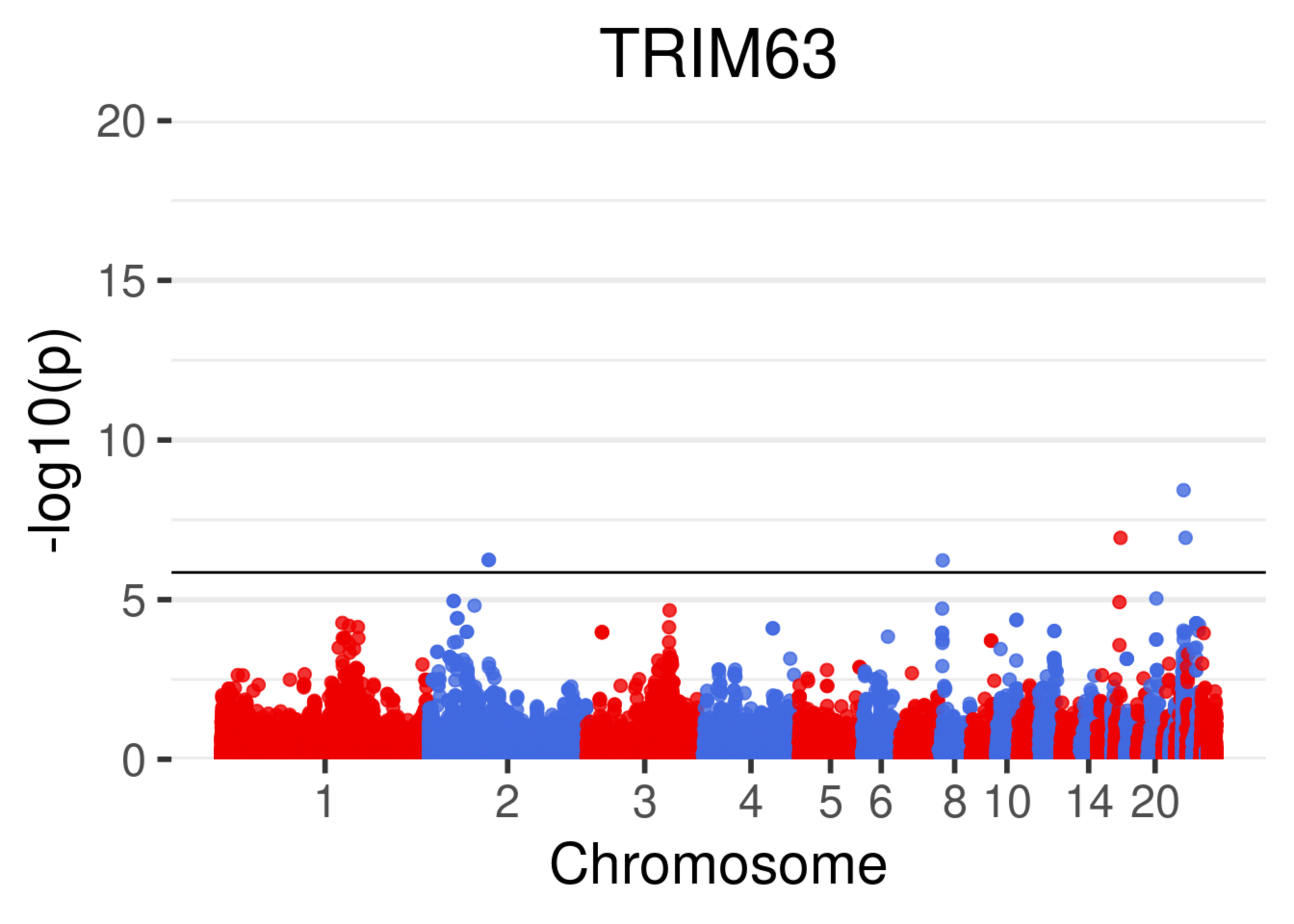

TRIM63
20
15
10
5
0
-log10(p)
1
2
3
4
5
6
8
10
14
20
Chromosome
