## Supplementary material for "Structural variation and eQTL analysis in two experimental populations of chickens divergently selected for feather pecking behavior": Graphical representation of ETV1 binding sites in proximity to associated differentially expressed genes

### Supplementary Information S3: Graphical representation of *ETV1* binding sites in proximity to associated differentially expressed genes.

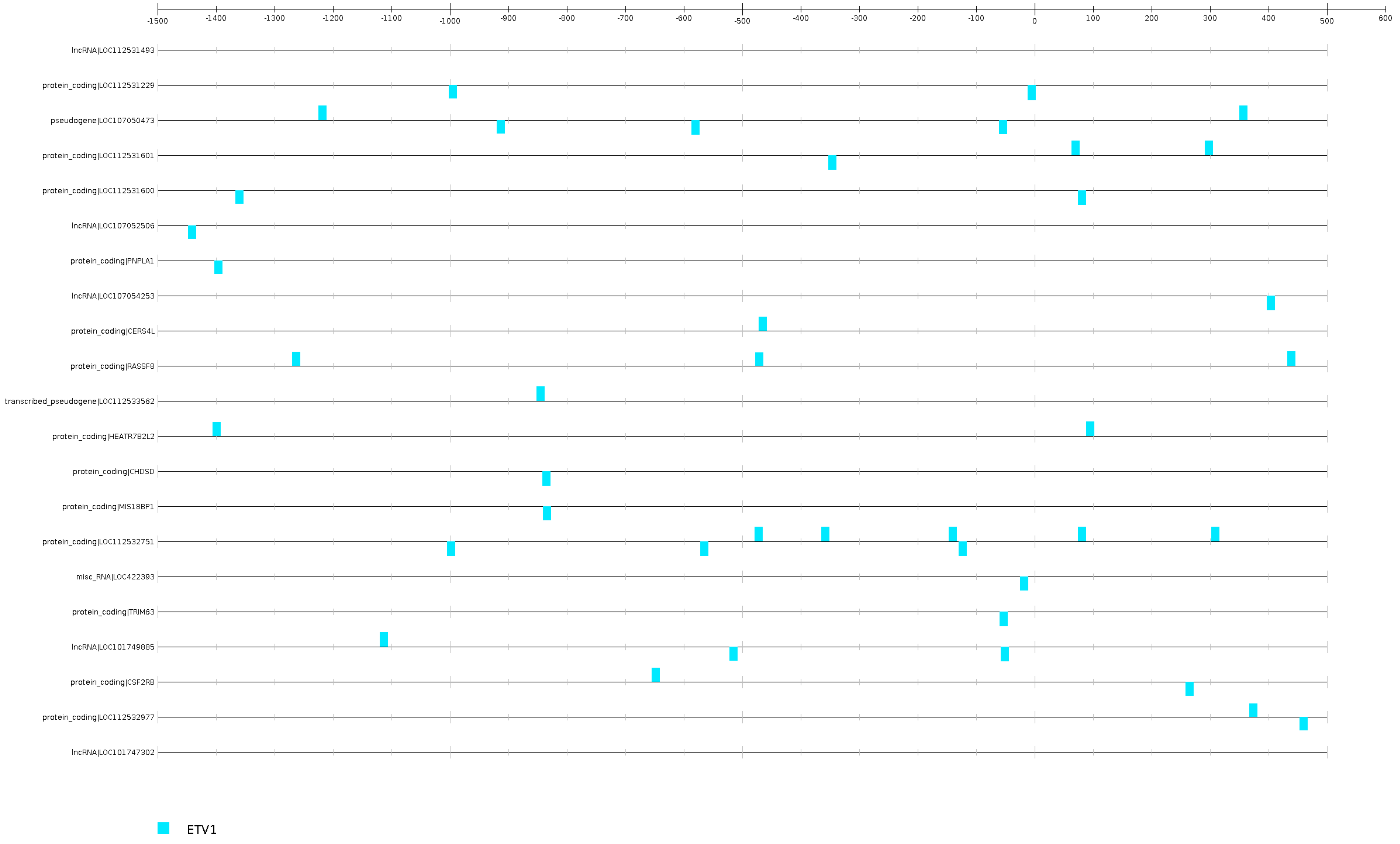
